# Asparagine accumulation in chicory storage roots is controlled by translocation and feedback regulation of asparagine biosynthesis in leaves

**DOI:** 10.1101/2020.04.27.063412

**Authors:** Emanoella Soares, Leonard Shumbe, Nicholas Dauchot, Christine Notté, Claire Prouin, Olivier Maudoux, Hervé Vanderschuren

## Abstract

- The presence of acrylamide (AA), a potentially carcinogenic and neurotoxic compound, in food has become a major concern for public health. AA in plant-derived food mainly arises from the reaction of the amino acid asparagine (Asn) and reducing sugars during processing of foodstuffs at high temperature.
- Using a selection of genotypes from the chicory germplasm we performed Asn measurements in storage roots and leaves to identify genotypes contrasting for Asn accumulation. We combined molecular analysis and grafting experiments to show that leaf to root translocation controls asparagine biosynthesis and accumulation in chicory storage roots.
- We could demonstrate that Asn accumulation in storage roots depends on Asn biosynthesis and transport from the leaf, and that a negative feedback loop by Asn on *CiASN1* expression impacts Asn biosynthesis in leaves.
- Our results provide a new model for asparagine biosynthesis in root crop species and highlight the importance of characterizing and manipulating asparagine transport to reduce AA content in processed plant-based foodstuffs.

## Introduction

Biosynthesis and accumulation asparagine (Asn) in crop species has gained attention since the discovery that Asn is a precursor in the formation of acrylamide (AA) during processing of raw materials at temperatures above 120 °C. In the liver, AA is metabolized to glyciamide, an epoxide that is more reactive to DNA and proteins than AA itself (Pedreschi *et al.*, 2014). Although AA is only classified as probably carcinogenic to humans (Group 2A), a recent study identified a mutational signature in human tumor tissues that appears imprinted by AA, through the effects of glycidamide (Zhivagui *et al.*, 2019). AA is formed through the Maillard reaction during the process of baking, roasting, frying, toasting and microwaving (Stadler *et al.*, 2002). One of the major concerns about the presence of AA in foods is its presence in many foodstuffs consumed on a daily basis, such as: French fries, potato crisps, bread, fine bakery wares, coffee, coffee substitutes and baby food (Commission Regulation (EU) N2017/2158). In an attempt to prevent and reduce formation of AA during the industrial processing of foodstuffs, an Acrylamide Toolbox was created by the Food Drink Europe (‘Food drink Europe. Acrylamide Toolbox’, 2019). This Toolbox is based on four parameters: (1) agronomy, (2) recipe, (3) processing, and (4) final preparation. Because the strategies 2 and 3 often alter the properties (i.e. flavour, appearance, quality) of the processed product and are food system specific, exploitation of the natural variation in AA precursors present in plants appears as a promising route. Therefore, investigation of biosynthesis and accumulation of AA precursors in plant tissues/organs submitted to processing, has become key to help reducing exposure to AA. The identification of genetic factors involved in the production, transport and accumulation of AA precursors is particularly relevant to address this issue globally with improved varieties with low potential for AA production.

Asn is the most important determinant for AA formation as its concentration in raw materials has been shown to be directly correlated with AA formation in potato (Zyzak *et al.*, 2003), coffee (Bagdonaite *et al.*, 2008) wheat (Muttucumaru *et al.*, 2008), rye (Curtis *et al.*, 2010) and chicory (Loaëc *et al.*, 2014). Strategies based on agricultural practices, natural variation and biotechnology have recently been investigated to reduce the AA potential of crop species (Loaëc et *al.*, 2014; Zhu et al., 2016; Curtis et al., 2018). Asn, besides being a proteinogenic amino acid, plays a central role in nitrogen metabolism in plants, acting in nitrogen transport, recycling and remobilization during development (Gaufichon *et al.*, 2010). The main Asn biosynthetic pathway in plants involves the assimilation of ammonium into Asn through a series of reactions with the participation of four enzymes: glutamine synthetase (GS), glutamate synthetase (GOGAT), aspartate aminotransferase (AspAT) and asparagine synthetase (ASN), all belonging to multigene families (Gaufichon *et al.*, 2010; Gaufichon *et al.*, 2015). Because ASN catalyses the final step of Asn biosynthesis, genetic engineering approaches to produce crop plants with low Asn content have relied on the organ-specific alteration of *ASN* expression (Rommens *et al.*, 2008; Chawla *et al.*, 2012). However, alteration of Asn biosynthesis can also lead to phenotypic variation and yield penalty (Rommens *et al.*, 2008). This reflects the importance of Asn homeostasis for the plant metabolism. It has also been demonstrated that over-expression of the protein kinase general control nonderepressible (GCN)‐2, a protein that phosphorylates the α subunit of eukaryotic translation initiation factor 2 (eIF2α) could lead to a significant decrease of total free amino acids in wheat grains and a concomitant reduction in *ASN1* expression (Byrne *et al.*, 2012).

Although Asn is associated with N remobilization *in planta* through transport in vascular tissues (Krapp, 2015; Tegeder & Hammes, 2018), the contribution of long-distance transport to Asn accumulation in crop plants has so far received limited attention. Chicory (*Cichorium intybus* L.) belongs to the Asteraceae family and is used as a root crop for the production of inulins as well as a coffee substitute. The latter requires processing at high temperature and is prone to AA production. Previous studies have demonstrated that the AA content in processed chicory roots is directly correlated with the Asn concentration (Loaëc *et al.*, 2014). Recent advances in genome sequencing and the availability of genotypes from breeding programmes aiming at reducing their AA potential make chicory a species of interest to investigate Asn biosynthesis and transport. The production of storage root during chicory development also offers the opportunity to investigate the role of sink organs in the Asn metabolism.

Here, we investigated accumulation and transport of free Asn between chicory organs and their impact on Asn biosynthesis. For this purpose, we identified the Asn biosynthetic genes in the chicory genome and characterized genotypes for their Asn biosynthesis and accumulation in leaves and roots during plant development. We took advantage of grafting experiments and Asn measurement in genotypes contrasting for Asn biosynthesis and accumulation to identify a negative feedback regulation by Asn. We propose a model accounting for Asn biosynthesis and transport between organs. Our study highlights the importance of studying Asn transport in crop plants with high AA potential and opens new perspectives to modify Asn accumulation without altering its biosynthesis.

## Materials and Methods

### Contrasting genotypes for Asn content in chicory germplasm

For this purpose a field trial (year 1) was conducted at Cosucra Group Warcoing SA (Pottes, Belgium) from May to November of 2017 with 18 genotypes of *Cichorium intybus* including S_3_ to S_8_ lines (L3427, L4108, L4115, L4118, L8014, L5040), inbred lines (inb14, inb40, inb42, VL54, VL57, VL59), hybrids (HYB50, OBOE), synthetic varieties (Larigot, Cadence, Hera) and a variety with uncharacterized genetic structure (Malachite). Storage roots of each genotype were harvested at 30, 60, 90, 120, 150 and 180 das for analysis of the Asn content. The experimental design consisted of 5 randomized blocks where 4 - 8 storage roots, in each block, were harvested and pooled for the analysis. The year 2 experiment was conducted with the genotypes L4115, L4118, inb42, L8014, HYB50, OBOE and Larigot that were harvested at 60, 120 and 180 das. For the second experiment (year 2), the experimental design consisted of 5 randomized blocks where 6 storage roots, in each block, were harvested and pooled for analysis.

### Asparagine measurement

The determination of free Asn content in storage roots was carried out with an L-asparagine kit (Megazyme, K-ASNAM). Measurements were carried out according to(Lecart *et al.*, 2018) with some modifications. Briefly, 200 mg of freeze-dried and ground roots were mixed with 1.6 mL of 1 M perchloric acid and homogenized in a TissueLyser II (Qiagen) in microtubes containing 6 glass beads of 2.8 mm in diameter. Samples were centrifuged at 10000 x g for 20 minutes and 1 mL of the supernatant was collected and neutralized with 1 M KOH until reaching pH 8.0 (~ 500 µL). The material was then placed on ice for 20 minutes prior to centrifugation at 10000 x g for 10 minutes. The supernatant was immediately used for Asn measurement according to the manufacturer’s instructions for the microplate format. Each sample was quantified in triplicate.

### Identification of Asn biosynthetic genes

To identify the genes involved in the Asn biosynthetic pathway in chicory we took the advantage of restricted access to the chicory genome and a chicory EST database (Nicolas Dauchot, University of Namur, personal communication). A BLAST approach was performed using orthologous sequences of asparagine synthetase, glutamine synthase, glutamate synthase, and aspartate aminotransferase retrieved from Arabidopsis (https://www.arabidopsis.org/) as well as Asteraceae family species (*Lactuca sativa* and *Helianthus annuus*) and Asterid clade species (*Solanum tuberosum* and *Solanum lycopersicum*) retrieved from NCBI. All chicory sequences are available in Notes S1.

### RNA extraction and RT-qPCR analysis

Total RNA was purified from freeze-dried grinded leaves and roots. Up to 50 mg of plant material was lysed with buffer RLT (Qiagen) followed by protein removal with chloroform:isoamyl alcohol (24:1) and RNA precipitation with isopropanol. The resulting RNA was washed with 70% ethanol and resuspended in RNase free water. Quantification was performed using the Quantus Fluorometer (Promega) and RNA quality was checked by agarose gel electrophoresis. Subsequently, 2 µg of RNA for each sample was treated with DNase I at 37°C for 10 minutes (New England Biolabs) and 0.5 µg was used for reverse transcription with the GoScript Reverse Transcription Mix, Oligo dT (Promega). The concentrations of the resulting cDNAs were adjusted to 10 ng/µL and 2 µL were used for RT-qPCR reactions performed with the GoTaq qPCR Master Mix (Promega) in a final volume of 10 µL. Amplification was detected using the CFX96 Touch Real-Time PCR Detection System (Bio-Rad). To further identify stable reference genes to our experimental conditions, viz., storage roots, leaves and contrasting genotypes to Asn, a normalization experiment was conducted with reference genes available in the literature for chicory. Primer sequences for actin (*ACT*); elongation factor 1-alpha (*EF*); histone H3 (*H3*) and 18S rRNA (*rRNA*) were retrieved from(Maroufi *et al.*, 2010) while for protein phosphatase 2A subunit A2 (*PP2AA2*); protein phosphatase 2A subunit A3 (*PP2AA3*); SAND family protein (*SAND*); TIP41 like protein (*TIP41*) and ubiquitin-conjugating enzyme (*UBC*) were retrieved from(Delporte *et al.*, 2015). A score was attributed to each reference gene according to their stability in the softwares geNorm(Vandesompele *et al.*, 2002), NormFinder(Andersen *et al.*, 2004), and BestKeeper(Pfaffl *et al.*, 2004) and a ranking from the most stable genes (lowest score) to the most unstable (highest score) was performed (see Figs. S11 and S12). The optimal number of reference genes for storage roots and leaves was chosen according to the pairwise variation V from geNorm (Fig. S11) and the selected genes were chosen based on the highest stability according to the ranking (Fig. S12).

### Grafting

The experiment was conducted under greenhouse conditions. A protocol for chicory grafting was established as depicted in Fig. S8. Seeds were germinated in the dark for 5 days to promote stem growth. The seedlings were subsequently transferred to a 16/8 h photoperiod for 1 month before grafting. Cotyledons and leaves were removed and plantlets used for grafting by the cleft method. The cuttings for the grafting were performed with a razor blade under a binocular stereoscopic microscope. The junctions were covered with a porous tape (Micropore ^TM^ – 3M) and placed high above the substrate in 3 L pots. A humid environment was created by covering the pots with transparent bags that were removed after 14 days. One hundred mg of NH_4_NO_3_ was added to each pot one month after grafting. Plants were harvested after 3 months; leaves and storage roots were collected, immediately frozen in liquid nitrogen, and freeze-dried for use in biochemical and molecular analyses. Analyses were performed on three to eight biological replicates for each grafting combination.

### Leaf infiltration with amino acids

45 day-old *in vitro* chicory plantlets grown at 25 °C under a photoperiod of 16/8 hours were used for leaf infiltration of amino acids. Asparagine (190 mM), aspartate (37 mM), glutamate (57 mM) and glutamine (190 mM), individually solubilized in water, were infiltrated with a 6 mL syringe (NORM-JECT) into chicory leaves. These concentrations were chosen according to the maximum solubility in water of each amino acid. After injection, plants were kept in the growth chamber for 36 hours. After this period, leaves were harvested to proceed with gene expression analysis. Each treatment was conducted in 4 biological replicates.

### Protoplasts feeding with Asn

Protoplasts from leaves were isolated according to Deryckere *et al*.(Deryckere *et al.*, 2012). The protoplasts were resuspended in WI solution (4 mM MES (pH 5.7) containing 0.5 M mannitol and 20 mM KCl) and a total of 8 × 10^6^ cells were placed in 24-well plates. Asn was added to a final concentration of 190 mM and a final volume of 250 µL for protoplast feeding assay. The protoplast plates were incubated at room temperature for 36 hours. Protoplasts were subsequently used for RNA extraction and gene expression analysis.

## Results

### Diversity of Asn content in chicory germplasm

We first identified genotypes contrasting for Asn content in storage roots to compare the specificities of low and high-Asn chicory genotypes at the molecular level. The germplasm bank included lines, hybrids and synthetic varieties. Our analysis revealed genotypes with Asn content in storage roots ranging from 27.3 mg/100g + 5.2 mg/100g (L4115) to 209.8 mg/100g + 35.8mg/100g (L4118) dry weight (DW) 150 days after sowing (das) prior to the initiation of senescence (Fig. 1a). A classification was also performed with the Asn content at 180 das (Fig. S1). Based on the Asn content at 150 das, the genotypes could be divided into four groups according to their Asn levels in storage roots: low Asn content (LAsn), medium Asn content (MAsn), medium-high Asn content (MHAsn) and high Asn content (HAsn). The Asn content in storage roots did not correlate with root diameter (r = −0.03) and root weight (r = −0.05) (Fig. S2) indicating that the Asn levels could not be explained by the phenotypic variation at harvest (Fig S3). Noticeably the genotype classified as LAsn displayed stable Asn levels from 30 das until harvest (Fig. S4). On the contrary, Asn levels increased over time in genotypes classified as MAsn, MHAsn and HAsn (Fig. S4). Several genotypes accumulated increasing levels of Asn in storage roots until harvest while Asn levels appeared to peak and reach a plateau before harvest in other genotypes (Fig. S4). A second year field experiment was conducted to confirm Asn accumulation profiles from selected genotypes. The selected genotypes displayed similar trends in Asn accumulation despite lower levels observed in the second year field trial (Fig. S5), possibly due to environmental variation as previously reported (Loaëc *et al.*, 2014). In particular differences in N availability could have occurred due to reduced precipitation and dryer conditions during the second year.

**Fig. 1.**
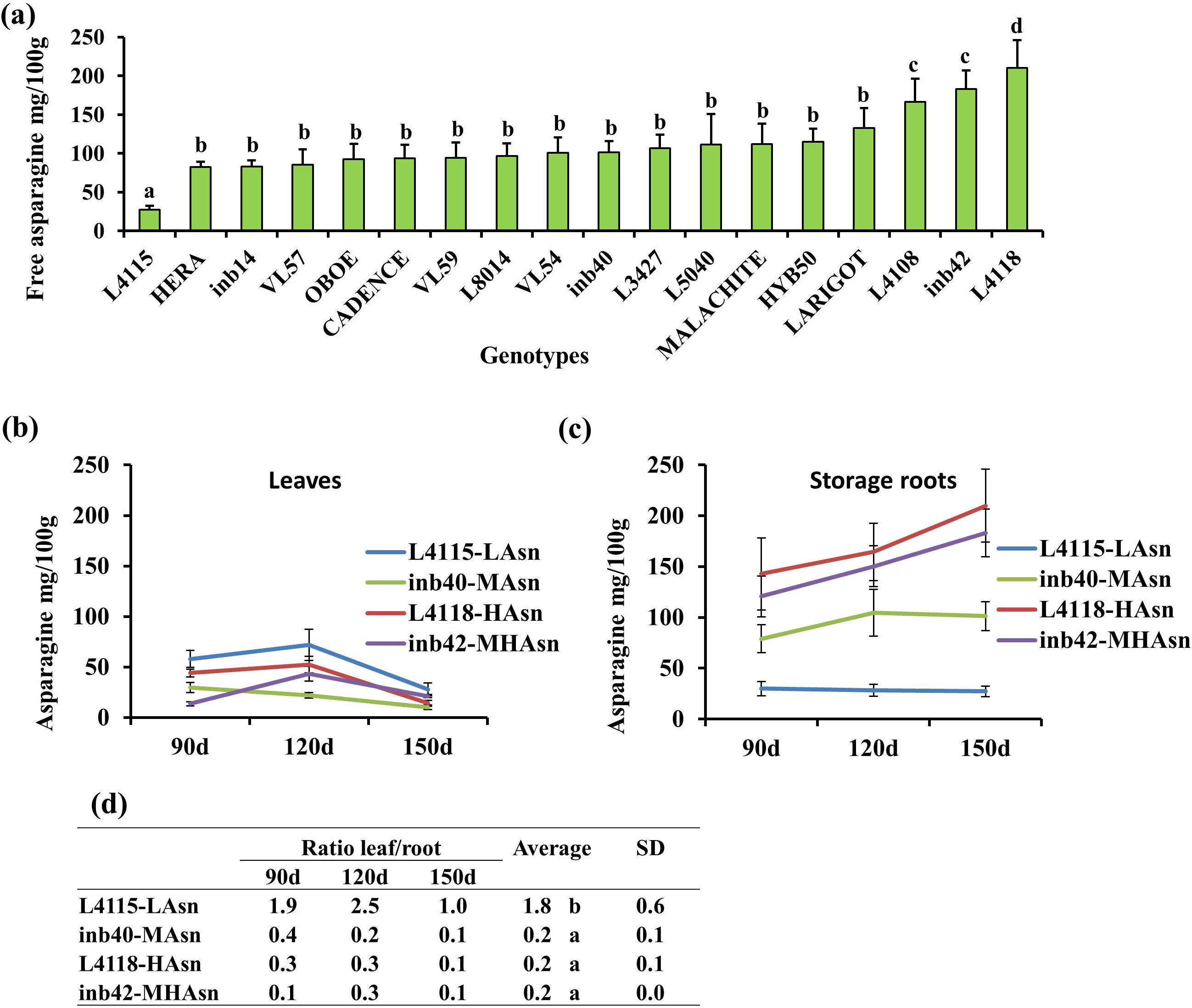
Diversity of Asn accumulation in the chicory germplasm. (a) Free Asn content (mg/100 g dry matter) in storage roots of selected chicory genotypes 150 days after sowing. Data are mean + s.d. of 5 biological replicates containing 4 or 8 storage roots (n=20 or n=40). Statistically significant differences are represented by different letters (one-way ANOVA, *P* < 0.05 followed by Scott-Knott test, *P* < 0.05). Genotypes with the letters a, b, c, d were classified as low Asn content (LAsn), medium Asn content (MAsn), medium-high Asn content (MHAsn) and high Asn content (HAsn), repectively. (b), (c) Free Asn accumulation rate (mg/100 g dry matter) in leaves (b) and storage roots (c) from selected contrasting genotypes at 90, 120 and 150 days after sowing. Each point represents the mean + s.d. of 5 biological replicates. For more details of other genotypes see Fig. S2. (d) Ratio of free Asn content between leaves and storage roots of selected contrasting genotypes at 90, 120 and 150 days after sowing.

We next measured the Asn content in leaves from the genotypes contrasting for Asn accumulation in storage roots (L4115-LAsn, inb40-MAsn, L4118-HAsn, inb42-MHAsn) to investigate the relation between Asn contents in storage roots and leaves (figs. 1b and 1c). By computing the [Asn]_leaf_/[Asn]_storage root_ ratio, we found that the genotype L4115-LAsn had a significantly higher ratio (1.8) as compared to the other genotypes (0.2) (Fig. 1d). Moreover, we found a negative correlation (r = −0.41) between [Asn]_leaf_ and [Asn]_storage root_ in those genotypes (Fig. S6). From those observations, we hypothesized that chicory storage roots could act as a sink for Asn whose strength varies across genotypes, inb40-MAsn, inb42-HAsn and L4118-HAsn harboring the strongest root sinks. Higher accumulation of Asn in strong sinks (spiklets) compared to source (flag leaves) has previously been reported in rice(Yabuki *et al.*, 2017).

### Asparagine biosynthetic pathway in chicory

Having identified chicory genotypes displaying a variation in both [Asn]_leaf_ and [Asn]_storage root_, we next identified Asn biosynthetic genes in the chicory genome. Chicory genomic sequences were BLAST searched using Asn biosynthetic genes orthologs from Arabidopsis in order to identify gene members of each family involved in the biosynthesis of Asn in chicory. We identified two genes coding for AS (*CiASN1* and *CiASN2*), four genes coding for GS (*CiGLN1;1, CiGLN1;2, CiGLN1;3, CiGLN2*), two genes coding for GOGAT(*CiGLU1, CiGLT*) and four genes coding for AspAT (*CiAspAT1, CiAspAT2;1, CiAspAT2;2, CiAspAT3*) (Fig. 2a). The gene families from Asn biosynthetic pathway appeared to be relatively conserved in size, the differences between species mostly reflecting the differences in ploidy levels (Fig. S7, Notes S1) (Bel *et al.*, 2018). However, it should be noted that our identification of Asn biosynthetic genes might have been partial because it relied on data from preliminary assemblies of chicory genomic sequences obtained through a shotgun sequencing approach (unpublished data) and sequences from publicly available chicory expressed sequence tags (ESTs). The molecular analysis of the *in silico* identified Asn biosynthetic genes revealed expression in both storage roots and leaves from selected chicory genotypes. In storage roots, the gene members coding for AS, GS, GOGAT and AspAT that displayed the highest expression were *CiASN2, CiGLN1;3, CiGLT and CiAspAT2;2*, respectively (Fig. 2b). Four gene members (*CiASN2, CiGLN2, CiGLU* and *CiAspAT3*) did not present differential expression in storage roots in the contrasting genotypes, while others (*CiGLN1;3, CiGLT, CiAspAt2;2*) were downregulated in the genotype L4118-HAsn. Overall, Asn accumulation in storage roots from contrasting chicory genotypes did not correlate with transcripts levels of Asn biosynthetic genes in their storage roots. On the contrary, eight gene members (*CiASN1, CiGLN1;1, CiGLN1;2, CiGLN2, CiGLU, CiAspAt1, CiAspAT2;1 and CiAspAT3*), representatives of all the Asn biosynthetic families, displayed expression levels that were significantly higher in L4118-HAsn leaves as compared to L4115-LAsn leaves (Fig. 2c). These results suggest that the high levels of Asn accumulation in storage roots could result from Asn synthesis in leaves and Asn transport to the storage roots. Noticeably, L4118-HAsn leaves accumulated lower levels of Asn when compared to the L4115-LAsn leaves, suggesting differential source-sink transport of Asn in those genotypes.

**Fig. 2.**
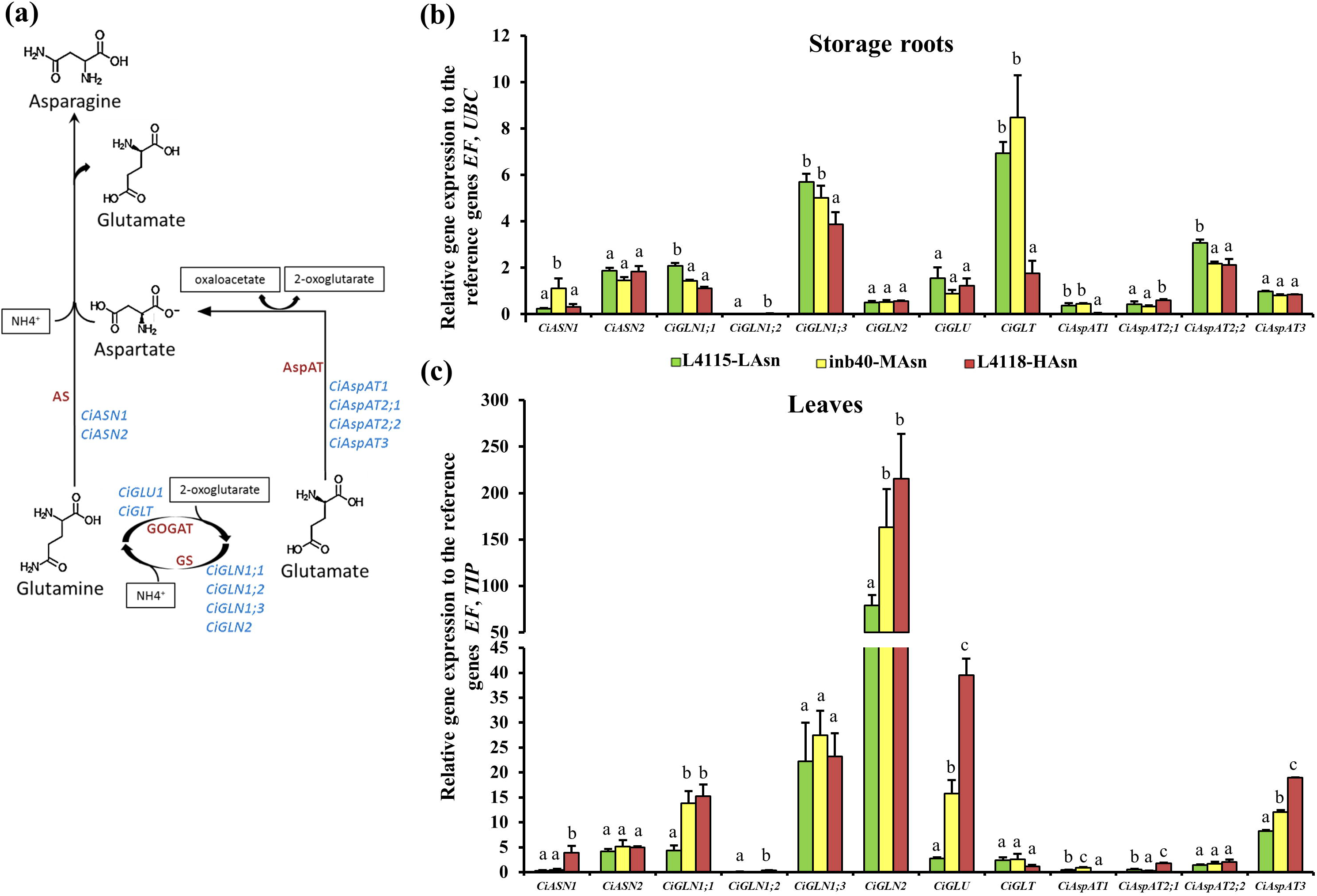
Correlation between Asn content and the expression of the Asn biosynthetic genes. (a) General overview of the Asn biosynthetic pathway in plants: asparagine synthetase (AS), glutamine synthetase (GS), glutamate synthase (GOGAT), aspartate aminotransferase (AspAT); and the corresponding gene members identified in *C. intybus*: asparagine synthetase (*CiASN)*; cytosolic glutamine synthetase (*CiGLN1*); chloroplastic glutamine synthetase (*CiGLN2*); Fd-GOGAT (*CiGLU*); NADH-GOGAT (*CiGLT*); mitochondrial AspAT (*CiAspAT1*); cytosolic AspAT (*CiAspAT2*); plastidic AspAT (*CiAspAT3*). (b), (c) Relative gene expression of Asn biosynthetic genes in storage roots (b) and leaves (c) of *C. intybus* from genotypes with low, medium and high Asn content (L4115-LAsn, inb40-MAsn, L4118-HAsn, respectively). Data are mean + s.d. of 3 to 4 biological replicates. Different letters for each gene member represents statistical significance according to one-way ANOVA, *P* < 0.05 followed by Scott-Knott test, *P* < 0.05.

### The synthesis of Asn in the leaves highly influences the pool of Asn in the roots

To further explore Asn transport and accumulation in source and sink organs, we took advantage of the contrasting genotypes to determine the contribution of leaf Asn biosynthesis to the Asn pool in storage roots. We established a protocol enabling successful chicory grafting (Fig. S8) and we performed heterografting of L4115-LAsn and L4118-HAsn (Fig. S9). Three months after grafting, we measured [Asn]_leaf_ and [Asn]_storage root_ in the reciprocal heterografts L4115-LAsn/ L4118-HAsn and L4118-HAsn/ L4115-LAsn as well as in the control homografts (Fig. S9). The L4118-HAsn homografts accumulated Asn levels in storage roots that were over two times higher as compared to the L4115-LAsn homografts, indicating that the newly established grafting model is suitable to measure differences in Asn accumulation and transport between organs (Fig. 3a). The [Asn]_storage root_ from the heterografts L4115-LAsn/ L4118-HAsn was as low as in the homografts L4115-LAsn (Fig. 3a), confirming the contribution of leaf Asn biosynthesis to Asn accumulation in L4118-HAsn storage roots. Intriguingly the L4118-HAsn/ L4115-LAsn heterografts did not lead to high [Asn]_storage root_ (Fig. 3a). In addition, [Asn]_leaf_ in the L4118-HAsn/ L4115-LAsn heterografts were also reduced when compared to [Asn]_leaf_ in the L4118-HAsn homografts (Fig. 3b). On the contrary, the [Asn]_leaf_ in the L4115-LAsn/L4118-HAsn heterografts remained unaltered compared to the L4115-LAsn homografts. We subsequently measured and compared expression levels of *CiASN1* and *CiASN2* genes in L4118-HAsn leaves from the L4118-HAsn/L4115-LAsn heterografts and from the L4118-HAsn homografts. *CiASN1* displayed a significantly higher expression in L4118-HAsn homografts leaves as compared to L4118-HAsn/ L4115-LAsn heterografts, while *CiASN2* remained unaltered (Fig. 3c). This observation suggests that the sink organ impacts expression of Asn biosynthetic genes in the source organ. The low Asn levels in the L4115-LAsn storage roots from the L4118-HAsn/ L4115-LAsn heterografts also suggest that L4115-LAsn storage roots are impeded in transport and/or accumulation of Asn. Because leaves in the L4118-HAsn/L4115-LAsn heterografts accumulate intermediary Asn amounts (Fig. 3b), we hypothesized that a limited transport and accumulation of Asn in storage roots could increase Asn levels in leaves and thus, negatively impact the Asn biosynthesis there. To test this hypothesis, we established an *in vivo* experiment in which the Asn, Asp, Gln and Glu amino acids were individually infiltrated in chicory leaves. Thirty six hours after the infiltration, RNA was isolated from the infiltrated leaves and the expression levels of gene members were evaluated by RT-qPCR. As a control, we analyzed leaves infiltrated with water. The infiltration of Asn into chicory leaves led to the downregulation of *CiASN1*, *CiGLN1;1* and *CiGLU*, but had no effect on *CiASN2, CiAspAT2;1* and *CiAspAT3* (Fig.3d). We confirmed the downregulation of *CiASN1* by Asn feedback inhibition using a protoplast assay (Fig. S10). On the contrary, leaf infiltrations of Gln, Glu and Asp led to significant increased expression of *CiASN1* while expression of *CiASN2* remained unchanged when compared to controls (figs. 3e-3g). These results differ from *AtASN1* and *AtASN2* expression profiles in Arabidopsis leaves as nitrogen metabolites, including Asn, Gln and Glu, upregulate *AtASN1* and downregulate *AtASN2* (Lam *et al.*, 1998). To further characterize the Asn model in chicory, we quantitated Gln in the grafted plants. Gln serves as substrates for Asn biosynthesis and we found that [Gln]_storage root_ in the L4115-LAsn/L4118-HAsn heterografts were significantly reduced compared to the L4118-HAsn homograft (Fig. 3a). This suggests that the Gln pool in chicory storage roots is also influenced by the Gln biosynthesis and transport from the leaves. In addition, the [Gln]_leaf_ in the L4118-HAsn/L4115-LAsn heterograft was significantly reduced when compared to the L4118-HAsn homograft (Fig. 3b). Because the Asn infiltration in leaves also led to the downregulation of *CiGLN1;1* (Fig. 3b) our results suggest that the decrease in Asn biosynthesis and accumulation could be a consequence of the deregulation of several Asn biosynthetic genes in chicory leaves. The mounting concentration of Asn in chicory leaves due to limited transport and accumulation in the storage roots from the L4118-HAsn/ L4115-LAsn heterografts could initiate and exert a negative feedback loop regulation on expression of Asn biosynthetic genes in L4118-HAsn leaves.

**Fig. 3.**
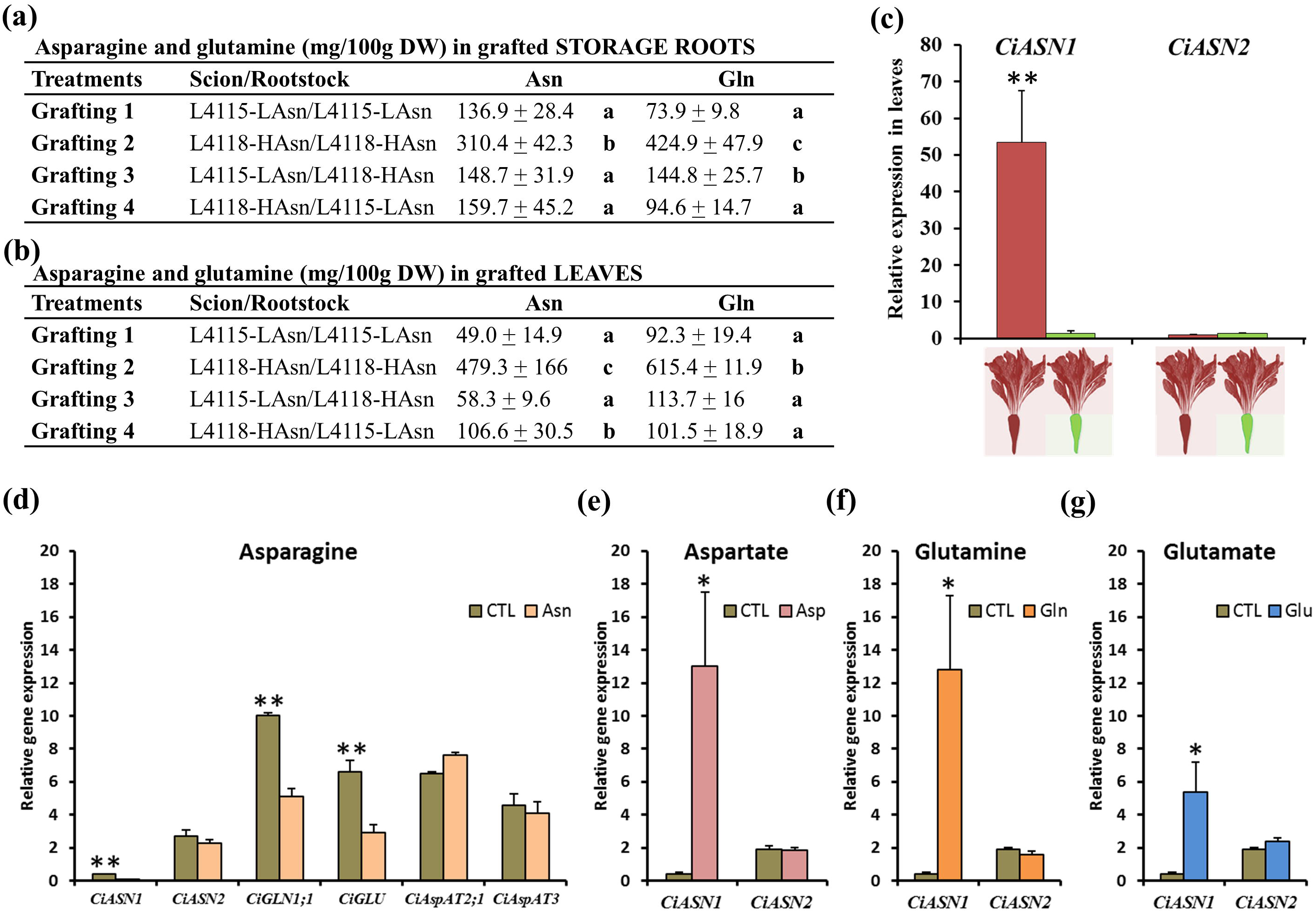
Grafting of genotypes contrasting for Asn biosynthesis and accumulation. (a), (b) Free Asn content in storage roots (a) and leaves (b) of grafted plants at 3 months after grafting, expressed in mg/100g dry weight. A detailed scheme of the grafting experiment is depicted in Fig. S4. LAsn, low Asn; HAsn, high Asn. Data are representative of 3 to 8 biological replicates. (c), Relative gene expression of the *ASN* gene family in leaves from the graftings 2 (red/red) and 4 (red/green). (d-g) Relative gene expression of Asn biosynthetic genes in leaves infiltrated with Asn 190 mM (d), aspartate 37 mM (e), glutamine 190 mM (f), glutamate 57 mM (g) and. Plants infiltrated with water were used as control. Asn, asparagine; Asp, aspartate; CTL, control; Gln, glutamine; Glu, glutamate. Data are mean + s.d of 4 biological replicates. (*) and (**) represent statistical significance according to the t-Test at a significance level of P < 0.05 and P < 0.01, respectively.

## Discussion

Asn is considered to serve as a major transport compound in the xylem from the root to the leaves and in the phloem from the leaves to the developing seeds in several model and leguminous species that have been investigated (Amarante *et al.*, 2006; Lea *et al.*, 2007; Krapp, 2015) This model is also corroborated by the high nitrogen to carbon ratio of Asn (Coruzzi, 2003) Previous studies aiming at determining the AA potential in crop species have revealed that genetic variability exists in the accumulation of Asn in plant organs (Halford *et al.*, 2012; Postles *et al.*, 2013; Curtis & Halford, 2016; Muttucumaru *et al.*, 2017; Curtis *et al.*, 2018) In the present study, we could identify chicory genotypes contrasting for Asn accumulation in storage roots and demonstrate that diversity in Asn accumulation relates to the expression of Asn biosynthetic genes. Noticeably, we found a positive correlation between the gene expression of the Asn biosynthetic genes in leaves and the Asn level in storage roots (figs. 2b and 2c), suggesting that Asn biosynthesis in leaves have an important contribution to the Asn pool in storage roots. This feature could be organ- and species-specific as suggested by previous studies in potato demonstrating that tuber accumulation of Asn mainly occurs through Asn biosynthesis in tubers with minor contribution from the leaves (Chawla *et al.*, 2012).

Our grafting experiment with genotypes contrasting for Asn accumulation demonstrates that a reduction in the biosynthesis of Asn in the leaves leads to a reduction in the Asn accumulation in the storage roots (heterografts L4115-LAsn/ L4118-HAsn, Fig. 3a). The expression profile of Asn biosynthetic genes in contrasting genotypes (Figs 2b and 2c) further indicate that the pool of Asn in the storage roots originates mainly from the Asn synthesized and transported from the leaves. Moreover, our experimental data also shows that the accumulation of Asn in storage roots plays a regulatory role on the Asn biosynthetic pathway in the leaves (Figs 3a and 3b; Figs S8 and S9). The observation that the [Asn]_storage_ root in the L4118-HAsn/ L4115-LAsn heterograft was as low as in the homografts L4115-LAsn (Fig. 3a) suggests that the functionality of the Asn transport system in chicory is dependent on the activity of transporters in both leaves, for phloem loading, and storage roots for phloem unloading through the apoplasmic pathway and/or post phloem transport via the apoplasm. Although the apoplasmic pathway of phloem unloading for C and N photoassimilates remains poorly understood, studies in crop and model species have demonstrated a key role of the apoplasm in the transport of hexose and amino acids (McCurdy *et al.*, 2010; Besnard *et al.*, 2016; Milne *et al.*, 2017) Transporters from the UMAMIT family have also been shown to be important in the post phloem transport of amino acids via the apoplasm (Müller *et al.*, 2015). We hypothesize that regulation of Asn transport in the phloem from loading to unloading steps contributes to the genetic diversity in Asn accumulation in chicory. Our grafting experiment also suggests that the accumulation of Asn in phloem tissues and/or the xylem transport of Asn from roots to leaves could contribute to a feedback regulation of the Asn biosynthesis in leaves. The reduction of Asn accumulation in the heterograph L4118-HAsn/ L4115-LAsn as compared to L4118-HAsn homograft (Fig. 3b), was also concomitant with a significant reduction in the leaf expression of *ASN1* (Fig. 3c). One hypothesis that can be derived from our aforementioned observations is that the reduced phloem unloading of Asn in the rootstock of the L4118-HAsn/ L4115-LAsn heterograft causes a negative feedback loop in the Asn biosynthetic genes in the leaves. Our leaf infiltration and protoplast experiments support a negative regulation of several Asn biosynthetic genes (i.e. *ASN1, GLN1;1* and *GLU*) by Asn in chicory (Fig. 3d). Pioneering work in Arabidopsis has demonstrated inducing and repressive action of sucrose on respectively *ASN1* and *ASN2* transcript levels as well as the partial reversion of those alterations when supplementing Asn to seedlings (Lam *et al.*, 1994, 1998). While studies in other plant species has shown the regulatory potential of certain amino acids on their biosynthetic pathway (Lam *et al.*, 1998; Gutiérrez *et al.*, 2008; Zhang *et al.*, 2015) further investigations are required to verify whether the Asn negative feedback regulation we characterized in chicory can be expanded to other plant species.

Based on the aforementioned results, we elaborated a model depicting Asn biosynthesis and accumulation in chicory leaves and storage roots (Fig. 4). In this model, the pool of Asn accumulating in storage roots originate mainly from the pool of Asn produced and transported from the leaves. Thus, plants with a defective transport system either in the leaves or both leaves and storage roots, accumulates high [Asn]_leaf_, which in turn downregulate the expression of Asn biosynthetic genes. On the contrary, plants possessing an efficient transport system can translocate Asn into the storage roots, hence keeping [Asn]_leaf_ low and thereby limiting the negative feedback loop on the expression of Asn biosynthetic genes.

**Fig. 4.**
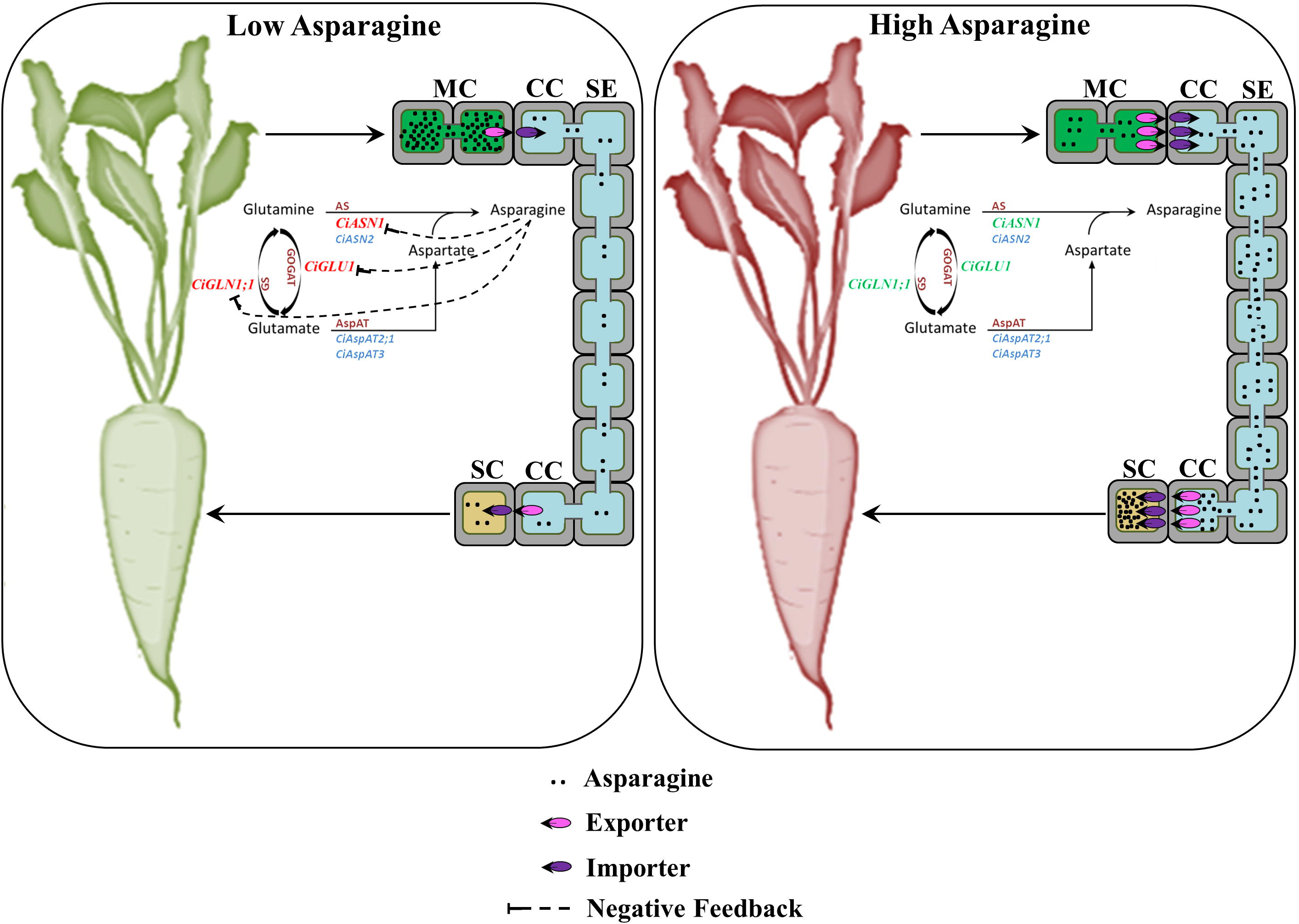
Model for Asn transport and accumulation in chicory storage roots. In our model, plants with low Asn content (represented in green) either lack or have lower transporter (importers and exporters) activity controlling the Asn transport from leaves to storage roots. Thus, Asn accumulates in leaves and in turn downregulates *CiASN1* and other genes in the Asn biosynthetic pathway. The overall outcome of this is that less Asn will be transported to the storage roots. On the contrary, plants that accumulate high Asn levels in storage roots (represented in red) have a high activity of the transport system in leaves and storage roots enhancing the transport of Asn from leaves to storage roots. The outcome is that *CiASN1* and other genes in the biosynthetic pathway can be expressed at high levels in leaves and thus, produce more Asn that can be efficiently transported and stored. Our results showed that *ASN2* is expressed at similar levels in the leaves of both (Low and High Asn) genotypes and is not controlled by the amino acids of the Asn biosynthetic pathway. Therefore, *ASN2* is proposed to be involved mainly in the role of NH_4_^+^ detoxification in chicory as observed in potato^8^ and Arabidopsis^13^. MC, mesophyll cells; CC, company cells; SE, sieve elements; SC, storage cells.

Using chicory genotypes contrasting for Asn accumulation, we could demonstrate that Asn accumulation in storage roots depends on Asn biosynthesis in the leaf. Based on grafting experiments, leaf infiltration of amino acids and gene expression studies, our results suggest that *CiASN1* is a key player for Asn biosynthesis in the leaf and that Asn acts as a repressor of *CiASN1* expression. Our model indicates that Asn transport and biosynthetic gene regulation by Asn are key mechanisms controlling Asn accumulation in plant organs. The feedback regulation between source and sink we report for the first time in chicory is likely to open new breeding perspectives to generate crop varieties with reduced AA potential. Our findings also highlight long-distance transport of amino acids and Asn transporters as central targets to reduce accumulation of free Asn in storage organs.

## Supporting information

Sup NotesS1

## Acknowledgments

We thank to The Flanders Research Institute for agriculture, fisheries and food (ILVO) to provide part of the chicory genetic material. This work was supported by a grant from BEWARE a programme co-financed by Wallonia and European Commission

## Author contributions

Conceptualization, E.S. and H.V.; Methodology, E.S., L.S. and H.V.; Validation, E.S.; Formal Analysis, E.S.; Investigation, E.S., L.S., N.D., P.C.; Resources, H.V., C.N., O.M.; Writing - Original Draft, E.S. and H.V.; Writing - Review & Editing, E.S., H.V., L.S., N.D., C.N.; Funding Acquisition, H.V., E.S., O.M.; Supervision, H.V.

---

**Fig. S1** Diversity of Asn accumulation in the chicory germplasm.

**Fig. S2** Correlation between weight (g) and Asn content (mg/100g)

**Fig. S3** Phenotypic variation of diameter (a) and weight (b) in chicory storage roots 180 days after sowing.

**Fig. S4** Free Asn content in storage roots from 18 genotypes of chicory measured at 30, 60, 90, 120, 150 and 180 days after sowing.

**Fig. S5** Stability evaluation, over 2 years, of the Asn content in 7 genotypes of chicory representative of the three contrasting groups to Asn level in storage roots.

**Fig. S6** Correlation between Asn content (mg/100g) in leaves and storage roots.

**Fig. S7** Number of genes from the Asn biosynthetic pathway in *A. thaliana* and crop plant species with high acrylamide potential.

**Fig. S8** Grafting method for chicory.

**Fig. S9** Overview of the grafting experiment.

**Fig. S10** Asn feeding experiment.

**Fig. S11** Determination of the optimal number of reference genes, according to the pairwise variation V from geNorm.

**Fig. S12** Selection of the best reference genes for normalization in storage roots (a) and leaves (b).

