## Supplementary material for "Asparagine accumulation in chicory storage roots is controlled by translocation and feedback regulation of asparagine biosynthesis in leaves": Sup NotesS1

**Supplementary data 1.** Genomic and RNA sequences (blue) from *C. intybus* obtained through a shotgun sequencing approach (unpublished data) and publically available chicory expressed sequence tags (EST) available in:

<https://www.ncbi.nlm.nih.gov/Taxonomy/Browser/wwwtax.cgi?mode=Info&id=13427>

***CiASN1***

>scaffold11106

AATTCTTATTGATCTTAGATTTAACTTTATTTTTGTCTAATAAATTAAATGTCATAAAATTAATAATTGTAACTTGTAGTAGAATATAAGAAAATGGATGAAAGATATAAAATAAGTATAGCATGTATTTAAAAAACAAAAATAACATAACGTTTACATAACAGAACCCCACTCTGTAGTTTTTAACTCAAAAAAGACTTCCCTTGAAGTTGTACACACAATGTATGCGTTATACAATCTTTAAATATGTTTTAGATAATACTTCGACTGTATACAAATAAAAAACCACAAAAGTTGAAATGATATTGTTAATCCTTTCAAAATAGTGTTTGGATATTTTTTTCGTCCTGCTTTTATACATAATAATATAATAGTATGTCGTTTAAAATTTAAATGTTTTAATTAACTAAAGATTAATAAATATTATAACCAAAAACTATTGTTATAATGAAATAACGTAACTTAAAATAACATAACTTAATTTTTAAATTTGCGAATTTTTTGTTTGATAGTATTTGTCTGAAGTTTTGTTTTGTTTTCTACAGCTAAACAATTATCGCTTTTTTTCGTATGAAATTATTAATAAAAATACATATCAATAGAGAAAATCTCTAAAATATCTTAAGATTGTGACAAATAATATACAATTTGACATACAATTAGTGATTTATCTCTATATAATTAGGATAGAGATTAAATCATCCATAAAACTAAATTACGGCTTAATACTCAAATCATCGAGTTCATGCCGCGCAGCGTATCAGAAAAGTAGAAAACGAAAATATAAAATAGGAGAAAAAATAGTGAGAAAAACGTCGCCGCTTGAAAATTAATAAACAAACAGATTTGAAGAAAATGTCGATGGGACCAGTCGCCGGAACCGTCTATAAATTAGCGGCGAATGAAACGGTGATAACCAACCAATTCCTTCTCACTTCTCAGCGTCCACATTTTACCCTTTTACGGTCATCTTCTTCGCTCTGCCCCTGGTTTTCTTTCCCCGTCAAAATTTGTAAATTTCCATCGTTACTTGTACGGTAGCCATGTGTGGAATACTAGCTGTTTTGGGTTGCTCCGATGATTCTCAAGCCAAAAGGGTCCGTGTCCTCGAGCTCTCTCGCAGGCATGTTTCCCAGATTCTTTTAATTTAATTTAAGAACCAACAAACCCAGATCCATAAATCAAAAGTAGAAATTAGTTACCAAAAGTTTTCAATGTTCTTAATGAAAATTGTTGTACAGATTGAAGCACCGTGGCCCAGATTGGAGCGGGTTATATCAGCACGGTGATTGTTATCTGGCGCATCAACGTCTCGCCATCATTGATCCTGCTTCCGGTGATCAACCCCTCTATAACGAAGACGAAACGATCGTTGTAACTGTGAGTAGAGTGACGATAATGATGAGTCGGTTCATTAGGTTATTAAACCGTTTGATTCTTAGGTCCATTTTTGTTCGAAAATGCAGGTTAACGGCGAGATATACAACCACGAGCAGCTCCGAGAAAGCTTAACCGGTCACAAGTTCAAAACCGGCAGCGATTGTGATGTTATTGCGCATTTGGTGAGTACTTACGAAGTTTTCAGTAAAATTATTGTTTAGATTTTGGGTTTGAATGTAATCGAGCTTTTTGACCTTTTATAGTACGAAGAACATGGCGAAAACTTCATCGATATGTTGGACGGAATGTTTTCGTTCGTGCTACTGGATACACGCGATAACACTTACATCGCTGCTCGTGATGCTATCGGGATCACGTCGCTCTATATCGGTTGGGGACTCGATGGTGATTATCAAACGCTTTCCGTTACATTATTTGTTTATTACAGAGATAATAGAGTAAACGAGTAACAATCGTCTTCTTATTCAGGGTCGGTTTGGATTTCGTCTGAGCTGAAAGGTTTAAACGACGACTGCGAGCATTTTGAGGTGTTTCCGCCCGGCCACTTGTACTCGAGCAAAACCGGTGGATTCAAGAGGTGGTACAACCCTCCATGGTTCTCCGAGGCCATTCCATCTACACCATATGATCCTCTCGTTCTCAGGCGTGCCTTCGAAGACGTGAGATTTCTTTCTTTAACGTTTTCCAAATAAATGTTTGTTTATATGAAGATAAGTGTCTAAAAGAACGATAACGTGAACTGTAGGCCGTAATCAAAAGGCTAATGACCGATGTACCCTTTGGGGTTCTTTTATCGGGAGGGCTAGACTCGTCGTTGGTCGCATCTATCACCGCCCGTCACTTGTCCGGTACAAAAGCTGCTAAACAGTGGGGGGCTCAGCTTCATTCCTTCTGTGTCGGCTTAGAGGTAACATACGATCGATATAAATCACTTGATTCCATTTCCGAAACTGTATTCAACGAAAAACTGAACCACCAACTTCACCCCATAAACAGGGTTCACCGGATCTCAAGGCGGCAAGAGAAGTAGCTGATTATTTGGGAACTGTTCACCACGAGTTTACCTTCACTGTACAGGTAACTTACTATATTCTTTAGGTCCGAACTGGGCCGTTTTCGGTTGGCCGGATCTTATTTCGAGTTTTCCGACGTCAAGGTCTTGATTTTGGATTCTATTGGTGTGTAGGATGGTATTGATGCGATAGAAGACGTGATTTACCATATCGAAACATACGATGTGACAACGATTAGGGCTAGCACACCGATGTTCCTGATGTCACGGAAGATCAAGTCGCTAGGCGTGAAGATGGTCATCTCCGGTGAAGGCTCCGACGAGATATTCGGCGGGTACTTGTACTTTCACAAGGCACCCAACAAGGAAGAGCTCCATCGCGAAACCTGCCGCAAGGTAAAATCACCGACTCTCAGTCTCACAGTAGAATTATTTTCAATTTTGGGGTTTTCAAGTTTTTAACCTGATTTACACAACGCAGATAAAAGCACTTCACCAATACGATTGTTTAAGAGCAAATAAGTCGACCTCGGCTTGGGGTTTAGAAGCCCGAGTCCCATTTTTAGATAAAGAATTTATCAACGTTGCCATGAGCATCGACCCTGAAGCAAAAATGGTAACAACTCTGCTTCTGTTATAATAAAAAATTCTACGATTTACCGAAAGTACCAAACTTAATGAGGTTATGTGCAGATCAATATGGATGAAAAACGGATAGAAAAATGGATTCTGAGACGTGCTTTCGACGATGAAGATCACCCGTATTTACCAAAGGTCTCCAAACTCGATCTGTGAAAATAAAACTCTGTTTCGAATCATAACTTTTGCTCTTTTTAACATCTCTCTCTCTCTCTCTCTCTCTCTCTCTCTCAGCATATTTTGTACAGACAGAAAGAACAGTTCAGCGATGGCGTCGGGTACAGCTGGATCGACGGACTCAAAGCACACGCTGAACTACACGTAACAGACAAAATGATGCTAAACGCCGCTCATATCTTCCCTTTCAACACTCCGGTCACTAAAGAAGCCTACTATTACCGAATGATCTTCGAGCGGTTTTTCCCTCAGGCTAGTTCTTAGATCTTAGTTACGACTTGTACATCATGCAATAACTCAAGTAACTTGAGCCCTTTTTTGACGTTTACTTTTTTTTTTTTTTTTNNNNNNNNNNNNNNNNNNNNNNNNNNNNNNNNNNNNNNNNNNNNNNNNNNNNNNNNNNNNNNNNNNNNNNNNNNNNNNNNNNNNNNNNNNNNNNNNNNNNNNNNNNNNNNNNNNNNNNNNNNNCAGAATTCGGCAAAGTTGACCGTTCCAGGTGGAGCAAGTATAGCTTGCAGTACTGAGAAAGCAATCGAGTGGGATGCATCTTGGTCAAAGAATCTTGACCCATCGGGAAGGGCTGCATTAGGGGTTCATAACGATGCATATAAACAGAAGGCAGGCCCAATTGCTACAGGGAACTTGGCGACAAGTATGATCGATGATGTGCCAAGGATGATGGATATTCCGGCCCATGGGGTTGTGATTCAGGGCTAGACAATTGTTGTTGTATAAGTTTTTACTACGATATGTTGATGTGTTAGCTTGTTGAGTTTGTTTAAATTTGTGTATTAATAACTCGAATGTATCGTTGTTGATAGACTTGGTAACATTTGATGTAATCTGTCTGTAAAGTTCGGTTTCTTGAAGGCACTGTTCCCTAGTTCTTCACTCCGTTTGTCAATCAATAATCGTTAAATATGGAACTTGTTGATTATATCAGGCCAATAAGATCTTTATATCAACCGAATCAATTGGGATTTCATATAGAACATAAGGAATTTACTTGGGAATCTGATTATATGGATATATTTTGACATGACTCCAAAGTTGCACCATAACAATCGAGCAAAAAGCAATACTGAAAGATTTACATGGTTCAGAAATGCACAAAAACCAAAATTTGAGGTCTAAAGATGTGCCCAGTTGCCAAGAGCTCATGTTGCAATACGGCAATGCCTGGTAATCTGGTATACAAGGGCCTCTGGCTTTGGGCCTATCTAGTCTACAGTTGAAAAATAACATATAGTGATATAGAAAGTTCTTGGGCTTAGAGCCTAATCAAGCTTTCTAGACCTTGGGCCTAATTTATATATATATTTGTTTCTTAAACCTTATCTTCTTCAAAAGACTAATAAAAACGTAGCTTCTACATACGTTTTTCAAATATCTTCCTTTACTAGTAGCATTTTGAGAAAGATTTATTATCCTATTTATGTAGTTTGTATTCTTCTTGGAAAAAGCTTTGTAATTATATATTATCAATTCAAATAAGAGAAAAGATATTGACTCTCAATTTGAGATATTTGGGAAGAAAATTACAACTTGAAAGTGGAGAAATTAGATCTTCTCCCTAAGTGGGAAGCTAAATCATACTGATTTTGTAAAACAAGGGATCCTTTACAACCTAGTTAGATGGTGATGGTCTATGTGCGATACCATGCATGTGTGCGCACGTGTGGCTATTGTTGTGGATATGAGTTGTGATATTTTGGGAGTGATCAATGTAGACGCATTTTCCATTTTTTTTCTTTGCATTTTACAATACAACACCCTGGTGCTGAATCACCCATGTTTTAAAATCGTACATATCGTGGTGTTGAAGTTCGGCATGGTCTTGAATACCTTGTGCCAAATTTGCATAGGTCTTGAATGTTGTGGTTCCATAATATTTACACCATCTACGTCATTGTCAACACTTATTAATACCTTAAACTGAATGAATGAGTTCTAATAGGGGTTAATGAACATTTATTTTGTAGAGCAATGGTGGTAGTTTGAGGTTGAACTTCACTACTATAAACTGAGAGTCATACTCACAGGCTAGAAAAAGAAATACACGCAAAAAATATGCTTGTTTCAAGCAACCCACAACATAGATAGAGATTAGAGAACATGTATATGCTCCGATGAATTTTCATGCAAATTGTATACTGCTATAGTTGGTGTATCGTGAGAAACAGAAGCATGAAACAAGCATATTTTAATAAAATTGTCTTTAGCACAAGCCTCAACCATATTGTGTGCAATTTCGGAATTTTTTATTTTATGACTTCCAGTTGCATCGTACTATTCTTTTGAAAAACAATGTTTATGTTTCTTTCCTTTATAACCGAATTAATAAGCACACTAAAAGAACTGCATAAGTTTTCATATGACAATCAAGACTTAGAACTCTATCAAAGCTCTATTTAATACTTGAGATCATCAATTAGCTCAATAGAAGTTGTTAGACACATAATTTGGATTCATCGCTTATTTTGAGGTTAGGAAACTCATTCTAATCTAAATATAACACCATATTTAAAGTTCAAAGCTCTTAATCAACATTTCACTAATCTAGGGTTCATAAGCTTCAAAATAACTCATGAACTTGCATCTAACCAATATAAAATTATGATATTTATAGATTAGTCAATAAATGATGAAATTCAAACTAGAAACTCAATCCAAATAATAACTATATAAAAACTTTGGATTAAAGAAGAAATCTAAACCATATGTTACTAGAATGAATATAATGGGTAGAGTTGATTACATACTTCTAATCTAGATGAGTAATTAAACTCCAAATCTTCAATTGAACTCCAAGCTAGCTTTAATATATACCAAAGCTTGTGATATTGTCTCTATAACTCCTAAAGCTCTCCAAATTTCTTTTCAAATGAGGGTGAAGAGGGAAATGTGTATATTTGACTTAATAACATTTTAGCCCCTAAACTCTTAAGTTTCCAACTTGTCTCCCCAATTAACTCCGAAAAGACAAAATGTATTAATTCTTCCACATAAATCCTAGAAAGTGAAGTTCTTAAAACTTTGGACCCAAAACTAGTTTATTCAATGTCAAACTTAAGTTTATGAATTTAGCAATTGGTCCCTCAAACTTTAATTTTTAGACTTTGATCCTGAAATAACATCCACTAATATTTATTGTAGAAACTCCATATCAACCCTCGAAAGGTTAATTACTTACAAAGTAAATACCAAAACCTACTTCATCAAGTATATCTAGTCATGCCAAAGTTTCCGAAACTTTGTAGAAATAGTCATTCACCATCAAAATCTTAATTTAACCCTAAGATAACATTCTAATTAACATTTATAGTTCCTCATATTCTAATTCCATCGTAATTAATAAAATTCTTTCTAATAAGTCTATTTCATAATTAAATGGGTTTACTCGAATGTTATTCATTAATTCCTAAACTTAAGGTTTTTAACTTCAACCATCACCTACATACTATAACTAAATTTTATTTTAACTGATTTTGAAACTTTTAGATATATTATTTTAATTTATTTTATTTAATTCAAAAATTCAAAATAACTTCACTTAAAGAAGTAAAATTTGTAGCCCCAAAAAATGTAGGCGTTACCAGTGTTGCAAAACTCGCTAGGCGCTGCCAGTTCGGTGGACGAGCGAGTAGCGAGTAATCGGTTAGGCGGATGAGTATTCGGTGAGTACTCGGTGACACTGTTCATGATAATATTATATTAGTGTTGTAAAACTTGGTCGGCTCGGCCGAGTGCTGGGACTTGATATACCTAATTAATAAGATAGTTGTAATTTTTTTAGGGGTGAAAAATGACTAGTTAGAAACTGCTTATTTTAATAGTTGGCTGTTGAGGGTTTAAAGGGATATTTAGGAGGACTAGAGGGGCTTTTTTTGCGTAAATCCATTTCTTGATCTTTGAACCATCGGGATTTAAGGAGACCGATTGCAAACCGATACGATTGTCAAACTGTAATTGGAAGCAGGTTCGCACTTGGTTTCGCGTATCAGAAGTTCTAGATTCGATTGAAACAAATCACGTTGCCGACTCTGGAACTAGATTGATTGAAACAAATCGATTACGTATCAGAAGTTCTGGAAAACGATTAGAAACGATTGGAACAATTAGAAACGAAAAGATGATAGTTTGACCTTGGAGAGAATCGAGTTCAATACCGTCTACGATGGCAATCGAGCTTGGAGTAGGAACAGGTCGAGTTGAATCTACGTGAAATCGAATGGGAAAGAAAGTATTGTTCAGTCGGATTCAACAAGTAGCGATGCTGTCGTTTGCTTGTAATATTATCCCATATCGCCTAAAACATAAATAAGCGAGGGTTCCTCTAGTTATAAAATAGCTGCTAAGGTTCACAGATTTTCACCCATCAAAAAGCCTTCAAATTGAACCGGGTTTACTCCTGGATCGAGTTTGGTCTAAAAATGGTCCGAGTTTGGGCTTAAAATGGTCTGAGTTTGGCACATTTTGGGCCGATTTTTCCTGATTTGGGCCGAGTTAATCGGATGTTGACCGAGGTTGACCGATTCCGATTTTTTGGACCGAGTTGGTTAAACTCGCCAAGGTCAAGGACCGAGTCCGAGTCCAAGGCCGAGTACTCGGCCGAGTTGGCCGAGTTTTGCAACACTGGGCGTTACACTTCAGAACAGTAGTTCCTGAATCAGATGTCTTTGGACCAAAGTTTATCTAACAAGCCATTCTTTAGACCTAAAATGTTTAGACGAACGAACTTATATCAACGGAACACTCTCAACTGTCTTGCATTAAAGGCCTTAGATCTCCGAGAGATTTGCATTATCAACACTTCTGACTAGTTAATCTAGGTCTACAGACCATAGGTTCTCAAACAGAACCTCCAGACTAATAGTCTACAAACTAGAGGTCCTGAACCAGCTGATGCTTCAGACCAATGGTCTATAGGACAGTAATATTGGGCCATCAATTATTTTGCATCAAAAGACTGCTTTAAAAGTCCCAGGTCAGTTTTGTTGATTTGTCAACATTCTTCGGTACAACAGAGATTTATCCATAACATTTTATCATAAGGGATAGCTAAAAAGACACCCTTATTTACCAATTAGACACCTTAATCATTACTTTTAAGTAGNNNNNNNNNNNNNNNNNNNNNNNNNNNNNNNNNNNNNNNNNNNNNNNNNNNNNNNNNNNNNNNNNNNNNNNNNNNNNNNNNNNNNNNNNNNNNNNNNNNNNNNNNNNNNNNNNNNNNNNNNNNNNNNNNNNNNNNNNNNNNNNNNNNNNNNNNNNNNCATTACTTTTAAGTAGGGTCAACTGTTTTTTATTTGTTGGAATTACAATGTTTTCCGGAAAAACCGATATGATTGTTTAAGAATGTTTGACTTTTATTTATACATGTATATTGAGTGTCTATTTAGTAGCACCCTTATCATATTCATTATAGCAGCTAAACGCCAAAATCAATAGGTCTACATTTGATATGAAAGTTATATTTAACTTTCATTCTCATGGCTTTTCGAGAAGCTTCACATATAAAAAGGGAGCTTACAAAAACTTATTTTGTTGATCAGACATTCTCTTGTCCACTTTGATTATTTTTTTATCATTGTCTTTCAGTAAAAAAGAATAAGGGGTTAGTTATAGTATGCACATTTGAAGATTTACTTCGTATGGTTTTTAGAGTGTGGATTTTAAAAAAAAATAATGATGAAGAAAAGAGATTAATTAATGACGTGATATGGCACTGATTTTTGGGGTTTTGATCAATTTTTTGGCCTCACTAATTAATCTTGATAGTTAGAGAAATCAATTAGTGAAGTGATGTGGCATTGATTTTTGGTCTTCTAATCAAATATAATTTTACTACTTGATTTCAATCTTATATTTGATTGGATTTGTTCAGTTCCCATATTTGTATTGTTCGGTATTAAATAAATCAACCCAAGAAGCTTCCTACTTTTAGTTAATATTCTTTTAAAGATTTTTCTTTTGTGTATTATGATATAGTTCTTCAGATATTTGAGGAACAAGAAATAGGTCAGGATTTTTGAAGCCGAAGTATGATTTGATGATGGGATATTTAGTTCACGCGTTCTAGAACCTTTGCACTTGATTTGCAGGGGACACTAAAAAGCTTTTTAAATTGAGAGGTTGTATATATTCCTACTAAATTTTTCTACCCATCTTCCTACTCACTAGATCAATACAACTGTTAAAAACAATGGATGAGATTAAGAGAAAGGATTAATGGATTCCACATGACATTATTTAGTGAGTAGGAAGATGAAGTAGGAAGATTTAGTATGAACAAATATAGTTTTCCTTTTAAATTTGGATCTTAATTTTCTGTCCTAAAGAAACTTAATGGGTACTCCTATATACACGAATCATTTCGTGTATGGGTATTGGTAAAAAAAAATCATGTGATGTGACATGTGTGTGTGAGAGTCAACCCACACAACACACTCACCTACATACATTGAGCGGGTTTTTTAAGATCACCCTTCGTGTATATAGGCATCATGGTTTTGGGGAATTGTAACGTAAACTGAATTTCTTTTCTGACAGTGATTGGACTTTTCTATTTTTTCCATTTTACTCCTTTGAATTTGAACAAAAGTTTCATAACATACCCCTTTTGATTTATATTATTTTTAACGCTAATAATGCATATTATATA

***CiASN2***

>scaffold46890

AAGAAAAAAATAAAAAATAATATCAAGGAGGGTATAATTAGATAATAGGGGGTATGTATAGAATGACCATTTTTTAATAGAAAGTTGACAAAATATAGTTTTTCTACCAAAAAGGTGAAATTACATTGTTTGTATTTTATAAATATATAATTATGCTGAAAAATATAAAATATATGAACATTGTAAAAACCGGGTTTTAAAAAAAATATTCTAAAACTCAGGTTTTTAGTTGACCCGGTGTGGTGACCGGGCCACGGGTCAACCGGTTCAACCGCCATGTCAACCTGGTTTTTACAATTTACGTATAGTGTTATTCTTTTCACCATAATTATATATTTATAAAACATAAACAATGTAATTTCATATTTTTGGTACCAAAAAAAATATTTTGTTAAGTTTCTATAAGAAAGTTCTATTAAATTTGATAAATTTTGTTTCAACCGTCGGATCAACCCGATTCTGGGTCACCGATCCAACCGGTTTTTTGAGAAAACCAGCTGGTTCAACCCGAGTCAGAAAATTAGCAAAAACTCGATCATAATTAACCCGCTTTTTTCATCGGATCACGGTCTAACCGGTTTGACCACCCGTATTTTAAAACATTGGTAGAAAAAAATACAATTTCTCGTACTTTAATTATACAGGGTCAATGTTGATGGGTAAATACATGATCGGGTTTCGGTTAAAACCCGATTAACTAGAAACGTCACATAAAAATATTCATAAGAAATTGAATGGAATCTATTCTTGATGTCCGAGCTGGAGTAAACCAACGGCCCTGATTAATCGAATCGATTTTACATATTCAGCGCAGCCAGGCCTCCATTTCCCCACACACACACTGTCTCCACGATTCTCTTCTGCATGCCCTTGTTCCATACCTAATTCTCTGTGTAGATTCTTAAGCCATTTTTGATTGATTTTAGAAACTGCGAGACGCCATGTGTGGAATACTCGCGGTATTTGGCTGTGTGGATTGCTCTCAAGCTAAGAGGGCCAGAATCATTGAACTTTCTAGACGGTAATGCCATCTTTTCCACCATTTTCCTTGTTTTTTTTTATTTATTTTCTGTTGATACAATTGCCTATTGCTTCCATCGTTGATCACATCCGGCCATTGTATATCATCTGTCCATATTTCTAAATTTCGTTTGTTGCTATTTGTTTCATTTCCGACGAATTTATGTGCGTTTTTTGTTTCTTTAGAAACATCGAACATTATATATCGCTTTGTGATGAAGAACAATACGAACTTCGCGTAATCTGCATCAGTTTGATGCCTAATTCTGTAGGCATTAATTTTATCCAAAAAATAGGAGAAATGGTAGCTAAGGAATGCCATATGCGGTGATAGGATTAAAGGAATATCATATATAGGAGCTAAGTAAAACGACTGGTACCAGTCCGGTGGCCTAAAGCTGCTAGATTACTGTTCTATTCTCTTATGGATTATTGATTAATCTCTACTTAGACGATTAAACACTAGAAGGTCACTTCTTGCGACATTTAATGCTTTGTAAAACAGTAAATATCAAACATCTAATTAACCTAAACCACAACTTTTGAATCCTCAATTTCGTACTCATTGTATTCATGGTCAACAACACACTCAAAAGCACCAAAATATGCACATTTGGTCCTGTAAAACAGTATACATTACCTGCCTCTTGACTATGAATGATTTTGTGAACATCTGAATTACATTAGATAACTTTTTAATTATAAATTTCTATTAACTGTTGCAGGTTGCGCCATAGAGGCCCTGACTGGAGTGGGTTGCATAGTGAGCAAGATTGTTATCTTGCTCATCAACGGTTAGCCATTGTAGATCCTGCTTCTGGTGATCAACCACTTTATAATGAAGACAAAACCATTATTGTCACGGTAACTGGAATATAATTTGACAAAACATGTTTCTTTTTTATAATTCTTGTATAATGAGGGTAGCCAAATGTTGACTTTTCTTTAGGTTAATGGGGAGATATACAACCATAAAGCTTTAAGAGAGCAACTAAAGTCTCACAAGTTCAATACTGGAAGTGACTGTGAAGTCATTGCCCATCTTGTAAGTACCAAATCAAGAAAAAATGATTTAAAATCAAATACCATTTGAATATTTTGTGTTAATTTGGGTTGTGGATTTACTAGTATGAGGAATATGGAGAAGACTTTGTGCATATGTTGGATGGAATGTTTTCATTTGTGCTTTTGGACACTCGTGATAAAAGTTACATAGCAGCTAGAGATGCTATTGGGATCACCCCTCTTTACATGGGCTGGGGCCTTGATGGTATAAATGTTAAATATTCTTTTTCTTTTTTTTTTTGGTNNNNNNNNNNNNNNNNNNNNNNNNNNNNNNNNNNNNNNNNNNNNNNNNNNNNNNNNNNNNNNNNNNNNNNNNNNNNNNNNNNNNNNNNNNNNNNNNNNNNNNNNNNNNNNNNNNNNNNNNNNNNNNNNNNNTTTTTTGGTAAATAAATTGTAGGGAAATACTTAAATTTATAAACTTGGTATTTCATGTGCAGGTTCTGTGTGGTTTGCATCAGAAATGAAAGCCTTGAGTGATGACTGTGAACAGTTTATGTCATTCCTTCCTGGCCATATATATTCTAGCAAAATCGGTATGGATTATTCGTTCTTTAATCAACCAATTTATTACTTTTTTGAATATTTATGTTCATAATTGTTTGATATTGTATTACAGGTGGGCTAAGAAGGTGGTATAACCCCACATGGTATTCAGAACGCATTCCTTCAACACCATATGATCCTCTAGTCTTACGTCATGCCTTTGAGAAGGTTTGATACTAAGAAGAATAATCATATCAATAATATGGGAAAATTACACAAAAGCCCTTACATATTCGCCCTTTTACCTAAAAAGCCATTATGAAATTGTTTTGTGCATAAGGGAAATCTCCAATAAAGTCCTAATATTTTGGCTCAGTTTATGAAAAAGTCCTAAATTAATTTTTATTAACAAAAAAGTTCCAAATTAATTTTTTATTAACAAAAAAAGGACAGAAAACTGGCTAATGTTGAGACTTTTCCGTAAACTGTGCAAAAATATTAGGACTTTATTGAAGATAAAAACATGTAGTTTCTGTCTTTTTTTTGGTTAATAAAAATTAATTTTGGACTTTTTTGTTAATAATAATTAATTTGGGACTTTTTCATAAACTGAGCCAAAATATTGGGACTTTAATGGAGATTTCCCTAAAAAATCCATAGATATTTATGTTTTTTCCCATTTCAGCCATTATTGACCTACAAAAACCGGTTGCCAGGTTTGATTTGCTGACATGGACAAATTTTTAATGAGGTGGCAAGAGAAATGTCCATATTAAGTGTGGTAAAGTGTCAAAAAGGGATGTCTTTTAGTGTTTCCATCGGTATAACACATGATTATGTGGACATTTTCCTTGCCACATCAGATGCCACCTCATTAAGTTTTTTCCATTGAAGCAAATTAAGCCTAATGGCTTTTGTGTAATTTGCAGGGATAATAATACAGTTGCCAAAAATTTTTATTTTATGTTATGTCTTGTCCTGTGTAAAATGGGGCGTTAATCAAGAATTTGCAAGATTATTTATTTATTTATTATTTTAAATTTCACGTATTTACAGGCTGTAATAAAGAGGCTTATGACGGATGTACCATTTGGTGTACTTCTTTCTGGAGGTCTTGATTCATCTCTTGTTGCTGCAGTTGCTTCACGCCACCTGGCAAATTCAGACGCTTCTTGGCAGTGGGGAACAAAGTTGCATACTTTTTGTATTGGATTGAAGGTTAATATTCTTTTATTATGCAGACGTTTCTTTAAAATTTTCTTTAATCTAATATTATCCAGTTAATTTCTTAATCTTTATTTTNNNNNNNNNNNNNNNNNNNNNNNNNNNNNNNNNNNNNNNNNNNNNNNNNNNNNNNNNNNNNNNNNNNNNNNNNNNNNNNNNNNNNNNNNNNNNNNNNNNNNNNNNNNNNNNNNNNNNNNNNNNNNNNNNNNNNNNNNNNNNNNNNNNNNNNNNNNNNNNNNNNNNNNNNNNNNNNNNNNNNNNNNNNNNNNNNNNNNNNNNNNNNNNNNNNNNNNNNNNNNNNNNNNNATTTTAAAAAAAAAAAAAAAAACTAATTTGATCAATTGTGAACCCATACCATCCATGTCAAGTTTTTAGGTTTATACGATCAAATTTCATTGTAAATTTTAAAATAAACCTAAGAGTAAATTACAGTTTTGGTCCCTGTGGTTTACCTCTTTTTTCAGTTTAGGTCCAAGTTTTGAAATTTCACATTCAAGGTCCCTATGGTTGCATAAAATTAACAGTCATGGTCCCTGGTTCAGGTAAAAAGACCATATTACCCTTTTTAATTTCTTTTTCCTTTTTCTTTTAAACTATTGTTTATTCTTTTTCTTTTAATTACATAAAT

***CiASN2***

>scaffold102796

AATTTCTTTTTCCTTTTTCTTTTAAACTATTGTTTATTCTTTTTCTTTTAATTACATAAATAAACAATAGTTTAAAAGAAAAAGGAAAAAGAAATTATAAAGGGCAATATGGTCTTTTTATGTGGACCAGGGACCATGAGTGTTAACTTTATGCAACCATAAGGACCTTGAATGTGAAATTTCAAAACTTGGACCAAAACTGAAAAAATAAGTAAACCATAGGGACCAAAACTGTAATTTACTCTAAACCTAATTTGATCATCAAGTTCTTGGCTTTTTATGATATTTGCCAGTAAAATAAAGTAATAAATAAAAATAAAAAAATGAGGGATCCCGGAAATGTTTCTTCCTGAATATGTGCTTTAATTTTTCTTTTTTAGGGTTCTCCTGATTTGAGTGCTGCCAGAGAGGTAGCTGATTATCTTGGCACTCGCCACCATGAGTTTTACTTTACTGTCCAGGTAATTCTTTTTACAGCTAAATTCAAGAAATAGCAATGTACTTTTCTGATTTTATGTATTATAGCATCCTACTTTCATTTCTTCCTATTACAGCTTTATACTTAAAAATTTCCACTCAGTTATAGGAAAATACGATTAGTATACAAAGCAATTTTCACTGCTGTAACCATGTAAGATTTTTTA+AAGTATAATATCCTACAAATTTACAAGTACAATGCTAAATTTGGTAGATTTTTCAAAATGCAATGCTAAATTTAAGAAAAGATTGAGGTAATAATAATAAACACTGTAGTAGGATGCTATAATACACAAATAAATGAAAGAATTAGAAACTTGCTAATATTGATTTGTTTTTTTGTAGCAACCTACTTTCATTTCTTTCTATTACATCATTATACTTAAAAGACCACCCGATTATAGCAATTTCATTGGGTAGATTTTTCAAGGGAAATCTCCAACAAAGTCCCATTATTTTGGGCTAATTTATGAAAAAGTCCCAAAATTAATTTNNNNNNNNNNNNNNNNNNNNNNNNNNNNNNNNNNNNNNNNNNNNNNNNNNNNNNNNNNNNNNNNNNNNNNNNNNNNNNNNNNNNNNNNNNNNNNNNNNNNNNNNNNNNNNNNNNNNNNNNNNNNNNNNNNNNNNNNNNNNNNNNNNNNNNNNNNNNNNNNNNNNNNNNNNNNNNNNNNNNNNNNNNNNNNNNNNNNNNNNNNNNNNNNNNNNNNNNNNNNNNNNNNNNNNNNNNNNNNNNNNNNNNNNNNNNNNNNNNNNNNNNNNNNNNNNNNNNNNNNNNNNNNNNNNGTTTATGAAAAAGTCCCAATGTTGAATTTTTTTACACAAAATAATAGGATTTTTTCATAAACATACCCAGAATAATTGGACTTTGCTGGAGATTTCCAATTTAGGAACCGGAAGAAAGGGCTAAACTGAATATGTTAGCCGGTATGTTGTCCTTTTCTGTAAACAAAATTAATTTTGGGACTTTTTCATAAACTAGCCCAAAATAATGGGGACTTTATTGGAGATTTCCCATTTTTCAAAGTAGAATGTCCTGCTAAGAAAAGAAGGATATATTTGGGTGGATTTTTAAAAATAAGAAAAATAAATCGAAAGTAAAATGTTGTAATAGAAGAAAATGGAAGTAGGATGCTATAATACACAAATAAATGAAAGTACGTTGCTACTTATTGCATTTTTGAAATACTGATTTCCTTAAGTTTATTAGGAAGGAATTGATGCATTGGAAGAGGTGATTTACCATATTGAAACGTATGATGTGACAACTATTAGAGCCAGCACACCAATGTTTCTCATGTCGCGTAAAATCAAGTCTTTGGGAGTAAAAATGGTTCTTTCTGGAGAAGGTTCTGATGAAATCTTTGGTGGCTACTTGTATTTCCATAAGGCACCCAACAAGGAGGAGTTCCATGAGGAAACATGCCGAAAGGTTCTAAATAAACTATTATTATTTATTAATATTTAGGGTTA

***CiASN2***

>scaffold17556

AAATGAAAATTTGATGGTTAAACCAGAAAAAGTGCAAAAATATAGTGTTTTTTTCGTAATTTGTCCTAATGAATAAAGTTATTTCTTGAATTTAACCCTTATTATTATTGTTATTAATATATTTATTCATTATTTATAATGTTTGTTGTTTTGATGCAGATAAAGGCTCTTCATTTGTTCGATTGCTTGAGAGCAAACAAATCTACTTCAGCATGGGGTCTTGAAGCGCGTGTACCTTTTCTAGATAAGGCGTTCATTGATGTTGCAATGAGCATCGACCCAAAATGGAAAATGGTAAACTAATATAACCCATTTTTTATCTATGGAATTTTAGGGAAGGGATGTTTTTGAATGGAATAAAAAGTTGAAAGTTTTAACCTATTTTTATCTACAAAATTATTTTGTCACAAATCTTATAAGAATTTTATGATTTTGGTTAAATAAACTATTGTTTTATTTTTCAAGATAACTAGAAAGTATTCTTTTTTAATTGGGTTTTACTATGTTATTTAGTTTTTGTAAATAATAAATTCTAGAGTAAATTACAGTTTTGGTCCTTGTGATTTACACCAACTTACAGTTTTGGTCCTAGTTTTNNNNNNNNNNNNNNNNNNNNNNNNNNNNNNNNNNNNNNNNNNNNNNNNNNNNNNNNNNNNNNNNNNNNNNNNNNNNNNNNNNNNNNNNNNNNNNNNNNNNNNNNNNNNNNNNNNNNNNNNNNNNNNNNNNNNNNNNNNNNNNNNNNNNNNNNNNNNNNNNNNNNNNNNNNNNNNNNNNNNNNNNNNNNNNNNNNNNNNNNNNNNNNNNNNNNNNNNCAGATCTTTCATTTTTGGTCCATGTGGTTTACACTAATGGTAACAGAGGGAAGGACCAAAAGTGAAAGATTTGGAAACCATAGGGACCACGAGTGAAAGATTTAGAAACCACCGGGGCCAAAAATGTAAGATTTTCAAAACTAGGACCAAAACTGTATTTTGTGTAAACCACAGGGACCAAAACTGTAATTTACTCTAAATTCTATATATACTTAAATAATATTTAGAATAAGTGTTGTAAATATAACCTGTATTTGCATTAAGTCTAAAAAGCTAATATGAAAATTAATTCTATTCCATTAACCTTGCTTATGCCATTACCTTGGTCATTCCATCCAACCAAAGCATAACAATATGTGCTACAATATAAGTTTTTAGATTTTTTATTTGATTCTGTAGGCTTTCATGCAAATGTTATAGCATTAGAGGTGGTTCTTGAATATAATAAAGCTAAAATGCAAGCAGAATATGGGCATAATTTACTGCTCAAGTATTTTTCAAAGATTTAATTTGTTGTAAAAATTGATAATAGATTCAAAAAGACATTGGGAGGATTGAGAAGTGGATTTTGCGCAATGCATTTGATGATGAAGAAAACCCTTATCTCCCAAAGGTATGCTGTTTCCTTTATTAAATTCATTATGAAAATATTTAAAAATATATATAAAAATTTCTCGTCTCTTCTCCAGCACATACTGTACAGGCAAAAGGAACAGTTCAGTGATGGAGTCGGATACAGTTGGATCGATGGTTTAAAGGATCATGCAAACCAACAGGTCTCTTGCATTTTTTCTATTTTTAGACAGGAAGGGTAAAACGGTCATTTACTCCCCTTCAGTCCAAGAAAGAAACAACTTATTCATCGTTCAGCACTTTAACAGGCTCTAAGACTATTTATTGGTGTGCAGGTTACTGATTCAATGCTAACTAATGCAAATTTCGTTTATCCCGAAAATACCCCTACGACAAAAGAAGCATACTACTATCGGACAATCTTTGAGAAATTCTTCCCCAAGGTTAGCTGAGTGGACTTTTTTTTTTTCTCCTTTGTGATACCCTGTTTAAAAGACGTAAATACCCTCATCATCAGAGTGGCATCTTTAGCATGACCATAAACCCCGTAAAAAAGACGTGAATGCCCTCGTCGTTGAAAACAGACCACATATTTCTGAGCATTTTATGACCCAACTATTATTCATCAGCCTGTATCATGTAATTCTTTAGACAAAACTTCATAAATGGTCCTTGTGGTTTCACAGGATATCAACTATAGTCCCTGTGGTTTGAGAACTTTTCATAAATGGTCCCTCCGTCAGATGAGGGCAAAATGGTCTTTTCCATATTTTGCCCTCATCCGTTATCTATTTAATGGCCAAAAGTTGACGGAGGGACCATTTATGAAAAGTTCTCAAACCACGGGGACTATCCTTGAAGTTTTTGAAAGTTAGGGACTAAAGTTGATATCCTGTAAAACCACGGGGACCATTTATGAAGTTTTCTCAATTCTTTATTTGGCATTTGAAACAGAATGCAGCACGGTTAACGGTGCCAGGTGGTCCAAGTGTGGCATGCAGCACGGCTAAGGCCGTGGAGTGGGACGCATCGTGGTCCAAGAATCTCGACCCATCTGGTCGAGCCGCTCTCGGGGTGCACTCAGCCGCCTACACCAACAACAAGATCGATGCTTCCACTTAATAAGCCCAGTATTTATTTATGAACCGATGCTTGTATTTCAATCTTTCTTTTAACGATTTTTTGGTTCATTTAGGGTTGGAAAGCGCTTGTTTCATAATCTTTGGGTTGTTAAACTTCGTTTCTGTGGATTCGATTTACTTTCGACTTGTGTTTTGATAAAGTTTTCTTGTATAATAATCGGTTTGTTTTTAGTTTTCATTTTTATGTTTCATATAAATGCTTGATGCTTATAGGGCTTTTTATTGCCATTATTAGAATAAAATTGTACGTATGTTGTTTTCATTGATAAACAAGAAAGTATTATTGCTTATGTTAGTAGCGAAATTAGAATACATTATGATTTTCTATAATTTGCCCGTGATTAAGAGGCTGTGTTTGATAGCCTCTTAATCTGAATGATTAAGAGGGTCTGAATGGGTTCAGACTGTTTGATAGAATGAATCTGAATAAGCTCTAAAGAGTGTTTGATAGAATAAATCTGAATAAGCTCTAAATTGTTCAGGCCTTTGAATGATTCAGATCTGAATGGTTCAGATACTACCAAACACAACCTTAATTAATAAACTAGAAACGATTGTTAAATCCATTAAGTCAAAAACAAACACGATGTATAAAACTTATCAGATTAAGTCCTATTCAGCTATTCTCATTAGATCCATTAAGTCAAAAAACCGTATCCTCTACAGCTTTCATTTGGGAATAAAGCTGAACAATAACCAAACGAATCTCACTAATCCCCAAATGAGACACATCCCCTCCCGCAAAATAATGCACCGTGTCCCCAATCTCCATCCAACTAAACCCCGTTTGCTTCTTCACTTCATTTTCTTTCATCAAACATCTAACTCGAGTCACTTGTCTCCACAAACCCAACGAAGCATAAATATTCGACAACATAACATACGCCCCCGAGTTTTTAGGGTCCGAACTAGTAATTTTCTCAAAAGCTAATTCCGCTAGCTCGGTATTCATGCTAAAACGACAGCCCGCAAGAATGGCACCCCACACACCCGAATCGGGTTCAAATGGCATCTCTTGGATCATTTTATAAGCTTTTTTAACTTCACCGGATCGACCCAAGATATCGATCAAACAAGCGTAGTGTTCGGACTCGGGTCTCACTTTGTAAAAAGTTTTCATGGAGTTGAACCAGTATAAACTCTCATTCACCTTACCTGCATCATGAATAGCAGTAAAGCAAATTATATATTAATATAAAACGAATACCAATATAAACTCTCCTTCACCCAAAAACAAAACAAGAAATTTCTAAAACAAAATTTTGTGTACCCGCATGACCGCAAGCGGAAAGCAAGCTAAGAAACGTGATCCCATCCGGTTCAATTCCCTTTAATTCCATCTTCTTAAAAAATCCAAGAGCTTTCTCATAGACTCCATGTTGTGCAAAAGCCGCAATAATTGTATTCCAAGACACAATATTTGGAGTTTCAATACATTCAAAAGCCGACTCGGAATCAACAAGACACCCACATTTGCTATACATTGTGATCAAAGCATTTCCTATGGAAACATGTGCACCAAATCTAAGTTTGAACACAAGTGCATGAATTTGTTTCCCTTCAGTTAATGATGCAAGATTTGAACATGCAGAAAGACCGGAAACAAGAGTGTATTGATCGGGTCTTAATGGAGTATGGATCATATGAATGAGTAGTCTTAATGCTTCCTCACCTCTTCCATTTTGAGTATAGCCTACCATGGAATCAAGAATTAGCAGTCTTGATAGAATGTTTGTCACCCATAATCTCAAATAAAGTCCCATTATTTTGTCATTTTATGAAAAAGTCCCAAAATTATTATTTTTATATACAAAAAGAACAGAATACCGGCTAACATATTCATTTTAGCCCTTTCATCCGGTTTACCAGGTTTCCTAGATTGGAAATTTTCAGTAAAGTCCCACTATTTTTGGTCAGGTTATGAAAAAGTCCCGATGTTGAATTTTTTTAGGGAAATCTCCAATAAAGTCCCAATATTTTGGCTCATTTTATGAAAAAGTCCCAAATTAATTTTTATTAACAAAAAAGTCCCAAATTAATTTTTATTNNNNNNNNNNNNNNNNNNNNNNNNNNNNNNNNNNNNNGTCCCAAATTAATTTTTATTAACAAAAAAGTCCCAAATTAATTTTTATTAACAAAAAAGGACAAAAAAAGGGGCTAATGTTGGGACTTTTTTGTAAACTATGCAAAAATATTAGGACTTTATTGAATATAAAACATGTAGTTTCTATCCTTTTTTGTTAATAAAAATTAATTTTGGACTTTTTGTTAATAAAAATTAATTTGGGACTTTTTAATAAACTGAGCCAAAATTTTGGGACTTTAATGGAGATTTCCCAATTTTTTTACACAAAATAATAGGATTTTTTCATAAACAGACCCAAAGTAATTGGACATTATTGGAGATTTCCAATTTACGAAACCTGCTAAACCGTAAGAAAGGGCTAAACTAAATACGTTAGTCGGTATGTTGTCCTTTTTTAGTAAATAAAATTAATTTTGTGACTTTTTCATAAATTGGCTCAAAATAATGGGACTTTATTCCAATTTCCCTTGACATTATTGTGTAAAGTAAACATACCTGTTATCATAGCATTAAAAGAAACATCATCTCTACATTGGATTTCTTCAAACAATTCTCTGGCATCTTCAATCTTTCCTTCCTTGCAGTAACCAGTAATCATAGCTGTTATAGCAACTACATTCTTCCATGGCATTTGACCAAATAGTTCTCTAGCCTTATCAAATTCATCATTTTCCACATACGCACTGATCATTACAGTCCAAGAAACTTCATTACGACATGGCATGGAATCAAATAGCTCCCTAGCTTTCTTAATTTCCCCATTCTGTGCATATCCGTCAATCATAGCTGTATAAGATATCACATTTCTGTTTGACATTTGTTCAAAAAGCTCAACAGCTTTTTCGAACCTATTATGCCTGATAAAGCCCGAAATCATAGCATTCCAAGACGCTGTGTTTCTCTCTGGCATCTCGCTGAAGTACGCAAATGCAAGATCTATCATGTTATTATCGATGCATCCAGCTATCATCGAGTTCCAAGAGACTACGTTTCTCAGAGGCATTGAATCAAACAGTCTCTTCGATTCTCCAAATGATCCATTCTTCCAGTAGCCGGTGACCATCGCGTTCCATGAAACAACATCTCGTGTAGGCATTTCGTCGAACACTTTACGGGCAAGGTCCATTTTCCCGGCTCGAGAGAGGTTGCCGATTCTGACATTAGAGGTGTACACATCTCGAAGTGAGTGAAAGTATAGAATGTTTGTCAGAAAATGGCGACTGACGCGATGGTCAGATGAATTAATGGAAGAGAGATTTTGAAGTATTGCCCGAGGAAGGCGCGAGACAAGCATACAGGTATTGGGTGAAACTTGGTGCACATTTCACAGTGGAGATGATAAAGTGATGGGAGTCGTTTACCGACTTTATGTGAGATTGCAAGTTAAAATTGATAATGTTATTTGATTCTAGAGAAACTAAGAAAGTATTTTATGAAAAAAAAAACTATTATATTTTATTATCTTTGTCAGGAAAAATACAATTCATTATAATTTCCCTTTAAAAAAAATAAAAGAAATATCATAAAAAGATAACTAATAATATCAAATTGTTTTCTAAGTATTGACGAGACTTTTTATTTTTGGTCGATTACACTATTGGAGACGTTTTTTACCGGGTATCACCGGTTACTTTTCCATGTGGACTCCATGTCAGATTAAACCCGATTAAACCATTCAATGTTCAACCCGTTTTTCCTCAAACACAACCCAGTTCGATGTCAACCACTTTTTCCTCAAACGAAATCGATTGAACCCGATTAATCGTATAATCCTATATTCCACATCCATTGATTCTGTGAAACGAAATCGATTAAACCCTTTGTTGTTCAACCCTCTGCTATCTGTTCTACATAATGATTGAGGCTATTCCACTAGAAGTTACGATTGTGAAAAGCGATTACACCATATGTGTAATTCTGACAAGCAATTTAAGAAAAGATAAGTTCTTTTTGTTGGATTTCTATTTTACAGTTGCTCAGTTCAGTTAATTGATTTTGGTCTGATTTTGGTTTGTGGAGAAGCATGCCATTGATTTTAGTTATGATTAATCGATTAAACCAAATTAATTCTATAATTGTATGAGGTTTTGTTTTATATGAGGTTTAGTTTTTTATTTCAGGATGTCCTTAAACTAGGAGTTCCTAATACTATGCAAGTGTGTTTGTGTAAAAACCATTAACCTCCGTGATGGTGTGATGATCTTTATGAATGATATTGACATTGCAATAATGGATTTTAAGGAGTTTGAATCTGTTCTTCAAAGGTTGACTGGAGATAGATGCAAAAACCTTTACTTTTGTTATCCCAAAAGCTCTTTAACTAATGGTTTATGAATGTTAAAGAGTGATGAAGACTATGCAGAATTTATTGATAAAGGACATGAAACTGGTAAAAGTGATGTATATGTTGACTACTTCTATGAAAACTTAAATGATTGAATTGATGAAAAGGCATCACAGGAGGTTCATGAAGTAGAAGTTAAAGCTGATTATGATGCTATTTCACGTGATTCGCTTGAATATGAAGGAATTGCCTTGGAGAACATGTGTACTGATCCGTTTTTAACCGAGCTTTGTACAAATACAAAGCAGGCTCAAGTTGAAGAAGAAAAATCTCAAAGTGAAGATGATAAAGTTCGTTTGAAAGAAGAAATGGAAGACTTGGAAGGTCATCTAGGCGTACATTACCCAAGACATGATCTATCACAATCATGGCAATTAATGAACCCTATTTTGTATATGAAATTTGAAAGTCCATAATAGTTAAAGCAATCTTTATGCAATATGTCATTGCTAACGGTTATCATTTGTGGTTCGGAAAAAAGACTATGTAACGACCAAAAATTCATACTAGAAAATTTTTCTTTTCAATTAAAATAAAACATCCAGTTCCATTCATTCATTCACCAATAAAATAAAAGTGATCACATAGTTTGCAAATCCAAATAGTCATAACAAGATAACAGCGGAAATCATGATAAACATATGCCCGTGCCGTGTAGTCACGCCATGACTGCACCGCCAGTCCCGTGAATTAAACTTAAGACTGTAAGCACATCGCTTAGTGAGTTCCCCAGATTCCCATATAATTTATAAATAAGACAAAGTCTCCAGGCCTTCCCCAGACAACCAATAATCATCTCATTCTTTGACTTTCAGGCTCGCCTCGATCCGTCGGTAATCAACTCATTCTTTGACTCTCGGGCTTACCCTGATCCATCAGTAATCAACTCATTCTTTATATCATAACTGTTGGATTAGGTGTCTAAGCCATGACTAATAATTTGTATTTACTTAATTGATAGTAGCAAAGTTCATTTGGGTTGCCCTCATAGCTAGAGGTTTGGCAGAAATTTCAGGAGAGAAAAAGATTGATTTATTATATGATTAATAAATCAATGGAAATAAATTAAATAGTTTATTAATATATTATGGTGATAATATATTAATTAGAAATTATATTA

*CiGLN1;1*

>scaffold641

ACCTATGTTTTAAATGAAAAATTTTTCAGGGGTGTTTTGTTTTAAAAGCCGGGTGTTACACTAATCCATCCAAGTTGACCCAAAAAGTCAACTGGGCAAAGTCAACGTTCAATGGTCAAAATCAAGTTAGCATTGCTTTCTCCTCGAGATAGTGTCGCTAACTCTACAATCATATTGGGCCATGACGACAGTTAGCGTTGCTATCTCCATAAGATAGCGACGCTAACTTTGAACTTTTCCAAAACTTCATTAAGCACTTAATTTATTAAGATAAGTGCTAATACTCTACAAAACTCGTCCATTAGGTCTGAAACTTCATAAAAGTCCAAACTTTCTTGATATGCATGCCCCATGCATGTCAAATCTTATACTTAGGTCAAAAAGTGAATCAAACTCATGCTTAATCACTTGATGACCAACATTCACATTCCTAATCTGATCCAGGAGAGAAAACAAACTAATGCAAGGGACCAAAAGGCCTAGAGTTTCCCCAACTCCAAAACCAAACCAAAACAAAAGCCAAACTCCTAATCATTTAACTTAATCAATAATATAATCTAATCTAAAGCACAAATCAGGTAAGAACAGCTTACAAAAGTCCAGATATGAAGTTTGATGTGGATTCCAAAGCTGAAAAGGATACCCAAGCTTCACTCCTATAATCCTCCTTCTTCAAGAATGATCCTCAAGCAAGCTTTAAACCAAAAACCCTCTCTCAAGCCTCTAAGAACGATTTCAGAGCTGCCAAGGGTGAAATGAGTGGAATGACGACCGCTAGGGCTACCAAAGCTTATATATGGGGCCTAAGGCCCAAGAAACTTAGGGTTTTGTGCAATAGGTGGTTAGCGGCGCTATCTTCTCCCGATAGTGTCGCTATCGAGGACCAATCTGCGATTCAAAAGCCGACTTAGCATCGTTATACCTAAAGATAACAACGCTAACTCACGATCCAACATTGCCCAAATCCTTTTAACTCTATACTATGTCATTTCGGGAACCGGGTATTACCATGTAACAACAATGGTTATTGGTTTCCTTTGCTTCAACTTCTCTACATCAACCGCTACAAAGGGATTTGGCTCATTCTTTACAAGCAAACACAACTTTCCTTGCTTTATCTTATTGTCCACTTCTTTCCCGTCTTCGTCACCTGAATACCATACTTCAACATCTCTCTTGGTTTGAACTACATATGCCTTCTCAACCATCTGCTTTCCAAATACTTTCTAGCCTATCAAGAAAGTACACCTCATTCATCACATTTTATTTCTTTGGGGTTAACTCTCATTCTTAAAGAGGTTAAGTCCTTGATAATTATGGCAGGTGATGGGTGTTCCATGAGAGGTTGGTCTTTGAGTTTTGCTTCAAAATCTAGATTAGTAAGTTAACTTGTAGTTAGCTATTTTTGATGGGGTGGTTAGGTTTCGATGTTTTAAAGTAAGTATTGGGGTTGCGAGTTAAACTTTGGTGGGTTGTAGTATGGTGATACTAGTTGTATGCTTTTTGATAGTGAGAAGATCTTAGGTGTAATGGTACTAGAATGTTAAACTTCTGGCTTATAAGACCAAGAGCTTATCGGTATGAACTTTTTTCCTTATCTTATTCATTTCAACCCAGATCGCGAGGGATCCACTAAAGGTAGATGTCAGTGAGGTTTTGGGCCAGAGTCAGATAACCAAGAGTATGATGACAGGACTCTTGTAGTGCAGTTTGTGCAGCAACAAGGGCTAAAATGCCTCCTAACTCTTGAGAAGTTATCATAATCATAAATGATTGAGCTTGTAATTCTCCGTAGAGATCACAGAGAAACATTCTTTTAAGATTATCATATCCTTTAATGATCATTTTAGCGGTTATCTAACTTTTACCAAGTCTATTTAGAAATTAAAGATTAATTTCTAGGTTGGATTTGATGGTTCCAACATTTTACATTTGAGTCGTGATGAGAATAAATCTATTGAAGGAATTGTCAAGGGTTTCACCAAAAATAGAAGTGAAACTATTGAACTCATTTAGGTTTCAGGTAACTTTTTTTTCATTCGCTTCCTTAGTACCCTCTAATATCTTTTGTAACGTTATCTAAATTTGTTGTGTGGTAGTACAATTTTGAATTCGATAAAAAATATCACGAGGTATTACCACGTGAAAGTTAAGTGAAAGCAATTTGATCACTTTCCATTTTCTTGGGATCAAGAATCATATAGTCATAGGTCTTCCCATGATGGTGACATTGTTGGTCAGACAAATAAAGATATTTTTATAGATGTTATGATGGTGGGACAGAATTATGGTACGTTTTTAACCTATAAATGCTCTTCAATTAAGAATTAGTATAATAGTGGTACGAGTCGAACCCTAAGAGTTGTGAATGAGTAATCTCTTGCCGGGTTTGCATACATCTAGAGATGAGACCAAAATTGTAAAGTTCAANNNNNNNNNNNNNNNNNNNNNNNNNNNNNNNNNNNNNNNNNNNNNNNNNNNNNNNNNNNNNNNNNNNNNNNNNNNNNNNNNNNNNNNNNNNNNNNNNNNNNNNNNNNNNNNNNNNNNNNNNNNNNNNNNNNNNNNNNNNNNNNNNNNNNNNNCATGTATCTTGAAATGCAGGAAATAATAAGAACCAGATTACAATAATATATGTTTAATAAACATATATGCAACTACTATATTTTATATTTTGTTCTTTTATCTTATGGTTTAAATTTTAAAGCACGTCCAACTTTGGCATAAACACCTTTTTTTGGTTTCATGCAATGTTCTAAGATGTAGATCCTATATGAGATTGTTTTCCATTTTGCAATATCGTGATTATAAATTTTAATCGGGATAGCAAGATCCTACCAAAAACATTTTAAAACATAAAAAATTCATTTCCATTAAGTAGTTAACCATTCAACTAATGACAAAAAATTTAAAACATAAAAATTACATCATTAATGAATAAACAAGTTAAAATAGGTTTTATAGGATTCATATTCGATTCACTTATTTAATCAAAGACAAAATATTGTTATTTTTTAAATAAACGAAAAACAATTTTATGATCGATCCTATCAAGGTTGTTAAAATCGCGATCCTGCTCGTAGGATCGTACGATCCTACGATCCCACCTAGGTAAATCTATCAATGATCATACGTGAGTCATTTTTTAGTAGGATCAATGTAGGATTGTAGGATCGAATCGTAAGATCATAGGATCAATCATAAGATCATAAGATTGTTCAAGATTGTTCCCATTTATTTATTATTTTTAAAATATAATTTTTTTGCTTTAGGCTAACTAAGTGAATTAAATATGAATCCTAAAAACTCATTTTAAATTGTTTATTGTTAAATGATGTATTAGAAATATATATTTTTACATTTTAAACTATTTAGTATTAATTAAATGGTTAATTGATCGATGAAAATGTGCTTTTTATGTTTTAAAATGATTTTGGTAAGTTATTACGATCCCAATTTGATTTCATGATCATGATCTTGCGAATCAGAAAACAATCCTATATAGAATTTGGATCATGATAACCTTGGATCCTATGATCCTACTAAAAAACAATCCAACTATGATCGGAATTGATTAACATGTGGGATCGTAATTTCAACCATTATATTTAGCATATTGTCAAAAGTAAACTATAATAGAAGCGTGGTGTCGTACTTGATAAACCATGCCTTTTTGCATTTTCTTAAATTTTGAATTGAATTAAAATATAACTTCGTATCATCATCATCATCGTCATCAAGAACATCAAACTGCACTTATAAAAACGCGCACCTGCCCTCCCCATTTTTATCGATTATCACCCCTCTTATCTTCTGCAACTTGCACTCCGATCTCATGGCTCTCCTTACCGATCTCGTCAACCTCGACCTTTCTAGTATCACCGATAAGATTATCGCCGAATACATATGGTTAGCACCTTCATTTCTTCATTTCCGAAAAAATAAAAAACTATTATTCTTCACTCATGAATCCCCTGTTTTCCGCCATGAATCTTTCACACTTCTTAAATTCTTCACAAGTTTCCCTTTTGTGCTTTGAATATCCGAACTAAGTTTTGATTATTTATGGGGTTTAGTAGCAGATAAATCTTTGATTCCGAATAGGATTATGCTGGATTTTGAATTTGTATCTACGGTTTTGTGGTCGATTTAGTTACGTATCCTCATTTATCACCGTATATGCTCATGGTCCATTGAAACCATTTGAACCAATGGAAATGATCTCGCCTTCTTCGATCTACAAGATTGATTTGATGATTGGTTGGTTCGAATCTGTACAAACAAAGCTACTAGTGAAAACGATTGATCTGGTGTTCTTTTCCTCTCTTGTACTTGATCTCGTAAACCACGAATGGAAAGTTAGAACTTTTTCGTCATTTACCGAATGTGTCAAAGAATCTAAAAGTAATTTGATCTAAAGGCTCTAACCCCATTAATCACGTCACTTAACTTGTTGAATGTTTAGACTTTTAGACTTCTTTTAGCTATTTTGACCACCGTCGATTTTTCTTTTTGTTTCTTTACTAATCTCAAATGACTGATCTTCAAACAGGATCGGTGGATCGGGTATGGATCTTAGAAGCAAAGCAAGAGTAAGTATTCATTCATTTGGTTGAGTTCAAGCTTTCGTTTCTTCTTTAAATTCATTCGATTTAACATTTTGGGTTTTTTTTAGACTCTTGAAGGACCCGTATCGGATCCCAAGAAGCTACCGAAGTGGAACTACGATGGCTCTAGCACAGGGCAAGCTCCCGGCGAGGATAGTGAAGTCATTTTATAGTGCGTAGTCGTCCCCTTTAATTTTTAACTTCTAATCTTTGAATTTTTGTTGTTTCTTGAATTAAAAATGTGTTCTTTATGAAATTTAAACAGCCCGCAAGCAATCTTCAGAGATCCATTCAGGAGAGGCAACCATATCTTGGTACATTTTCACATTTTCAAGATCCATTCTTTTTGGGATTTATATTTAGTGTTAGATCTTGAAGTTTGCAACTTTCTTACCAAATTTCTATACATTTACGTAATGTAGGTGATGTGCGACGCTTACACCCCCGCCGGCGAGCCAATCCCGACGAACAAGAGGGCCGCCGCCGCCAAAATCTTCAGCAACCCAGAAGTCGAAAAGGAGGTTACCTGGTATATTACCTTTCCACCCCTCAAACTTTATAAAAATTACAACTTTCTGACCAAATTTCTGTATCATTTTTCGCCGTTGATAGGTACGGAATTGAGCAAGAATACACCTTGTTGCAAAAGGACACCAATTGGCCGGTGGGCTGGCCTCTAGGCGGCTTCCCTGGTCCACAGGGGCCATACTACTGCGGTATCGGTGCTGACAAGGCTTTCGGACGTGACATTGTTGACGCACACTACAAAGCTTGCCTTTACGCCGGCGTCAACATTAGCGGAATCAATGGAGAAGTTATGCCTGGACAGGTAAAATTTTGAATCTTCTAAACAAAACCCATCTCTCTGTTTTCCTGTTTTTAGACTTTAAACCGATAAAAGTCAAATTCTTTGATGTTCGTATGCAGTGGGAATTCCAAGTAGGACCTTCCGTCGGCATTTCCGCCGGTGATGAATTGTGGGTCGCTCGTTACATTCTTGAGGTACCTTAAAAATCTTGACTTTTTCGTTTTCTTCACGGACTTTAAAGTCGACGTCAAAAATTGAATCTTTTTCTTTCTTTTTTTCTTTAGAGGATCACGGAGATTTATGGGGTGGTTGTTTCATTTGACCCCAAGCCTATTCCGGTTAGTTCTCTTTACATATTTTTTGATTTCTAATCAAAACTACGACGTAAGTGTCAAGTATAAAAGATCGTTTTTTTAACAGGGTGACTGGAACGGGGCTGGTGCTCACACAAATTACAGGTATTTTATGACAAGACGATTATTAATATCTATTTCAATGGTGGGGTTGTCTTTATTTTAACTAGAATCATAATCTTTCAGCACAAAATCCATGAGGGAAGAAGGAGGATACGAGATTATCAAGAAAGCTATTGAGAAGATGGGTTTGAGGCACAAAGAACACATTGCCGCATATGGTGAAGGCAATGAACGTCGTCTCACTGGTCGCCATGAAACAGCTGACATCAACACCTTTTTATGGGTACTACGTCATGTCACATAAATAAAATCTTATTATATACACATTTTGTTACAAAAAGATTAATTTTGTGGTTTGAATATTATTTGTGTAGGGTGTTGCAAACCGTGGGGCATCTATTCGTGTTGGAAGGGACACTGAGAAAGAAGGGAAGGGGTACTTTGAGGACCGAAGGCCAGCTTCAAACATGGATCCATATGTGGTGACCTCCATGATTGCCGAGACCACTATCTTGTGGAACAAATCTTGAAAAGAGATTGAAATTATGAAGCCTGATTGGGAGGGAGATTTGAAAAAATGAATTAGAATTGGGAACAACCCTTGTGTTTATTTGCATGAAAATTAGCTATTTTCTTGTTGTTTGTGTTTCATCTTTGATTATGGTAGTTTTGTATTCTATGTTTCTGGTAGCAATGCATTTTGTGTCGAAAATGCTTATAAGCTACTAGTTGATGATTTGAGAATTCTCCCATGGTGTTTCTTGGGAATATTGTTGATATTGTCTATTCTAATAACAAATATGGAGGGTTGGTTTCGATCCCTTGTATCTTGCCTTGTTTTGCATTTGCTAGAACATCATTTCATGGATGTCACTATTATCAAGTATGCATGCATTACTATATTTTGGTAAAACTTTTTTGGAGTTTTTTAAGGGATTGTATTCAGTCAACTTAAATTGTTTTATCTAGATATTAATTAAATATGTATTTGAGGATTTCTATAAGTTTGTCATGTTTTGAGTTAGTTTATTGGGCATTTAACGAAAGAGTATCTTTTGTGTGTATGTGTTCTTGAGCGATAGGACAGCCGAATTGACGACTACAATGTGGTGACTATGTGTGCATGGTACAAGGGAAGTTAAATATCGTTTTGTCTTTATAATTTATCTTTGTGTTGTGTGCAAACAAGAAGCCGAATATGTGATTCTTTAGAGACAATTCGAGACTTTCTTTTTCATTCCAAAAATTTTTTGGGTCATCTTCTATCGATTTAATAGGATAACGACTTAATGGTGGACATGTGAACTTTAATTTTTTATTGTACTTGGAGTAGATATACTTTTTTTTTATATAGTTTGTGATTTGATCATTTAATATGTTGCTAAAGTTATTCCTTATGTTGTCGAGAAAAGAGCTTTGCGTTGTAAGAATTGGAGAGCACAAAGCAAGTGTGGGCCTCAATCCTCTCACCATCATAGTTCTATGTTTTATAAAGATGTATATGTGTTTTAATTTTTATGTTTAACATCTTTCTTTCTACCAACTCTATCGATAATGACAAAAAGTGAAATTATAATGTTAAATAATCATTGTAAGGTGCTTGGAGTTTAGTTTAAAGTCATTTCGGTAAGGTGGATACAGTATTTGGTTATTAGGATATATGGTTCTTGGTGATAGTAGCATTTGTATATATAAATTTGCTCTTCAGTTCATTAAGGTTTCAAAAATGATGCCATTAATATGAGCGGTGGATTTAGAAAAAATGATAATAATAATAACCAAAGTTCACTATATAATTAATATAATATAAATAAAACCAAAAATGTTATGACATAAGTAAAAAAATTAACACTTTCAACAGTAATAGAAACAAACGAAACGGTAAGAATGAATTAATAAACCATATATGATTCAATTCAAGAAAACGTAATTGATTAATGCAAATTACAAACACAAAATTTACAAGATTGAGGCATGACAAGAAGTATCCTAAAATATAAATTATACTGGTATGAATTTATCTTGAGAATACAACAATCAATTCAAGTGAATCTTGAATCTGTCAAATCGAACGTATACTTCATTGAAGAACACATGAAAACCTGCAACAAATGTCTGAGTTTTGTGTATATTTAGTTACTATGGAATCTCGTCATAAGTGTGGTATATATAGGCTCTAATATGGCAATTATTAATCATAACTGTAAAAATTGGTTAAAAACAAATTGGTATCATTAAATGAATTAAAATTGTTTTCAGGTTTAATATTCATAAAAATTTACAAGTTGGGGCTAGCCCACTTGGCTAACAAGGTGATTACTAACAAAGAGGTCATGGGTTCAAACCATAGCACTAACATAACAAGGAGTAGTTTAGGAGTAGGAGTAGGATAGATTTGCCGTTCAAAAAAAATATTCATAAAATTTAATTAATTAGTCAACAAGAGCCCAAGGGCCTAAGTCTTATTTCCTATCTTTAACCATTAATTATATTTAATCCAAGCCCAAAATTGGCTAGCCTAACACCACAATCCAAGCTCAAACCGTTTAGCCCAAATCTTTTCCAATTTAATGAGTAGAATAGAGGCTTATAAGGGGAATCCTTTTCTTTATTTGTTCACGTTACATAATCAAAGTTCCAAAACACCTACAATACCTCGCATTTGAATAGAAAAGAAGGAAACTTTGCCAAAACACCAACAATACCTGTATAGGGAAGTCCACATTACATAAATAAAGTTCCAAAACACCAATAATGCCTCACAAAACTTTGATATTCAAAGCAGGAATGCTATATATGACACATCTAAAAAGATGTTGTAAAAACTTGAACATTTTATAGTGAATAATAATCAGATTTATTTATTGGCAGTAAACTAGAGACTTGAACTAACCACGTTTGCAGTAGAACTAGACAACAAGCATCCCATATACCATATCAAAAACATAGGTAAAATCTTACCGTTTTGTTCATTTCGGCCGTGAATAATTCTTTTTTCATGAGAGCACTTAGCCATACCAGCTCTCAAAAATATGACCTCACATTAATTTCATATAGGTAATGCCAATATGGAACAATTCGTGGTATTTTACCATAATATGTTATTAGTATGATTAAGAATAGATTCATCTCTAACAAAAACACATACTAATAAATACCATAAGATGCACATACATGTGTGTTTATGCTTTTTGAGTCTTCTTATTGAATCCAGATCGCGGGATCTCCAATGTTGCTAGCATAGGTTTCCGTTATGTCATGACTCGTTGTGTTAGCTTTAAGACCATTCTCCTTGATACGAAGAGAACTCAAGGCTTGTTGTCATAGGATTAGCTATATTTTTCTTTGACTTTATATAAATAATGGAAATGATTCCATTTTTCAAACTTTTATTGTTTTATGTCTACGCCTAATATGTATTTGTAACGTCCCAAAATCTAGAGTAAAATTTTTCGTTTTAAAACGGTTCAAATACATGAAACGGGTTGTCAAATAAAAACCATTCAAATTAAAACTTGGTATTCAAACATATAAAATCGTATTTAACAAAACATTTGTAAACATTATTCAGCAGAAGACTTGATTATCAAAACCTCAGATGCGGTACAGTCGCGCCGTGTTCCAAGCTCAAGCCAGTAAATACCTGAAAAACAGTTTGGGTGAAAGAGTAAGCACAAAGCTTAGCGAGTTCCGAAAAATACCACACGATTCATCACATAAAACACAATCACAAAAAGGTCTCCGAGCTTGCCTCGAACAACATGGCTCTCGGGCTTGCCCCAATCCACCTTGGCTAAGAACTAACGAGCATGCTCCGTTCTGTCCGTCACTGAATCAACTAATTCTTTGGACTCTCGAGCTTGCCCCGATCCGTCCGATTATCAACTCAGTCTTTGACTCTCAGGCTTGCCCCGATCCGTCCACAATCACGTAAGGCACGTACAACACATTACTACACATAGTAATTGAGCTACCTATGCATGTAATTTCACCCATAGTATAGTGAGAAGACTCACCTGATCCGCCAAGCCGGACAACACTCCCCACACGTGTCCTCAGACCCGTCAAACAACTTCCTAACAAGGATAAACCTCACTTATAACTTCCTAAACCCTAAATAGGATAAACCTCTAAGACCGGTATCTCGAACCCTCAAGTTTGACCAAAATCAAACTGTCAAAAATTCAACGGTCAATGGTCAAAGACAAAGTCAACGGTCAACATCCAGTTAGCACCTCTATCTCCCCAAGATAGCGTCGCCAACTAAGAACAGGCTCAAACTTCATCGCGAACCATGATTGAGTTATCGCCGCTAAATCCTCTGTTCAACCAAAAATCCTCCAAAAGCTCTTAATCCAATAAGTCATGGTCTTAAACCTTCCAGGACCCTTTCAAACAGTCTCAAGGTCCATAAAAGTCCTAACTTTATCGATCAATCAAGGCCATGCAAGTTCATCAACAAGATCCAAGTCCAAAATGGTCCAAAACTCAACGAAACCACCCAACGAGAGCTTGACTTACTCTCTAAATCAACCTAATGGCTCCAAATTAGGACTTAAAGACATAAAGTTCCGAACTTTATCCTTCCCTACCAAGGACAACACCAACAAGGGATCTTTAATCCATTTTAATTCATAAAATCCCACTTAAACACAATAAGAACCTTAAAAATGCATAAAGGTGAGAACTTTACTCACAATGGTCCAGAAACAAGGAGCTTTAGCTGATTTACACTTGAAGAATGCCTTCTAACAACACAATGCACCACCACACCACCTCTTCTACTCCAAAAATCCTTCCAAAATGATCCAAAACACCAAAAGTCACACCAAAGGGGCTAGAGAAAGAATATGATGACCTTGGAAGCTGAAAAGTACCGAATGGCAGCAGCCCTAGGGTTCCCCACCCTTAAATATGACACATAGCTCAAAAACTAGGGTTTTGGCCCAAAATCAGTAGTTAGCGTCGCTAACTGAGATCGTGCAAAATCACTAACCAGAAGCACATTAGCGACGCTAAAAGCACATTAGTGGCGCTAACATCTTTTCTCCCAAAATTTGATTTCTTTTAACTCAAGACTTGACATTTCGGGATTTGGGTCTTACAGTATTGACTTACCATTATATATACAATTTTGTGTTTTAGTAAGAAAAACCATGCTATTACAATGAATACAAATGACAGTTTAAGGTTTAGGCCATATAAGATTAGATTCACTGAAGCTTTTAATCCACTCAGCTTCTTGAGCAGCTTTGGCAAAAGCTACAAATTTTGCGTCTATAGTAGGTATTCATAGTTTTCTTAAATGATTTCCAAGAAACAACAGCTCCACAAAGTTTACATAGAGCCAATTGCGGATATTGTTAGATTTACTGGGAATCCAATTTTCTTCACAATATCCTTCTTAAATAGGACGTGTAATGCAATCCATGGTTCATTGCAAGCTTCAAATATCTAAGCACCATTGGTAAACATATCGATGATTTCCCAAGGTTGTGAGAGTATCGACTCAATCTAATTACAATATATGCCAAGTCTAGTCTAGTGTAATTCATATAATGCATCAAGCTTCCAAGAACTTCAATATATTCTAGTTTTAAGCTACACAATCACCTAGATTTTTCTTAATATGGCTAGAATGATCAAATAGGGTAACAATCGGCTTGTCATAATAATAATAACATAGCTATCGAAGGTTCTACTTACTTTTATACCAAGCATTACATCGGCTACTCTCGCATCCTTCATGTCAAATTAGGAGTGTAACATTTTCTTGGTTTGGTTTATAACATTTAAATCAGTTCCTAAAATAAGCATATCGTCCACATAAGAGCATATGATGACATATCCCTTCTTATATTGTTTAACATAAAAACATTGATCAAACTCATTTATTCAAAATCTATTAGATCTTTGATCGGAAAAGTGTACGTATGGATTTTTTTATTTAGTATTAAAATTTACATGATACAATTTTTATATACTATGATATTAACATATGAGAATCAACCTTCTGACTCGTTAGATAATAAACGAATAATGAATAATCCAACATTTTACAGATAATACTTCTAACTAAGCCAATATGCTTAATAGACTGACACAAAGAATCCTACTGAACTATCTTCAAAAAGCTTTCAAACCTGTTATCTTCAACAACGTTTATGTTCAAATTGTATAGATCACTTTCAAAATCCATCCACAAATACGACGTCTCTTCATCACATATCACTTCTTCTAACTGTGACAATTCCCTCATGGACTCCGGCAACTCTTGTAAACCGTGGACACCGCTCATCTTAAGCACTCTCAAGCTGCATAGCTCCCCAATTTGTTCAGGAAGTAAAACCATACTCAAACAATCAGATATATCAAGGAAGCTAAGATTATGAAGACTTCCAATCGATTCTGGTAATTCTTGTAATTTTGTGCAGCAATGAAGCGTCAGAATTTGAAGGTTCGAAAGGTTCCCTAACTTTTTAGGAAGAGCATCAAGCTCATGGCAATTCGTGATGCTGAGATTCTGGAGATGAACAAGACTACTATATAGCCCTGAAGGTAGTTCCGTCAGATCATAACAACAATCGAATTCCAGATCTATGAGATTTGATAGAGTATAAGGAGAATCAGTGACGCTGTTTATCAAAGCATCGCCTATCTCACACATAACAAAGGAAAGTTTCTGAAGATTCGGCAATTTGAAGATTGGTTGAATAGAGGAAGAAAATGAAATATGTTCGAATCTGATGCTTGTTAGGTTAGCGAAGGAACCTATTAATGGAAGATTGAGAAGTTGACCAGGATAAACACTGCAGGTAGTCGAATATCCACCCGCAGACAAGGTAACAACTGAATTGTCTGCCAAACCGTAAATACTAATTGAATTCGTAATACTATTGATCATCGTTTTCGCCTGCACTTGAATCTGGAAGAACGTCAAGAGCTCGTTGTTCAATCGAATCAGCTTATTTGAATGAAGAATTTTGTTGTAAATAGTCCGATACTTGATACTCGAGCACTTGAGAACGATCTCTTTAGCATTACACAGATAACAGATGAACGTTAATGTCTCTTTCTCAGGACGATCCAGCACTTTGCTTAACCTCCAGCTTTCATATAATACAGGCTCGATCCTTTTTAGGGTATCTTCAAGGCGTTTGAGAAGGGTTTTGAATCTGGTAGTTTGTATAATCTGCCCTATAACTGTTTTCTGTAATTCACCCAATACAAGCTCTAAAGCAGTTTCAAATACTACCGCCATTGATAATCAGACAAAGTAAGTTGAAACCCCGATTAAGATTTTATGTGAAGTGAAATCTGAGATTGAAGGGATAGAATTGATTTCGATGAAGGGATTAGTAGTGAATATGATGTATATATATAATCATGCACTTGATACAAAATCCTGAATAAACGAGGATGTTTCTTGCAGGTTGAGATTGAACCTTCCATTGACGCGTTACGGTGTAAAACTGCGAGCGTTTGTTTTAAGTCGCTGAAATTAATTTAAGCTGTATTTAACGGAAGGAAGACGGAAGGTTGACACGGAGATGCTGATTCTTGAATGGTGATGGCGAGGTTGGAAACGTGAGGAAAGTGACGGGCTGGAACGGAATTTTTGTAAATTACGATGAAGGTCCCTGGATTATGTTCGAATTCGGAATCTTTTATTGCTTTATAAATCAATTTTGTTTTTAGTCCTTCACGGTATCATTCTCCTATTGAGTCCTCTTCTTAGTTACTAACTCACTCTCACCTATTTGGGGGTAAAAATGACATTTTAAAAGCACATGACTTTTCGAAAAGGGTATTTGTGTTAAAAACATGGATCTCTTTATATTTTTATTCATGCTTCTTCTTTGGATAATTTAAATTCGATAAAAAGAAAAACAAGGGGATCGAACCCAAACTTGAGACCTCTATTTTGTTTGATCCACGTCTTATCTTTCTTGATTGTTGTAATGTTTAGGTGAAGACAAGGGTCGAAGTTGAATTTTAAGAAACTTTCCAATCAAACAAATATGTTATTGAAAATTGGGAGTAGGTTTTGGCATGCAAATATTAATTAAATTAAGAAATCTTGGTTTTCAAATTTGGTTGTGAGGTTTCATTCGATTTGGTTAACGGTCAATGGAACTTGTATATTTATTTGATGGAATTTAGGTATGGAGATATAAGATCAAAGTTTTGATCAGAAATCTAAAATAATTGGTTGTCAATATACAACATTTTTTAAAGAGACCAAATTTGCAGACTTGAACATATGCCGGGACTAGTTTTGTATATTCAAAAATGGAAATGAGGTCATTTACTTCCACTACTTAAGTACGTACCTATATTTTTTTAAAAAAATAACTAAAATAGTCTAATCCCATAAGTGTTTTTCTCAAAACACCATAAATTAAGGTCTATTCCTCAAAATAGCCAACCAAAACGCAGATGGCCATTGCCATCTGCGTTTTCCTCCCATGGGTTGCTTTTTCCTCATGATGGGCTCCCAAGATCTGCGTTTTCCTCATGATGGGCTGCGAATTCGTCTACAAGGGCTGCATTTTCCTTGCTACCGGCCCATCAGCCCAAATATATAAATCAGAAAAAAAACGTAAATTTTATCGTTATCGTAGACGAGAAACGCTGTCAAATGGCTTCGCAAGATTTATCGACATGGAAACAGGTAAGTCTCAAATTTGATGTTCTCTTTATGTATTAATTTTCGTTTTAAGTGTTAGATTAGTGAACGTACAAATTTTTATGCCTAATGTACACGCCTTTAAAGCATAAAAAATTAGAGTTTATGAACATAGAAAGCATTTAGCAGACCATAGAGAGGAAAACGCAGATGAGGGCCATGAGCTGCGAAATGCTCTTGAGCATTTAGCAGATTTTGAGAGGAAAACGCAGATTTTGAGAGGAAAACGCAGATGAGGGCATGAGCTGCGAAATCCTCTTAAGCATTTAGCAGATTTTGAGAGGAAAACGCAGATGAGGGCCATGAGCTGCGAAATGCTCTTGAGCATTTAGCAGTTGATGCCCCAGCTGCTAAATGGTCGTTTTTTTTCCGAAAAACCTTAAAAATTAAAACAAATGACGAGTGTTCAAAGTTGGACAAAAATTTCAACTAAACCCATAAACCACCAAGAACCCTAACCCTAACTTTGAGGATTTCGCAGGTGATTAAGAGGAAAACGCAGACCATAGAGAGCATTTCGTAGACCATGCTGAGCATTTAGCAGATTCCGGATTTCTTAGAGCATTTCGCAGACCATGGGAGGAAAACGCAGACCATGCTGAGCATTTAGCAGATTTCGGATTTCTTGGGAGCATTTCGCAGACCATGGGAGGAAAATGCAGATGACAATGTCCATCTGCGTTTTGGTTGGCTATTTTGAGGAATAGGTCTTAAATTGTGATGTTTTGAGAAAAAAACTTATGAGATTAAACAATTTTAGTCATTTTCTCTTTTAAAAATGCACTTTAACCACTCTAATTTTAAAATTTAGAATTATTTTTTAAAAAAATGACACAAAGACTCTTAAGTTTATCGTTTTTTTCCATTTTAACCCAATATCCTTTTTTTTGGCATAAAAGTCTGAAACTTGCAAATATTTTTCATTCTTGGCATTTATTAGGTTTCTGTCTTTCTTAACCGTTAAACACTGTTGACATGGCTAGTGAAGTGCTATGATATGAAACTTTTGACTAATATGCGCATGCCTAAAATAAATCGACAAAATGCCCCCTCATAGGAATCAAACTTAAGACCTTCAGCATATGAGGCATTCCCTAAACCAGCAAACTAGCTAATGTTTTTTTATAGTTAAACATTTTCTAAGTCGATATAAAGTTAATCTATTCGTATATTTTTACCCTTTCTATTAATGATATTTGTGTTTAAATATGTTTTTGATGTATAAAAATGTATATTTTATCTGAAAATGCCTAGCATATTTTCGTTAATTTTTTTCCATATTAACATGGTTTATACGATAAAATGTGTATTTATATGAAATAAATACATATGTATACCTACTTATATGTATGAACGCATATAAACATGTGTTTTATTCGTATTTTATCATATAAAATGTAATTTTACAATTAAAATTTTACATAATATCATAAATACATTTAAACATGTTTAAGTATTTTTTTTATTAGTGTTTTACGTAGGGATGAGAATCAGTCTGGACCTGGACCCGGACCGAATCGGACCGAGAACTGAATAACCGATAACCGATTTCGTAGGAAGCGAATAACTGGGAACCAAAGCGAGTACCCTAGTTCTAGTATGGTTCGGTTATACAAGATAGTGTGATATTAAAGGATGTGTTCGGTTCCGGACCCGGCCCGAACCGGACAGAGAACCAAAGAACCAAGAACCAATTTAGTGGGAAATAGAGAACCGAGAACCTAACCAAATATACGGTTCCGGTTCCTATTCAATTCTGATTCCGGTTCTCCGGGTCAAAAGCCCATCCCTATGATACATTATTTTGATGATGGAAGATAAAGAGATATATTCACAAACGATTTTGCCTGTGGTCACATGTGATTGATATCCCCTTAATTTGAATCTGGGACACTTCATTCTTCGATACAAATCAAGTTATTGATTCATCATGTTCTTATTTTCTTTTTATTCATTGTTACATCAATTATTTACAAATTATAACAGTCATCATTAATATATCTATGTTTCTTATTATGCATAATATCAAGGCCCAAGCATATCACTAATTAGGGATGAGCAAAACCGAACCCGGAAACCGAGAAACCGAAAAAAACCGATAAAACCGAAACCGAAAAACCGAAACCAAAGCGGAAACCAGTGGTGAGGGTTTGATTTTTTTAAAACCGAAACTGTCGGTTCGGTTTCGGTTTCCCATGTCAAAACCGACCCATAGGAACCGAACCGACCCATTTATCAGAAACCGACCCAACCCATCATTTGAGGTTTCGAATTTAAAGTTTTCAAATCTTCATGAATTGTCGGCCTTGATTCTANNNNNNNNNNNNNNNNNNNNNNNNNNNNNNNNNNNNNNNNNNNNNNNNNNNNNNNNNNNNNNNNNNNNNNNNNNNNNNNNNNNNNNNNNNNNNNNNNNNNNNNNNNNNNNNNNNNNNNNNNNNNNNNNNNNNATGGAAATGGAGTGTGTCTGAAATTGAATTGTATAATATTTTGTTTGGTCTTAAACACTAGTGTGTGCTTTCGGTATAAACACGACATAACAAAATAGTAGATTTGTTTCTATGGAAACTTCCATTCCCTGTAACCAATTCATATCTTTTTAGAAACGAAAAAGGTTAAATCTAAAAAGCTCACCAATATACATATACAAAAATCGTCATTTCCAATATCCATATAAATAATTCAAGTCCCGATGATATAATTACAACATAAATCTCTATAACTAAACATACCATAAACATGGGTTCAAAACCGACCCACGGGTTTAAACTCAAAACCCACGAAACCATGGGTTCAAACCGAAACCGACCCAAAATCGAATGAAAAAACCAAAACCGAAACCGACTAGAAACCAAAACCGACGGTTTGTAAAAACCAGAAACCGAACTGATCGGTTTGGGTTCGGTTTTAGGTCTAANNNNNNNNNNNNNNNNNNNNNNNNNNNNNNNNNNNNNNNNNNNNNNNNNNNNNNNNNNNNNNNNNNNNNNNNNNNNNNNNNNNNNNNNTTTGTAAAAACCAGAAACCGAACTGATCGGTTTGGGTTCGGTTTTAGGTCTAAAACCGACCCAAACCGAACTATGCTCATCCCTAGGAATTATATCGCCGAAAAAAAGGAGTCGTCTGATAAAAATGGAGCAGGTAAGATCCATCCTTGAACAGTTACTTGGAACACATTGGGTGGATTAAATCCCATTGGAGAGAGAGAGGnAAACCGACCCATGCTCATCCCTATCACTAATTATCAACAATAACAATTAGAGTAAATTACAGTTTTGGTCCCTATCGTTAACACGTTTTTTCAGTTTTGGTCCAAGTTTTTCATTTTCATAGTTTTGGTCCCTATAGTTGCCAAAAGCTTGTATTATTGGTCATTGGTCCATGTAAACATACAAAATTACCTTTTCTAATTTCTTTTCCTTTTTTCTTTTAACTTATTATTTATTTTAGTAATTAAAAAAATATACCCATCCCCACCCCTACTACCTCGTTTCCTTCTCTCTCTCTCTCTCTGTGATCCCATAGACAACCAGGTGGATATTTACATAACCATGAGAGAAATCTCTTTCCTTTTCTAATGGTTGCTAGCCGAAAGTGAACCTTCATAAACCAAGTACAAGAGAGTTACAGCGGTATCGTTCATTGGCAGACTCTTGCGGGAATTTTTCGTTGTTTCGGAAATGGAAGAGGTTTCGGGGCAGCGATATCAGTTCCTTTTTATCATACTATGTTTTCTTTATCGCCTCTTTATCTCTATCCCTCACATTATACACCGTGGGTTTCACTAATTCCTCTGTCGAAAATTTCTTCAAATTATCATTCAATGGTTTCCCTTGTTCGTATAGAGCCACAATGTGCCCATGAAAAGGGCAAATTTTCAAATCCCTTCGCGGACAAAGTCTAAACTTTTTCAAAGGTGCATGATATTGTTGAATTTCTTTGTGCCCTTCTTCATAAACGGTTCCCTGAATGTTTAGTTCGACAATTTTATTAGCTGAAATAACTGCATCATGGTCAACCCTACCCTAATGAATTTCAACCTATAACCCTCTCTGGTTAGCTATAGACTCATGCTTCGAACCCCAGTTGTCCAAAAAAGAACCCCGTTTCATGACCGGAGCTTCACCTATGAGCTTTCTTTTTAACAGAGCCAAATCAATATCACAGACTTTGTTTGTTTTTACAGAAATGTTTCTCTTATTTACGAGATCCTCACTTTGCTTACTTTGGAGTAATTACAGGTTTTGAAATATGAAATTATTGTTGCTTTCTCAAACTCAAGGTGCGGCAGTGAAGCGGTTTTTGACTCACAACGTTGCAGGCTTTGGTTAATAAGGTACACAAAAGCTTTAAAATCCTTGATATTGACTTTTTGGATAAAGCACAAGGTGATTGAGGGTACCAACTGGTGCAAGATGGGTTTCTGAGCTTGAGATTGGTTTTGGAAAACCCAATTGGGTTACAGGTCCTATAGGTGATGGTGTGCCTTTCTTCATTAATATTGTCACTCTAATGGAGACGTAATGCGATAAAGGAATAGATGCATGGGTAAATTTGGATGAAACAGAAATGGAGATTCTACAGAAGAATCAAGACTTACTCGCATATGCATCGGTGAATCCAAGTCCTTTAGTACCTGATGAAGTTGGTGCATTTGTTCGAGCAGGGGGCGAAGATATTCAAGCATCAAAGACAAATGATATTGACATATTCTCCATTGTGTGTGTGTGCGTGTGAGAGAGAGAGAGAGAGACGGGGTTGTGGGGTGGTGGTTTATTTATTTAATTTTTTTATTACTAAAATAAATAGTGCAAGCTTTTAGTAACTACATGGAACAAAATTGTGAAATCAAATTGGACCAAAAATGAAAAAACTTACCAATCACAAGGACCAAAATTGTAATTTACTCTAACAATTAAAAAAGATCCTAGTAGAACCAGTCAGAAAGGTATTGTATACGGCAAACCGGAGAACAAAACTATAAACCGGGAAAAGGGCGGTCGGAAGACAGGCATCCTTACTTGGACCTGTAGTCGAAAGTATCTTGGTCGTTGTTGCAGATGGGACCCACGAATTTCATTTTATGAATAAAAAGTTGTAAAGCGATAAAACTACTCAACCCACTTCACCACCCTGTCACCACCCGTACCTGTCCCCACCTCCCACCCTCTCCCAGTTGCCGGTCACCCCTCTTCTCACTCTCCTCGGACCGGTCCGCTTCCGCACGTGCCAAGCCCCAAAACCTCTGTTTGTGCGTGTGTGGTGCAACAAATAAAAGCTAGAGAGAGAGAAGAGACATTTACTTCGTCTTCATCTTCTTCGTCTTATTGGACGCAACACACACGCATATGAGAATAAGGAAGCGTTTCTTGTCTCTTTCCGCAGCTATTACTGCACCGCCGCCGTCGGATCCTCACGGCGGTGACCACAAACACAAGCATCAACAGCCCCTACTGCTGGTGCAACAACAAGTGCACCCAAATGGGAATCTACGTGTTCTTTACGATAATACCCAGTCTCGGCCGTTCGATCCTCCACTAAATCATTGTAGCAGTCATCTTCCACCGACCTCCTCCGATCTTAGCACGCAGATCGGGCTCCAAACAGCCACTTGGGTTTCCACCGGTGCCGGCAGTGAGGCAAAGTACCTTGTTGTTTCTCCTCACAAGGTACACATTTTCACGAACAGAAACTGCACCATATACTTATAACTAGCGTGTATGTGTATATATACAGATCTTGCATCTATATATTTTGTTTTTTATTTTATCTTTTAAACTTTGAAGGAACTGGAATCTGAGAAAAAAGAAACAATCGAGACCGATGATACTGGCAGGTAAGATATAGCAAAGATCCCGTCCTCTTCCCCTCCCCATCCCCTTCTTTCTCCTCATTTCTCGGTTGATGTTCACCGGAAAATAAATTTTGAATCTTTTTGGGATAGACCCAGATTTTTTTTAACACCCACAACACAAACAAACATTAATTATTATTTATCTGATTAACCGAAAAAAGAAAGAAAAATAATTGCAGCAAGTGCAAACACACTGTAGAATAGTGATTTTATATCAGGTTAGAGAGCCAACCATATGTACAATCCCAATCCTATAAATTGCACCCATGTGGTGCATGTTAGCATGAAAACTGGTCCCCAGCAACTTCATCTTTGCGCGACTAATGCTCTTATCTTATCTATCGTTTATTTGTTCTTACTTTCGTTTTAGCAATCCACATAAACACACGCTTTTTCTTGTTTTGAATCGTACCGGATCTTTTTCTTCTTGTTCTAGCTTGCCTGTTTCCAGGAAAGAAAGCACCGTGTGTGCAGAAGCAAGCAGTCGTGTTAATACTGCTTCACTACCTCGACCACCAACAACTGCGCACCTTGGTATATATGTTAACCGTGTAAGCTTTTATAGTATTAATTTGTACGTAATGTATGATTCAGTTGTTTTTTTATCTACAGTGTTGAAAGGGTGGCTGGATGGAGATAGGCCGATTCCAATAAAAAAGAGAAGAGGAAGCTTTGGGAAAGGAACCAACCATCATCTTGATGAAGATAATGAGATTATTAACATAACAATGACGCCGAAGCCAAAGAACTATGATAGATGTGTCCGTGTGGTATCAAAGGAGGTGAATGCAATGTGTGCAAAGAACAAGAAGAATGGGAAGAGAGGGAATGTGATAATGGAAGGATCAAGATGTAGCCGTGTAAATGGAAGGGGATGGAGGTGCTGCCAACAAACTCTTGTTGGGTACTCATTGTGCGAGCATCATTTAGGCAAAGGAAGGCTTAGAAGCATGAATAGTGTTCGTGGGCGGGCACAAAAGGTTTCACTCAAAGAACAAGATAAAGAAGTTGAAACGATATCGATGGATCATCCTGATTTTGAAGATGATGATAAAGATTTGGAGGACGACTCAGTGTCATCTATGGAGGGTAAGAAGAAGATCAATAAAAAGAAGAAGCTTGGGGTGGTGAAGGCTAGATCGTTAAGCAGCTTGTTGAGCCAAATAGGCAGTTGATGCATGGAATCTTGAATTGATTGGTATCTTACGAAAATTAAAAAGAAAGAAACTGGTGATAGCGTGCATGTTCAGTCAGAATGCAAGAAACAATTAAACTATATATGCGCATTTTTGTTAGAAGGCTGGTCCTCAATTATTATATGTGTGATCATATAGTGCAACTCCTCTACTCATTGTTGGTTTATTTTCTTCTTTTTGGGTTATCATCACTCCTTAATTATGATCAAAAGCTTCATATGATGGGGTATATATATACGTTGTTGTTTTGAGCACAGCTATAAAAGTATCATATTTTTGTGTCATCTTCTTGGTAATCCGTATCCAAATGAGTATGGATTCGGTGCACACGAGCATCTAGATCAGCATTATAAGCATGTCATTAGTTCTCAGTTGAGGATAAGGGTTTTGTGAAGGTTCAACAGCCTTCAAGAATAGAAGATGCAAACCAAAACTGCACAAGGTATCGTGATTTATTAGCATCTTTTTTTATAGTTCCATAGAGTCCAGAAGTATCGCTAATATTGGCAAATTAACAGTCTGTTGGAAGGTTCTT

***CiGLN1;2***

TTTTTTTGGACGGAGGACGTGAAGCTTTTTGAAAACCAGAGGTAGCTTTCTGGTACTTGCGGCGGTGAGCTTTTGATGGATGTCAACAGGGCGGTGCTCTCCGTTCATTCCTCGGTTCCATGCGTAATCGAAGGTTTTCAAAAGTTGCGATTCTGTTTTCTCAGCTTCCTTTTTGTTGTTCATCTGCAAAATCGAAAGCGAACATGAAGCAATAAGAAAAAGAAAATTGGAATCAGATTATGTTTGTAACTTACCGGAGCCCATCGAGATTCCTTACCGATCTCCTCAACCTCGACCTTTCTAGTATCACCGATAAGATTATCGCCGAATACATATGGATCGGTGGATCGGGTATGGATCTTAGAAGCAAAGCAAGAACTCTTGAAGGACCCGTATCGGATCCCAAGAAGCTACCGAAATGGAACTACGACGACTCTAGCACAGGGCAAGCTCCCGGCGAGGATAGTGAAGTCATTTTATACCCGCAAGCAATATTCAGAGATCCATTCAGGAGGGGCAACCATATCTTGGTGATGTGCGACGCTTACACCCCCGCCGGCGAGCCAATCCCGACAAACAAGAGGGCCGCCGCCCCAAAATCTTCAGCAACCCAGAAGTCGAAAAGGAGGTTACCTGGTACGGAATTGAGCAAAAATACAATACAACCGCCAAAATCTTCAGCAACCCATCGGCATAGGTATTTGTTGTTTTGGCTTTGACCTTCAAAGCAAATTCGTGTGTTTATTCCTAGACCTAGAATTGGTTGTGCACGGTCGGTGGTGGTTGCTGGTAAGGCTAGATGGTTTTGTTGAGTAGTAATGAAATATATAGTTAAATTTAAAAAAAAAAGCCACGTGGACGTTATTTAAGTGCCACATGGACCATTCTGATGACTAAGATGACAAATGGCATGCCACGTAGGCATGTTAACCGGATGACCCGGTCATATCCTTTAAAATGATCTTTTT

***CiGLN1;2***

>scaffold6152

AGGTCGAAAGACAGTTGGGTGCAAATGGATCTTCAAGAAGAAGACCGACGTGGATGGAAAGGTACACACTTTCAAAGCTCGACTGGTCGGGAAAGGTTTCACTCAAACCCCTGGGGTTGACTGTGACGAGACCTTCTCACCAATGGCTAAGATAAAGTCTATTAGGATAATGTTCGCCATAGCTGCATTTCATGATTATGAAATTTNNNNNNNNNNNNNNNNNNNNNNNNNNNNNNNNNNNNNNNNNNNNNNNNNNNNNNNNNNNNNNNNNNNNNNNNNNNNNNNNNNNNNNNNNNNNNNNNNNNNNNNNNNNNNNNNNNNNNNNNNNNNNNNNNNNNNNNNNNNNNNNNNNNNNNNNNNNNNNNNNNNNNNNNNNNNNNNNNNNNGTAGATGCAAAGTTTCCCAATAGAGTGTGTAAGCTTGAGAAATCAATCTACGGATTGAAACAATCTTGTCGCAGTTGGAATCTCTGCTTTCATGAGAAAGTCAAAGAGTTTGGTTTCTCTAGGAGCGAAGATGAGTCCTGTGTGTATATCAAAGGCTAGTGGGAGTATAGTAACTTTTCTGGTGCTGTATGTTGATGACATATTGCTCATGGACATTCCAACCTTGCAAGAAGTTAAATCTTGGCTTGGGAAATGTTTTGCCATGAAAGCCCTTAGAGAAGCTGCCTATATTCTCGGGATAAGAATATTGAGAGATAGAAAGAAGAGACTGATTGGACTCAGTCAAAGTGTATACTTGGAAAAGGTACTAGAAAGGTTCAGTATGGAAAATTCCAAGAAATGAGAGCTGCCGATCCAAACTAATGCCAAATTGAGTAAGACTCAAAGTCCCAGTACAGAAGAAGAGATAGTTAAAATGAGTCGGGTACCTTAATCCTCAGCTGTAGGGTCGATCATGTACACTATGACATGTACTCGTCCTGACGTGTCCTTTTCCTTGAGCATGGTCAGTCGTTATCAGGGAAATCTGGGCGAGGCTCATTGGACAGCGGTCAAGAATATACTCAAGTATTTGCGAAGGACGAAATACATGGTCCTAGTCCTTGGTGGCAGTGATACGTTGAGAGTAAGTGGATATAGTGATGCCAGCTTCCAGACAGATAGAGACAACTTTCGGCCTCAGTCGGGCTGGGTGTTCCTGTTAAACGGTGGAGCAATTACGTGGAAAAGTTCTAAGCAAGAAACGATGGCTGATTCAACCTGTGAATCAGAGTACATAGCTGCAAATGAAGCATCAAAGGAAGCAGCTTGGCTGAAGAACTTCATCGGTGGCCTTGGAGTTGTCCCGACCATTCAAGAGCAAATGGAACTGTTATGCGATAATGAAGGAGCGGTTGCCTTCGACTAAGGAACCAAGAGATCATGGAAAGTCCATACATATTGATTGAAAGTACCATTACATCAGACATAGAGTAGAAGAATGTCACCTCATAGTGAAGGGGTATTATATGAAGATAATCCTGCAGATCCTCTGATAAAGGGACTGAGTAAGGTGAAGCACTATCAACATGCTAGGAGCATTGGCATGAGAGATGATATTTAGTAATTAGATAGTTATTTCGAAACTTGTAACAAAGTAAATGTAATTGACATTTGGTGATTAAATAATGTAGAATTATTTATGAGTATTGTTTACTGCTATTTGTCAATACTTCATTATTGTGTTTCAAAGTTTGCATGTTTTACTTCCTGAGTAATTTGATTATTCAAAAGTCCACAGTCGCTCATACTTTCGGAAGTAAGTAGTGAATTAAGACTGTCATGAATTGAATTGTAGACTGTCTAAGGAGTTAGACATAGCAATAAAGTTTCTGCAATGTTTATGAGTACTTAGAAATGGAGATTTGAGTATTGGACCAACCCGCGCTCAGAGAATTACTTCATGGATTCTATCACGAGTAATTCTGAGACGATAATATCTTATGATCTTGAAACAGAGACATATAAGTTGAAATTTACAAGTCGGTTGTGCATTAATAATGCGTAAACGCATCGGTATCTCGGTGTTATAAAACGTATTGTTGTGCGATTTGATGAGTAAAAGACACAAGCATATGAGTCAAAGTTTATTCGTTCCTTTTGTCCTAACGGGAAAATCTATATCTGTGGGCTCCTCGGTGATTTGGTGATTGACTTATAGTGCAAGGCCTAGCCAGGACTGAATTGATGTGTTCAATTAGAAGTCCTATTTCATCACAAATCATGAAACCGGGAAACATAGTATCAGACTGAGATTGATTCTAATCCATGTCTAAGCCATGACTATAATTTAGGAATATACTTGATTGACAACAGCACAGTCCTTTTGGGTTGCCCTCAAAACCTAGCAAAGGACAAGATAACTTAAAGAGATAGAGAAATATGATTTATTAATCTATTATGAGAATAATATATTAATTAGAAATCATAATATTTAATTAATAATTAATCAGAAACTAATTGGAATTAATTTTGGGGTTAATTAGATTAATTAAAAGTGCAGGGTCCAAATTGTAATCGTTCAATAGTTGAACAACATACAATTAGGGTATACCTAATTGGTTTGAAATGTAATATTTATTTCATATTGTTAATATACTTATTAGACAAACTTTAATTCAAAATATAAAATAAAAATACACTTGCTTCTACTTGAATAACGCTATCCTTCTAGTATTTCTTGAGATAAAGTTTTCAATTGCATCTTCATAGGGAATGTTATCCAATAGGTTGTTCTCAATAGCCATCAACACCAATCCATTAAGTCGTTCTTGTCATGGTAGATCGCAAGTAGGACTTCAATAACTTCAACTTGAAAAAACTTTTTCTGCAGATGCAACAGTAATATAAATGGTTAATAAAATTCTATATGCTATAGTTGTATTAGGAAAACAACAAACTCTTTTTAAAAATTTTAAAATATCAAAAGCTGTCATCTTTTCCTTTGGTAAAAGATTTAAGAGTAATTTTAACTCAATATATAAGTCATTTCCATCAATATCGGATTGTTCATCTTTCTTAAGTGAAGTTTCCAAAGACATACAACAAGCCTTTAAGCTCATATCATCTAGTGAATGCAATTTTTCAGAATTAAACAAGAAACCAAGCATATTATCATATTCTTCATATTCTTCATATTCTTCAAACCTTGTAGTGAGAGAAGTAATTGCTTGATCAACAATATATAAAAATTATTTAACCCCAAAAGATTCTTGTCTGAATTGTGAAACACTAGAAGATGCATCATTTGAAATCTCATCAAAATGTATTTTTCTTATAATTTGACGTCTTTGTGGAAATATTGGATCAATGTTCATTTCAATAGCTATTTATTTTGCAGTTTGTAAAGCATTTAGAAAACCATTTTCTCTATATTCCTTAAAAAAGAAATTAACCCATTTATTTTTTTCATATCAACATCAATAAGCAAATCTTTTGAGCAATTGTTTGCTCACTAAATTAACTGTATATAAAATTTCATACCAAATTAGTACCACAAAAAGCCAAGTTATATTTAGCATGAAATTTCATGATAGAGATAATTCTCAACAAAACTTTTCTCCATTAATCTTTTTCTTTATCTAATTGTTTTTGAGCTACATTATCAATAGTTTTATTATTTTGTAACCTCAGACGCAAATCATACAAAGTATTTATATTTCTAACATGTTCCATACTAGTTTCATGCTTTTTAATTATAGTGTTAAAATGTATCCAATCCCCAAAACCCTCATTTGCTAGTTGACTTTTTCCAATACCTTTTTTTTAATTACAACAAAAACAAAATATGTTATCGAGTTCTTTAGAATAAACAAGCCAATCTCTATTACATTTCTCCCAATTTGGTAAAAGTCTAGTATAATATATTGTAGAGAATCTTCTTAAATCTTTATCTTTAGGGCCTTTTTCAATGGACAAAACTCTTTTAGGACCTTTTATCTTTAAAATATTATCATGTTAGAGTCAATATAATTCCAATTTCTTGGATCAAATATATCAAATGTATGAAAATCATTTTCACTATCAACATCATCATCAAAATTATCATCATTAACGTTAACCTCATCAACATTATCATCATTCACATTAACATCATCAACATTAGCATCATTCACATTAACATCAACATTATCGACAGCAACATTTACACAAAGATTATCATTATTAACATTTTGGTTTCATTTTCTGAAAAGTTTGATGTTTGTTTGACTATAAACTTATCAATAACTCCGGTTTGAGATTTAGTTAAGTCTTCAATTCTTTGTTTTTTTCTTAGGTTTTTGATATCCTGATTAATGTTTTTTTTAATTCGAGGAGGCATATTAAATAAACAAATAATAGTAAACAGATTAAGAGAGAAAACATCTGAATGATTTGGAAGTTATTGATTTCCTTCAATCATTTTTCAATGAACCAAAATGAAAACTTGAAAGAAAAACAACAATACAAAATTAGAAACCGTGAAATAGAAAATCAATTTCAATTTCAGAAAATCCTAAAACATATACCTTAATTGCAATTTCAGAAAATCCTGTGAATCAATTGCAATTTCAAAAAAACAGAAACCATGATGGCGTGAATCAATTGCAATTTCTGAAAAGTGAGATTGGAAGTTGGAAAAATATTGATTCGATAATCTGAACATGGAAGTGAATCAGAAAGTCAGAAACCATATTACCGTAATTAGAGTCGTCGGTCTCCCTTCAATCCCTTATCTGCTTGCTACTCAGCTTCAATCTTGACAATCTTGAAACAAGAATTACAGATAATCGGGTCAAAAAAAATGAGAAAGAATGAAGAGTGAATTGAATTAGATTAGATTATTGGAATCTGATCAGAAAACGAGAAAGAAAGAAAGGCTGAAAAACCATTCGAACCTTGAACATATATTTGAGAATTAGATGTACCGTAGTAGAAATTCAATCTTCATTCCTGATTTTTCGAGATTGAAGTGCCGATTACCGAGACTGGCGGAGCTAATAATGATTATAGCTTCACGTTTCAATTAGCGATTGATTACCGCTCTTTACGATCTTTGCGATTTCACGTTTACGTTTTTACAACTATTAGGTTATGGAGTTGGGACTTAGGACGGCCTAATATAAGTATACATCATAGAACTAAAAAATTTTGGGCCCTTGATAGAATATGAGTCCTGTTCAAATGCACCACTTGAACCCCCTTTGGGCCGGGCATGTTTGTGACAGATAAACGAGAGAAACCACTGTCAACACAATATATCAACTTGATTACATCAGCCCTAATGTTATTTGCTTCAACTTTAAACCGTCATCACTATTTGCCCAATAATGCAAAATTTTGAGCTTTAAGGCTTAATACCGAGATCACCTTTCCTTAGATACAATAATCTTTTTCCCAAATAACCCATGCCATTTTTCGCCTCTTTATATCTTCACCCCAAAGAACCTCATTCTAAGCTTTTTTAGACACCACTGTTTTGAAAACCGAACCGAATTGGTCGGTTGAACCGGAACCGATCAGGTTATCATTTTTTAGAGCTCGAAAACTGTTTGTCGGTGTACCAATAAAAATTGGGTGGTTCGCATTAAAACATGTTTGACCACCCTCGTTCAACCACTTAAAAAACCAGTTTTGCAAAAGAATATAGATTTTTTTGGATTTTTCAAAAAAATAAATACATCTAGCCTAAAGGTCCTAAATGCTAAACCACTAAATTAGAACTCAGTTACACTCATCCCCCAAGCCCTCAACCTTAGTGTACAATTTTACACTTTACAATTCAAACCTGGAATACTCAATCGATCGGTCGATCAATGGTTGGGCGGTGTAATAATGGCATCATCCGGTTATTGGACCGGTTAAACCGGTAAGGCTATGCGGTATGTGGAAGAAATCATCCGCATCCTACGGAGGTGGCATAGGAACGCGCATGGAATGCCATGAACACCCCCTACCGCATCCCGTGCTCTACGTTCTCACACATTCCTGGGAAGACAACCCTGAAACGCCCCGCCCTCTCTTCCTATTAGTCATTTTTCTTCTCTCTCTCCCCCTATCGTGTTTTTGCGTCTTGCATACCCCCACAATCTAAGGTGACGAATTCGGTGACGGTCCCGTTTTGAAAACCTTGCTTGACACTTAAAACCATGCTCGGGGACTTATATGTGGTAGAATAATACATGTGGATACTTCCTAAAACCGATTTACAATACATTGCAAAATACAAATAAAAATACCTTGTTTTCTTACGGTTATTTAATTTTGCAACGGTTTTCACTAAACTATCTGGAATAACATTACATCATTTGAAAGATTATATTAAATACACTGATCATATCAAAAGATATCTAGTTTTTCACAATCAAATCAACAAGATATTTAAAATTTTACCAAATTTGATTCAGAGAAAATAATAGTCTGGGTCTCTAGGAATACACAACGTTACTTTATGAGTTTATTCCTTTTCCTTTAAATAATTGGTGCCAACAATGTTTGATTTATTCAAGGTTTATTTTGGAGATAATCAATTTAAAAAAAACCTATAAAAATTAATTTAATGTCTTCCTAACAAAGAGACCGCTGCACTATAAAAGGCAAACCCGTCTCCTTCGTTTCACGTCTCTTCCATTACTCTCTCCTCTATACACACAGTGGAAAGATTTTTAGAGAGAGAGTAATTAATCCTTTCAAACATGGCCCTTCTCAACGATTTGATCAACCTCAACCTCACAGATTCCACCACGAAAATCATCGCAGAGTATATCTGGTTAGTTTCTTTAATCTGAGTCGTGATCGATATCTTTTGACTTTTTTGTTTACGTTTCGAATCAAGAAACCTGAATCCCCTGTTTTACGCTATGAACGTCGTTTTTAATGTTTGGATTCTCAATAATAACAAGTCAACTCTATACACCTACTCTTCGCCATTAACACACACACACACACACACAAAAGAAACGATTATTTGATCTGGGGTTTTGGGGTTTTGATTTTTCATGCTTGATCTTATAAACTATCTTCAAAGTTCAAATTGAAAGTTTGACTTTCTGGTCAAACTTTTAAAAAGTAGACAAATTAGGGATTTTTCCAGTAAAGACTTACTATGGATTTCCACAACACTTGATATGATCGTCTTGGAGTTTTGTTTGATTTAGACTTATTTATCTCTCTACGATTCAAAACCAAATAAAGAAAAACATTATTATAGTTATAAATATCGTCAACATCGTTTTTTAGTTATACTTAATTTTGATTATTATCCAAACCACCCCCTAAATTTCATGAAACTATTTATAGGATCGGTGGATCCGGTATGGACCTTCGAAGCAAAGCACGAGTAAGTAATTTTTTATAATTCTTTAAACATTTTTGCTTATAAAACGTCGATTTCTTTGTGTTTTATCCATTGAAATTATCTTTTTCTATTTTTCGATTTTATTTCTTTTAGACTCTTCCCGAACCTGTGTATGATCCCAAGAAACTGCCAAAATGGAATTATGATGGGTCGAGCACTGGTCAAGCTCCGGGCGAGGACAGTGAGGTCATTTTATGGTAAGTTTTGTCCACTTGATTTATAATTTGAATCATCATAAATTTTTGAAAATTTTGTAAATTTAATCCATGAGAATGAGAACTTCTTGAATAAAACAGGCCTCAAGCTATTTTCAAGGATCCATTCAGGGGTGGAAACAACATTTTGGTATATTCTCAATCCTTTTCTTATTATTTGTTTGGTAGTGGGGGTGTTGTTATGACAACTGTGATCTTCCATGGTTTTACGTTACAAATTAATTGAAGGATTAATGGTATCCAAGAATCTATAAATTGATTTGTTTATTTTCAAAGAAAGATACTTTTACTTTGTAACTTTATAGAAAATCTTTAAACACATGATAAAACTGATACATACGTCATAAACTTTATATTTAATTGTTTTTGACAGATAATGGGGAGTGGGGTCATGTATTTATGATTGTTATCGAACACTTGGTAGAACAATAAATAGGGTATATATTTTTACCAAGTGTAAAAACATTTAAAATTACTTTTACGTTAAATGTCACAAATTGGCTATATTTTATTTTTCTCTTATTAATTTACAAAATGTTACATAAAAGAAGGCAACCATTGATTCGGTTTCTTTGTTCCTTTTTATATCGTATAGGTGATATGCGATGCGTACACCCCTGCCGGAGAGCCAATTCCGACCAACAAGAGGTTTGCGGCGGCCAAAATCTTCAGCGACCCCAAAGTCGAAAAGGAAATCCCATGGTATTTTCCCGATTTCACTTCACGAACTACATATTCGCCCCAATTTCGATGTTTATAATTTCAATTTCATGTAATAGGTATGGAATCGAGCAAGAATACACCTTGTTGCAGAAGGATGTCAACTGGCCGTTGGGCTGGCCCCAAGGTGGCTTCCCCGGACCTCAGGTAACCACGGCGGTCTCAGATTTACACCTATTTCTCATAATTTTGTCCAAATTTTAACGAAACTGTATTCGTTCGATGTAATTTAGGGACCGTACTACTGTGGTATTGGTGCCGATAAGGCTTTTGGACGCGACATCGTTGACGCTCACTACAAGGCGTGTCTTTACGCCGGAATCAACATCAGTGGCATCAACGGTGAGGTGATGCCCGGCCAGTGGGAGTTCCAGGTAGGACCATCTGTCGGAATCTCTGCCGGAGATGAGATCTGGGCGGCTCGTTATATCCTCGAGGTACTGCAAAAACCGATGTTATTATAAATAGAGCGATTTTTTCAATGTGCTCTTGAAGCGAATTGCTTTTTCTTACAGAGGATCACTGAGATTGCTGGGGTCGTTGTTTCGTTTGACCCCAAGCCCATTAAGGTAAAGTTTCGTTCGTTGTCAAAAAATTTCAATTTATTTGTTGTAAAAAGGAGTAATGTAATACTCACCGGAAACGCCGGATTTTGATCAGGGTGACTGGAATGGTGCCGGAGCTCACACCAACTACAGGTATTTTCCGGCGAACTTACGTAAACCACAATGGAAAATTACCTTTTTCATAATCTTAATCATACCCATTTGCAAATTTCAGCACCAAATCAATGAGAGAAGAGGGAGGATATGAAGTTATCAAGAAAGCCATTGAGAAATTAGGTCTTAGGCACAAGGAACACATTGCTGCCTATGGTGAAGGAAATGAACGTAGGCTCACCGGTCGCCATGAGACAGCCGATATTAACACATTTAAATGGGTACGTAACATAAATAAATAAATGTCATTTCATAGATATACTGATAGATGAGACTGTGTCTAGCTGATGTGAGGCAGTGACAGTGACAGTGACAGTGACAGTGACAGTGACAGTGACAGTGACAGNNNNNNNNNNNNNNNNNNNNNNNNNNNNNNNNNNNNNNNNNNNNNNNNNNNNNNNNNNNNNNNNNNNNNNNNNNNNNNNNNNNNNNNNNNNNNNNNNNNNNNNNNNNNNNNNNNNNNNNNNNNNNNNNNNNNNNNNNNNNNNNNNNNNNNNNNNNNNNNNNNNNNNNNNGTGACAGTGACAGTGACAGTGACAGTGACAGTGACAGTGACAGTGACAGTGACAGTGACAGTATTACTGACGTTTTAAAATCTTGTGTCAGGGGGTGGCGAACCGCGGTGCTTCGATCCGTGTTGGAAGGGACACCGAGAAAGAAGGAAAAGGTTATTTTGAGGACCGTAGGCCGGCTTCGAACATGGATCCATATGTTGTGACATCGATGATTGCCGAAACTACCCTTCTGTTGTGAACATGAATTTGTGAAGTCTTTATTTTCGGGAGATTTCAATTAGAGTCGGGAAACGATCCTTCTGTTTATTATTGTTAAAAAGTAGTATTTTCCTTTGTTAATTCTGTTTTAGTTGTTTCTGGCTTTAAATTTTTCGCAAAAAATGCGGTTTGCTTGGTGATATCTCCCATGGTGTTTGTTGGAAATAACGTTGATAAATTCTATTGTCTTAATAACAATGTGAAGGGTTGGTTTCGATCTTTTGTATGCTTTTGGTAGTTTTTGTTGGATTTTTGGGTTCATCGATATTGTTTTGTTTTCTTGTTCCCTTTGGCCCATTACCTTTGTCATTGTACTATGATTGTTACATTGGAATATGATTGGACAACAAACTTTCTTGGACAATTTTGATGATGAGGACAAGGAATGCATTCATTCATGTCTCTATTTGTCAATGAATAATCTACTTTTGTTGGATGAAAAAATTACAAATTTTATTTAGTCTTAAATACATTTTGTCGTCACTAAAGTAGGAAACAATATCTTATTATCTTCTGAACAAACAAGATAATAAGGGGGAGTTTGTTTTTTGTCTGTACATCTTCGTGACATCTTTGGTCTGCAGCCCGTAAACGTTAGCCCTTGTAGATTGTTTGCTTTTTAGAAGACCGCAGACCTAAAAACGTTTGCACGTCCTCTTCTTCTTGCAGACCTGGATTGAAGTTTGCAGTATTTTAAAAAACAAACAGTATTCTTATTACAGATTGCAGACATTTAGTCAATCTTTTTTTTATAGGTGCTTGCAGATATGGTCTATAGACGGTAGACATCTTAACTCTTAAAAAACAAACATAAACCAAGTCTTAAAGTAATGCCCTTTGTGGACAAGTATGAAAGCTAATGTGACAAATGACCGGTTGGTGGGTGGGATTACATGTAACTTTAGGAGTATGGTATGATGGGTTGCCGTTAACCTAAGTTGACGTGTTAGTGTTAGATATAATTATGAATTAAACTATATTGTGACAATAATGACATGGTTATTACAGCACATCATGACATGGATGAACCAAAAACTGAACAAAAACAAGCGGAAAGTGACAAAGATCGAAAATAGGCAGTCACATTTGTTATCGAAACAATAGAATTTATTGACGTTATTCCCAACAAACACAATGGAAGATATCGGTATATCACTAACCAACTGTATTTTGCTGCTAAAAACTAGAAAGCCAGAAACATCAAACATCTGAAACAAAATAAACGAAGGTGGATATTTATTTTTACGATAATAAACAGAATGACAATTTGCAAATACTAACAATAAAAATCATGATGCTTGATACTTCACAAATTGGCATGGGGTTCACATAGTGTTTCTTGCAACGAGAAAAAAATTATTTGTTGTTAATGGCATGATTACGAGTCATCCTTTTGTTTTGGCCTACATATACCTTCTCCACGCTATCAAGAAACAAAAAAATTGAAAAAGATCAATAATGGATTAATTTAAATGGTATTGAGCGACTAATTTCATCCACTCAATTATTCAATTTACGGTGGACATTTGTGTACCGGATTATAAAAAATCTATTTTATTTATTTTTAAAAAATTCTAGGCCTTATTTTTTATTAAATATTGAATTTTGTTGAAATTATGATTTCTTAGAACATCTTATTATATAAAAGACTAGCAAAAAGCTACAATTCTTTTACAAAATTATAGCTAATAACGGATTTTGTAACCAAGAAACTCAATTGGCCGTAAGTTTACGTTTTCTACTACCAAACCATATATACGAAAAATCAACAATATAATCAAATATTATGTTTTATATTTTCAACACAATCTCAACACCTCTAGCCACTAAATTTTGTCTCGTTAGTAATCGATTAAGCATGATTTTCCAAACAAGGATATTTACTTTCCTTTGAACACTATGATTCCATCTAGTCACCATATTAGCATCTTCCAAAATAAAATCATCAATAAACATTCTCGTGACATTGAACGTGAAACTATATGAATTATCAAGATCCTAAATCCATTTATCTTCAACATCTCTTGGCCTGAAAGAGACCAAGATATAGCAAAACCGTACGTGTTAGGTGGATTGTAGGTGTCTAAATCTCCAAGAAACAACCCCGGTTTACATATATTCGTGAATAGGTAAAAAGAACCGAAAACAGTCCCGCGTAGGATCGAGTAACAACCCAAAACACGATAATAACTATGTGGACAAACCTCGCAATAGGATAAAAAAAATTAGTATTTGAGATTTAATTTAACTTATTCTCAAAATCCCAAAAGAAATATATTCCATTGTAGTAATTATTTGCAAGATTTTGGATATTTCGGCTAAGATTGATTGTTTAAAAAGTAAAAATGGAAAAAAGTCAAATTTTGAACATTTTTTTTCACTCACTTAGTTCTTCTATTATAACTTTTTCTTTATAAAAAAAAGTTTTTAAAATTTAATTCTCCATAAAATGAGCATAATCATGAAATTTTGAAAGAAGGTATTTTAAAAAATAT

***CiGLN2***

>scaffold36799

CGATTAACGAATGCAGAAAAGGTTAGTGATTGTATAATTTGTATTGCATTACAGCATTAGCGATTCTCGATTTCTGATTGCTGGTGTGAATAACCTTTCTGATTTCTTATTTCTGATTATGGTGTGAATGAAGTTTTCTGATTTTTGAAAAGAAACTATCAAGCTATAAATATGTAAGGTTTTTATTTCTGTGTTTTTATTCCAATTATTCCGTAAGTTTCTGATTACTGTTTTTGCTGAAAATAAAATTTTnTGTGTTTTTATTCCAATTATTCCGTAAGTTTCTGATTACTGTTTTTGCTGAAAATAAAATTTTCATTTCTATTATTTAGATTTTGAATGTAAAGTTCTGCAATAAGGTAATTTCTTTTTTCAATTTCTGCTATAAGTTTTTCACTACATTGATGTTAATTATTCAGTAAGTAAGTTGTTTTACTGATGTCTGTTTTCTTTCAGATTTTTTGAATTTATGTTATGTTTCTTTTTTATAGCAGTTTGCATCTTTNNNNNNNNNNNNNNNNNNNNNNNNNNNNNNNNNNNNNNNNNNNNNNNNNNNNNNNNNNNNNNNNNNNNNNNNNNNNNNNNNNNNNNNNNNNNNNNNNNNNNNNNNNNNNNNNNNNNNNNNNNNNNNNNNNNNNNNNNNNNNNNNNNNNNNNNNNNNNNNNNNNNNNNNNNNNNNNNNNNNNNCTGTTTTCTTTCAGATTTTTTGAATTTATGTTATGTTTCTTTTTTATAGCAGTTTGCATCTTTATATTGATGGAAATTTTATTCATAGTTATAATTTTTTGTAGGAAATGTTAGAGATGAGCCATACAAATGTGAAATTCCTTGGATAGCTATTCAAATTATCGAGTTCGCAAATTCCTAATATGTTGGTAGCTTACAAGATTTTGTCATAAAATAATAGATGATTTTGCTTCAAAGAATGCACGAGACAACGTTTTAGATGCTTTTAGTAGGTTATTATTTGTAATTTTGTACTACTATGAATTTTGTTTGATTATAATATTTATTGCATTTTTACTATCTAAAATTTTTTATACAAAGGGCATTAATTTAATATATCGCCCCAGGCATTGAAAATCAATGGACCGGCCCTGCTTTCTTGATAGCTTCAAAGCACACATAAAACTTACTGAAGTAATTTTTTCCATCAGGCATTGNNNNNNNNNNNNNNNNNNNNNNNNNNNNNNNNNNNNNNNNNNNNNNNNNNNNNNNNNNNNNNNNNNNNNNNNNNNNNNNNNNNNNNNNNNNNNNNNNNTGGGCATTAAAAATCAATGAACCGACCCTGCGTATCTTCACCAAAGGTGACCACATCCCCACTAACTTAGAAATTAAAAGGGAAGCCATCGAGGGCGGTACGACTAACCCATTTTACTATTCAATGAAGTGCATAATTTTCCTATACAAATAGAAATGTCTTTACCATAAATTGTTAAATATATACTACATAAGTTAAATATTAATTTATAAGAAACTATTATTGTAAATATTTAATTAATAGCTTTAAAATGAGCACCATAGCATATATATACGATTTATAAATTTCATTTTTTTCAACATTTACATTGATTTTAACAAGTTTTCATAACATGTTAGAAATTAAGAAATAACTCATTACATATTATAATCATTATATTTTATTTAATATTAGCATACTAATAAAAAATATGTTATTTTACTATCAAACTATAAGTACATACGTTTCGAGTTGATGCTTTTGGTATATGATTGGAGATGAAAACTATTTGATGTTACAAGACACCAATTTATTTTTTGGAGTCATGATTGAAGTTACTCTAATGGTATCTCACAACAATTAATTACATATTAACCAAAACAAATCTACTTACACAAACAATTTGGAATTTTGGATAATAATACTACACACCACTTTTCATAACTAGTAATACAACTCTACAATAACAATTATGCACGTGATTTAAAAATACATCACACATATATCTAAAAAAATATCAAACACATATGCAATTGAGAACAAAAATAAATTAGAAATCGAATGAGTCCATCAAAGTTTTTAGGGATTTTGTTGATAAAAAGTTGGGGTGATAAAGAAAACCTTATCCTATTGGAGTAGAAGCAGGAGAAGCATGAGAATGATCGTTTCTCTTGATGACCTTAAAGAGACCATTGACGAATTTCCTCCACCCAAAACCCAACTCACCCCTATCTCCTAAGGAAAGGCCAATCATCCTTTTTTTATTATTATTTTATTATAAATAACCTTTTCCATAACCATTCTACAACTTCCACTTTTCTCTTTCCACTATCACTCGCCGGCTGCCTTTCCGGTATGTTCTTCATCCCCTACACTGCTTCTTTTTTGTGTTTTGATCGGAGTCGGAACTCGTTACTTTTGTAATGGGAATCGACCTCCATTGCCGTCGATACTTTGTGCTTTTTGTTTTAAGTCTTCGTTTTGTTCACTGTTGTTTTGAATGCATTTTATTTACATTCGGCGATGGTTTTTTTGGGTTACGGATATTTGACTGCCAAGTGATTATCAGACATTGGCCATTGCGTGTCTGCAAAAAGAAGACCTGAAAGTTGTAGATCTGTTTGCCATTAATCATTGTTTGACCTAAACAATTCGTACTTTCTCGTCCTTCTAGAATCTAGATCCCTCCCTTGTTCTCCATGTGCTCTGAAGCAAATGACCAAACTGTCCTTGTATACTTGTCAAAATACTTTATTTTTCTTCTTTAATCAATAGTTGGTCGTATATAATATAAATTTCTAAGTAGATTGCATCACTAAGCATCTTATTTATATATATCGCCTTTTGAATCTGATACCCTTTACGAATATTATACAAAGTAAAGGTCACAATTATAAATAAATGAAAGAAAAGTCACGATTAAACCAACCCCATTACAAGAATCTTGGATAAAGACCAAAAGTAGGTCATAAACATATGAAAGATATAGTTTAAGGAAGCAATAAGTTTTTGATCGTGGTGTGTGGCCCATGATAATAAGATATTCAAAAAATAGCATTCTACTTTCATTTATTTGTGTCATATGACATTCTATTTTCACTTCTTCCTATTACAACATTTTACTTTCCAATTTTTTCTTATCAAACATTTATACTTTTGTGTCAATGATCTCCAGCCTAGCTGGTGGTGAGGCGACGGGTTTTACCCATGAGGGGCAAGGTGGGGAGTTCGAGTCCCACCCTCAGGTGCCACTCGTGGGGTTTTTTCCCTGGGAATCCCTAAAAATGGGTGAGGGTTTGCCCTGTAAAGGAGCTCGAACCCGGGTCTCCGCTGTAAAAAGGCGGGGAGGCTTCCGTTGTATTCGCCGTTTCCAAAAAAAAAACATTTATACTTTTGTAGCTTTTTCAAGAATAATGCAATGAAAAGTATGAGAGATCCAAAAAACTTTTTCACTTTTTCACTTTTTACCTTTTTACTTTTTTACTTTTTCACCTTTTTCACAAGAATCTACATTATTGGCTTTTTGTGTTCTTATTGACTTTGTATGTTTTCATTAACTTTTTTTTTGTTGAATGAAATGTGTAGGTGAAAAATGGCTCAATGTTTAGCTCCATCAGTGCAATGGCAAATGAGGTTAACAAAGAATGCAATGGAAACAAGCTCCATGACATCAAAAATGTGGAACTCTTTGTCCTTGAAACAAAGCAAAAAAGGAGCACTCAAAACTGCTACAAAATTTAGAATTTGTGCTTCATCAAATGGAACCATTAACAGAATGGAGGACTTGCTAAACTTGGATGTAACTCCTTTCACTGATAAGATCATTGCTGAATACATTTGGTACATTCTTTTCCCTTTGTAAAACAATAAAGATTCGTTTTGTGGTTTTGAGATAAAGAAAAATGAGTTCTTACGTTCAATTATGGTATAAAGATTTTTACTTTTTGACATTTTGAAACAGGATTGGAGGTTCCGGGACAGATGTGCGCAGCAAATCAAGGGTATGCTCACTTTTAATAATCTAATTATTTTTTTATGATAAACCATTATCTTTTTTAAATGAAAAAAAAATCATATTTATTTTTTTATCATTTCTTTTTTAGACACTCTCGAAACCAGTTGAACATCCTTCTGAGCTTCCAAAGTGGAATTATGATGGATCAAGTACCGGACAAGCACCAGGAGAAGACAGTGAAGTTATCTTATAGTAAGTCCAATTTAATAAAAAAATAAAATAAAACTCAAATCTCATTTAACTTAATGAATTAATTATATAAAAACAAAATTCTATTTTCCAGCCCCCAGGCAATCTTCAAGGATCCTTTCCGTGGTGGCAACAACATCTTGGTGAGTTTAAGTTCGGTGATTCTAGTAAAATTATCAAGAAATAGCAACGTACTTTCATGCATACATGTATTTTTCTTTCTATTACTAACATTATACTTTCAAGTTCCAACTTTTTTCTTAATATGACATTACACTTTGAAAAATCCACTGTATAATTGGGTTGATTTTTCACAGTACGACGTTGTAATANNNNNNNNNNNNNNNNNNNNNNNNNNNNNNNNNNNNNNNNNNNNNNNNNNNNNNNNNNNNNNNNNNNNNNNNNNNNNNNNNNNNNNNNNNNNNNNNNNNNNNNNNNNNNNNNNNNNNNNNNNNNNNNNNNNNNNNNNNNNNNNNNNNNNNNNNNNNNNNNNNNNNNNNNNNNNNNNNNNNNNNNNNNNNACGAAAGTAGGATGCTATAATAAACAAATAAGTGAAAGTACATTGCTATTTCTTGCATTACTTACTATCTTAACTCTATTTTTACATTTTTAAAGGTAATCTGTGATGCATACACGCCACAAGGTGTGCCTATCCCTACAAACAAACGTGCAAAGGCTGCTGAAATCTTCAGTGACCCTAAAGTTGTGTCACAAGTGCCATGGTAGCTTTTTTGAATGAAGATTTTATAATTTTTTCATGCAAATAAACTGTTAAACTTGGTTTTTAATATTAACTAATACATTTTTGTTGAATGTTTTCTGATTAAAGGTTTGGAATTGAGCAAGAGTACACTTTGCTTCAGCCTGATGTGAAGTGGCCTTTGGGTTGGCCTGTTGGAGGGTACCCTGGTCCTCAGGTATTATCAATTTATCATTATCATGATATTTCTTCATATCAAACACTAAACATATTTTAGATGAAAGCCACAATTATTTAATTAATTAATTATTATTATTTTTTTAAAACCGTAGGGCCCATACTACTGTGGTGCTGGAGCAGACAAGTCATTCGGAAGGGACATATCGGATGCGCATTACAAGGCTTGCTTATATGCTGGAATCAACATCA

***CiGLU***

>scaffold2394

TTCAAAGTAAGAAAACATAGATCTTTTATAGGTTGCATGTGAACTTAGGTTATTTAGTTCATGGTTGCGAAAAACCTTTCAAAATCATCGTAAACTTGTGGGGGATTGCACGGTTAATAAATAATATGTGCAACATACCATATTTGAAAAAATCACCCTCTAAAATAGGTTTTAGTTGTAGTTTTAGTTTGTTGTTGGCGTTTGTAATTGTTTGAGGCATTATCGAGTAAAATTGAAATAAAATTATATTCTAATTTATATTTCACAATAACGTCTTGTAAATTTTTCCTTGAATGTGCACCGGTGTGCAGATAACCAAAATGTAATGATATATTTCATCATTAACCAAGATTCGGGATGAAAGCAATGTACGATAAAGCTAAGTAGGAATTGACATTTCCATGGGGACGTTCCAAAGTCACAAGTAAGCGACACTTCGTTGAAATGAAATAACAATCCATTATTTATGTATTAGTTAAACATAAATGAACAATATGTTTCAAAACTGATAATTAGAAATATCTATTAATATAGTTTACATTGAAATGAAAAACAATCCATTATTTATGTATTACTTAAAAGAAAAAGTATATATATGTTTCGGCCCCCGTGGTCATCATACTCGTGTTCCGCCCCTACAACCGACAATTCGTTGGTTGCTATGCTGAGCATGAGTTCAGAATGTTGAAAACGATCCTGGACACCTTAGTAGAATTGATGGCTGAACGGATGAAGAAAACATTGAACGAAATGGCGAAGAGCATAAGATTGCATGCGGGGCTGCCAAAAATGTTTTAGAGCGATACCATGAACATAACAACCTACCTAATCAATAGAGGCCTTCTGTCCCCATAGGGCTCAAGATTCCAGAAGAAGAGTGGCAGCGTAGTGAGATATCCCTCTGTCATTTGAGATTCTTTGGTTGTGTCTCTTATGTCAAGGTCAAAGACAGTGAGAAGGATAAACTAGAGGCGAAGGCGAATAAGTGTTCCTTTAGAGGGTAAGACTTGGATGACATGGGGTACTGCTTATGGGAAAGCAAAAATCATAGGGTTTTCAGGAGTTGGGATGTGACATTCAATGAAGGTGCATTGTCGAAAGACGTGTTTGCAGATTCTTCTGACCAAAGCTAGAAAAACCCAAGCCGAAGAAAGTTCAAATAGAGGTGGATATTCCTGGAAACGATCTTGAAGAGCCTTTCGGTACTGCAGGAAAATACGGAGTAACCAAAGGCAATGAGAGCGCAAAAACATTTGGTGACATAGGGTGAGTGAGAGTAGTTTAGCCTCAAACGACGATTCCTCAGATGAAGGGGAAGACGGTACTAGTGATGCAACATGAAGCATCCTTCGTCGGTCCACTGATGAAAGAAAACCTTCGGTTCAATACCCAACATTGACTTATTTGCTGCTAAATGGGAATAGAGAACCAAAAAATCATCCAGAGGCATTGAGGATGAAATAATCCATATAGTGGAAGAAGTCTATAGAGGATGAGTTGTGCTCACTTGACAATAACGAGCATTGATCTCTTGTCATGTTACCATCAAAGAATAATGCATTGACCAATAGGTGGATATTCAGGGTCAATGACGAAGCTGATGGAAGCAAGAGATACAATGATAGATTGGTGGTGAAGGGTTTCCAACAGAAGGAGGGGATTGTCCATCAAGTATGTCTCTATATGCATGGTCTCAAAAAACCCTATATATGAATGTATTAAAATGCATCATTTGCTACCTCTATGGTTCTCTCGCTTATGGTTTTTAATTTTGTTTGTTCTAAGTCAACCGTTCATCTCTTTTACCGATCAATTCGAGCAAGATGTCCTTAGAATCATAAGTCTACTTTTGATATATGTGTCATTCTTGGTGATAATCTTGTTTATTGGTCAATACAACACCTTATTTCTTGTTATAGTACTAGAGACTAATATAAAGGTGTTGCGAACACAGCCGTTAAGGTCATATGGATTTTTAATCTCCTTTTTAAACTTCATTATACTTTGTCTCATGCCACAATAGTTTTTGTGATAATTGCAATTTTGTTTTTAGTGTCTATTACAACACTGCATAACTAAACATATATAAATAGGATTACACTTTGCAAGAGAACATATTACTTTTGGTCATGTTATAGTTTTTATTATCTTATCAATTTGTTGACATTTTTACAACTAGCTTACTATTGCAATTACTTTAAAACTTCAGATTATCTAAGCATTCTACTGAGATTGTGGATATCTCTTAAATGAATAATATACTTTTTGTATTTATTTTGTATAGAGTTTAGAAACTCGTATGTACTTTTTTCGTACTTGTATATGGATATATAAAAACAACCCACTTAAAACTATTTCATGGGTCTTCCTTATTATCTATTAGTATCACTTATCAAAGCATAACCAATATTTTTATTAACTTTTTAATAATTGTAGTTGTTATTTGTAAACATTTTGTGGCGTTTTTAAAGAAAAATCATCCTAAGTCGGCTGGTAGGTGCGTTGCTCAAGGCTTTAGAAAGGAGGTATCGAGTTTAGACCCCCCCTATTTTCATATGTGGATTTAAATCCTAAATCCAGGCTAACCTAGCTCCTACACCTGATATGGACAATAGACCCGAACCCACCTATCCAACTTTCTCTTTTTTGCATATTTTACAACGTTTAATTTTTCTTACGAATCTAGTCCACAATCATTTAGTTTTTCTTACGAATCTAGTCCACAATCATTTAGTTCAGCCTGCCAAACTTTCACCTTTTACAATTTTGGCTTTGTGTTTACTATATTTACGAGTTTGGTCCACTTTTTATGTGTTTTTTTAACTTTGTCTTCTTATATTTTTAAAGCTTAGTTAAAAAAATACTTTTTATATTTCTTGTATATGTTTTAAAAATTTGTTTTCAAAAAATTACCTTTATACATGTTTGTTTGTAGGCATATTTTTAAAATTTTGACACTTTTACAAAATAGTTTTTTCTTCAAATTTATTTGCATGCATAAAACTGAAAAAACATGTAAAAGTAGTTAACTTTTATTTTTTGTCACAAAATTTAGTGCAACGTTTATCTAAATCCTACCGTAAGTTATCTTATATCTTAGACCATCTCCAATGGGAGTTCAATAGAGAGTATAATGAAATTAAATTGAGTTTAATATAGGAGAGAGATGATGACATCTATAGTAGAGTTCAATCCACATTGAACTCACATTGAACTCTACAAATAGAGTTTAATGGAATGATGTGTGTAAAATTTAATTTTTATTAAATCAAAATTCAATTCAATTTGTTTCTCTCTCCTACTTGACACTTCATTGAACTCTTAAAAAGTTAAATGCATTGGAGATGACCTTACATACTGTGCATAAACAGAACTTCCGCAACATTGTAGGGTATATAACCACTACTTTTATAAACATGTAACACCGAGTCCATGACTCACAATGGATAGAAAATGTCCCCAATTCTTATCGGTTTCAAACTTTTAGAACCGAAAATGGCCTCAACGAAATTGGTTTCGGATTTGGACTTGCGAGTTCTTGAATCAATTCCAAGTCTAAACTCAGGTATCGGGCGTTCCCTCTACTAGATTATGAAGTCGTGGCTCCATTAGTCTTTATCTTTAGGTCATAAATTAGAATCAAAATTGTTGACCTTTTGATTAACTTTGTGTGGGTAGAATCTCAATAGCATACTAGTAGCATACTAGCATCCAAAAACTATCGAGGAATGATTATGGCCTTTGGTGTCAACATATGGTCAGGTTACATGTGTCCACCCCTGCATCCATTTCACATCCATGAAGTAAACATCAACCAGGAACAAGCTTTGTAAAAGAGAGATAGAGCCGCGGGTGGCTCCAACTTCCCTTCCATTTTATCTTCACATCCCTCATTCTCTCCCATTCTCCACACGATCTTCCCCCCCTTTTTCCTCTCTCTTATTACACTTCATACTTACAACTACATGTTCTTGTTCTTCTAGCGCTTGCGTCTGTTATTAGCATGTCTCTTCAGTCTGTAGCTCATGTGAATGGCTGCTACCTAAAACCCACTTCTGTTTTCGCCACCAAAAGAGACTTTCTGTTTCTAGATTTTGTGGGTTTGGGCGCTAAACGAAGTAATCGGAGATTACTTGGAGCTGGTGCGAGTGTTCTAAAGAGTGTTACAGGGTTCCCGAAGCAAAGGAGCTGGTCGTCTTCTATCAAATCGGTTCTCGATTTTGATCGTGTTGATCATGCGGACACCCAACAACCTTCCGATTCCAAACGGAAGGTGCGATTTCGTTTTTCCAATTCGTTGATTGTTTCAATTGGTAATCCAATTTATTATTCCATTTAGTTTCAGTAAGTTCTATAGCAGATCGATAATTTTTCTAATTAACAAAGGGGGAGTGAACTCGACGACCAAGACACCATTATACGAACAGAAACTGAACCTTATACTTCTCTTCAAAATCGAGAGTTGACTAAATTGAGCATCTAGAATATGTTGACTGATTTGTTTTTTGTATAGGTTGCAAATTTGGAAGACATATTAGCAGAAAAAGGGGAGTGTGGAGTTGGGTTTATCGCTAATTTGGACAATAAGGGTTCTCATCAGATAGTTGAAGATGCACTCACGGCTCTTGGCTGCATGGAACATCGGGGTGGTTGTGGGGCAGATAATGATTCTGGTGATGGTTCAGGGTTGATGACTTCAATTCCATGGGAATTTTTCAATGATTGGGCAGAAAAACAAGGGATTTCCCCCTTTGATCAAATGCATACAGGGGTTGGAATGGTTTTCTTTCCAAATGATGAAGACCTTATGGAAAAAGCCAAATCTAGTAAGTTTTCACTTTTCAGTACCATTTTCTTGATTTTCACTAAACATAAATTTCTTTCTACAAGTTTTCTCACATATTCCTTATGATCTCTTTTAAAATGCTTGTCTAGTTATTGTAAATATCTTCAACCAAGAGGGCCTTGAGGTGCTTGCATGGAGGTCTGTTCCCATAAATGCCCCTATAGTTGGTTACTACGCAAGAGAAACCATGCCAAACATACAGCAAGTTTTCGTTAGAATCATCAAAGAAGATGACATTGATGATATTGAAAGAGAACTTTATATATGTCGCAAATTGATTGAAAGGGCAGTTAGTTCAGAAACATGGGGAAATGAACTTTACTTCTGTTCATTGTCAAATCGAACAATAGTTTACAAAGGAATGCTTCGGTCAGAGGTTCTTGGAAAGTTTTATTTTGATCTCCAAAATGAACTTTATAAATCCCCTTTTGCTATTTACCACAGGAGGTATAGTACAAATACGAGTCCTAGGTGGCCACTTGCTCAACCAATGAGGCTTTTGGGTCATAATGGAGAGATCAATACCATACAGGTTACATTGCCATGAAATTTCCTTTATTCCGAGATAGCAATTGCTCAAGAATAGCAACGTACTTTTGTTATATTTGTTTATTATAGCATCCTACTTTCATTTCTTCATATTACAACATTGTATTTTGAAAAATCTACCCACTTGATAGCAATGTGATTGCTATAAATAGGGTAAGTTTTTAAAGTATAATGTCCTAATATGGAAAAAGTTGGAAGTACNNNNNNNNNNNNNNNNNNNNNNNNNNNNNNNNNNNNNNNNNNNNNNNNNNNNAGTTGGAAGTACATTGCTGTAATTTGGTAGATTCTCAAAGTACAGTGCCTTAATAAGAAAAAAGAAAAAGAAAAAATGGAAGTACAATCATGTAATAGAAAGAACTGTAAGTAGGATGTTGTATTACACAAATAAATGAAAGTACATTGCTATTTCTTGCATTTAACCGTTCTTCTAATTCATGCAGGGAAACTTAAACTGGATGCAATCACGAGAGAACTCATTGAAGTCACCCGTTTGGCGTGGGCGTGAAAATGAAATTCGGCCTTTTGGAAATCCCCGGGCATCTGACTCAGCAAATCTTGATAGTGCAGCAGAAGTAAAGCATACTGACTGCGCCTTTTACTTCTTGTTATTTTCTTTTGTATAATCTTTGTAATTTTTCAACTGATTAAACATTTATAATTGTTACTAGTTGTTTATACGAAGTGGGCGTACCCCAGAGGAAGCCATGATGATTCTTGTCCCCGAGGCATACAAGAATCACCCAACTTTGTCAATCAAATATCCCGAGGTAAGTTTCTTTCTTGAACATGTACTTAAATAACAGAACAAAGATGATTTTTATATATTATAACTATAAATATAATGTCTTTTTTAGGTTCTTGATTTCTACAACTACTACAAGGGCCAAATGGAGGCATGGGATGGACCTGCATTACTCCTATTCAGGTAGAGTAGAGTTTAGAGTCTCTAATTACTGTTTGCTATTAACTTTTTAATTTTTGTCCACATTTTATGTCGGTAGAAGTAAATATTTCTTGTAATTATTTAGATAGGACATATAATTTATAAGTCAAGTTAAAAAATTGTTTTGGAAAACAGTGATGGGAAGACAGTTGGAGCTTGTCTTGATCGTAATGGACTTCGG**CCTGCTAGATATTGGCGAACA**GTAGACAATGTTGTCTATGTTGCATCTGAGGTGTGTCACAATATCCCTATTAAAATTTTTGTATTTTGATAGTTGGATACAACTTATACAAGAAATAGCGACGTACTTTGTTGATTTTTTTTATAGCATCCTACCTTCATTTCTTCCTATTATAACGTTATACTTGTAAAACTCTATCCAGTTATAGCAATGTCATCCAAAACAACTTTTCAAAGTATAATGTCCCGATAAGGAAATAATTGAAAGTACAATGCTATAATTGGGTAGGCTTATAAAGTTACAATGTCTTTATAAGAACAAAAATAAATGAAAGTAGGATGCTTTAATATACAAATAAATGGAAGTGCATTTTGTTATTTCTTGCAATTAGTCCTTTTTTAGGATTCTATCTTTTACATTTATCAGAGGTTATTTTCTGAATGTAGGTGGGAGTTCTACCAATTGATGACTCAAAAGTTACAATGAAAGGGCGTCTAGGTCCAGGGATGATGATAACCGTTGACCTAATTAATGGCCAGGTTAGTACAAAGATTGTACTTTTTGTTACACGGGAATAGACAAGCTTTCTAGGGCTTTTAAAATGAAGAGGGCATTCACGTCTTTTGCAAAGCGAGTGACCGTCTCACCTTTTTGGGTGGTAAAAAATGACAATCTTTTTAAAACAAATATATAAACTGAATGTATTCTGATAATGGAACAGGTTTATGAGAATACAGAGGTTAAGAAAAGAGTTGCTTTATCGAGTCCTTATGGAAAGTGGATAGCTGAAAACATGCGAAAGTTGGAGTCTGCAAGTTACCTTTCCGCCCCAACCATGGAGAATGAAACAACCTTACGTCGCCAACAGTAAGGAGGACATTCATGTCTTTTTGCATTTTGATAGCAAGTTTTGTCCCATTCTGATATCTTATCTGACATGAACAGGGCGTATGGTTACTCGAGTGAGGATGTCCAAATGGTTATTGAAACCATGGCTTCTGAAGGGAAAGAACCTACTTTTTGCATGGGAGATGATATTCCTTTGGCTGTATTATCTCAGAAGTCACATATGCTTTATGATTATTTCAAGCAACGTTTTGCTCAGGTTCATAATACCTACCATCCTTTTTCATCTATCTGCCATTTATTAAAATGAAATTAATTTCATTATTATTTCCAGGTTACGAATCCTGCTATTGATCCACTTAGAGAAGGGTTAGTGATGTCCCTTGAAGTCAATCTTGGTAAGCGTGGCAACATATTAGAGGTCGGACCCGAGAACGCCTCTCAGGTAAATTCTCCATTTTGGACGTGATTAATATCTCATTATGTATAATATTATTTTCTGTTCTACAAGTTATATGATTCATATGTTTGGGCAGGTGACTTTGTCTAGTCCTGTACTCAATGAAGGCGAGCTCGAATTACTATTCAAGGATCCGTATTTGAAAGCTCAAACTATACCAACATTTTTCGACATAAGGAAGGGACTCGATGGCTCCTTGGAGAAGACACTCAATAAAATTTGTGAAGCAGCTGATGAAGCCGTTAGAAACGGCTGCCAGCTACTCGTTCTCTCTGACAGATCCGATGAAATCGTAAGCACCGTATATATACAATTCACACAATGCGCTTGATACACAAAACTTGTGTATCATGCACAATGTGCCTGATACATGTATATCAGGCACTATTAGTATCAGGTGCATTTGTAAAATGCACAATGCGCCTGCTATATAGTTATATAGTCAGACACATTGTGCCTGATACATACAAGTATGTATCACACATAATGTTGTATGTGTGTGTGTGTGTGTTTGGCAGGAAGCCACGCGTCCTGCTATTCCAATTCTCCTTGCGGTAGGTGCGGTCCACCAACATTTAATACAGAATGGACTGCGGATGTCTGCTTCTATTGTTGCCGACACTGCTCAATGCTTCAGTACACATCACTTTGCTTGTTTGATAGGATACGGTGCAAGGTATTTTCATTTTTTGGTTTTGTAACATTTTGTTGTGGTCCAGATGGTTGATTAGTTGATGTACTTTTGAATCAATTTTTTGTAGTGCTGTGTGTCCTTACTTGGCATTAGAGACATGTAGACAATGGCGGTTGAGCAAGAAGACTTTAAACCTTATGCGAAATGGAAAGATGCAAATGGTTACAATCGAAAAGGCACAAAATAACTTCCGGAAGGCAGTCAATGCTGGCTTAATGAAAATTTTGTCCAAAATGGGAATTTCATTGCTGTCAAGTTATTGTGGTGCACAGATTTTTGAGATTTATGGATTGGGACAAGATGTGGTTGATCTTGCCTTTTCTGGTAGTGTCTCAAAGATTGGTGGATTAACTTTTGATGAGGTAATCACTAACCACATCTTTTTCTATGTTTTCACCATCAACTTAAACCTAAAAATTTTCATTAAAAGATCATCACCTTTTAAAATTTAATAATTTTTTATACTGAAATTTGCGCAGCTTGCAAGGGAGAGTTTGTCATTTTGGGTGAAGGCTTTCTCTGAGGACACAGCTAAAAGACTTGAAAATTTTGGATTTATACAAATGAGACCAGGAGGTACTTAAGAATGTTGATACATCTTCTTATAGCTCATTTTGAAAATCCATACATGAAACTAATTGTGTTTTGTTTTATAATTTAAAAGGTGAATATCATGGTAACAACCCTGAGATGTCAAAACTTCTTCATAAAGCTGTCCGTGAAAAACGTGAAAGTGCATATTCAGTTTACCAACAACATCTCGCTAACCGACCTGTCAATGTAAGTATGAATGATAATATATTTTGTTTTGGTTTTTTAGAAATAAGAATGTGAGAATCCTCCATAATTGAAGCATTATTCATGTAAAATTTTCATCACAGGTTCTTCGGGATCTCTTTGAATTTAAAAGTGACCGATCCCCAATTCCAGTCGGAAAGGTGGAATCCGCCGCATCAATCGTTGAACGTTTCTGTACAGGTGGCATGTCTCTCGGGGCCATTTCCCGCGAAACCCACGAAGCAATTGCAATTGCAATGAACAGAATCGGCGGAAAGTCCAATTCCGGAGAAGGTGGTGAGGTAATTCATCTCCTATATAATCAATCATACACCATTGTCTTGTTGACTTGTTGACCTAAATGAACATTTGATGCTGATATAATATATATTTTATTAAGGATCCGATTCGGTGGACCCCACTTTCGGATGTTGTTGATGGATACTCGCCCACACTGCCACATCTTAAAGGTCTTCAAAACGGGGATACTGCTACAAGTGCCATTAAGCAGGTGGCATCAGGGCGGTTTGGTGTGACCCCGACTTTTTTGGTGAATGCTGATCAGATTGAAATTAAAATTGCTCAAGGGGCAAAACCTGGAGAAGGTGGTCAATTACCGGGGAAGAAAGTCAGTGCGTATATTGCAAGATTGAGGAATTCGAAACCTGGGGTCCCGCTTATTTCTCCTCCACCACATCATGATATTTACTCTATTGAAGATCTTGCTCAGTTGATTTTTGACCTTCATCAGGTGAAAAAGATGTGAAATGTTGCACAAGTTTTACTGATATTAGTTTATAAAAAGATAAACGAGAATAACGAGTTTTGGGACTTTTTTTACCAGATTAACCCCAAGGCTAAGGTGTCTGTGAAGCTTGTGGCGGAAGCGGGAATTGGTACCGTGGCTTCTGGAGTTGCAAAGGGTAATGCTGACGTCATACAGGTATGTAAACAAATCAGAAACAATTGTTTTTGGAGAAATGAAAGTAGGGTGCTATAATAAACAAATTAGTAAAAGTATGTTGCTATTTATTCCATTGAAATACCTTACCTAAGGTAACACATGATCTAACGATTTAAATAAAACCATAAAAATCTATTAAGTAGTTGTTTCTGTATCAAGTAAAAAACACATGATGGATACTTCCCGTGACAAATGACAATCCTACCCTGCTTTGCAGATATCAGGACATGATGGTGGAACTGGAGCCAGCCCGATAAGTTCAATCAAGCATGCTGGTGGTCCTTGGGAACTTGGACTCACCGAAACACACCAGGTAATATAATATTTTAATTTTGTTTCAAATTATTTAGAGACTTGATTATTGTTTGTTATGTGAGTTAATAAGTTTGAGTTGATGTTTTGCAGACTCTGATTTCAAATGGTCTTAGAGAAAGGGTGATTCTAAGAGTTGATGGCGGGTTCAAAAGTGGAGTTGATGTTCTTATGGCTGCTGCAATGGGTGCTGATGAGTATGGATTCGGTTCGGTTGCAATGATTGCTACTGGATGTGTCATGGCTCGTATTTGCCACACTAATAACTGCCCTGTTGGCGTTGCCAGTCAGGTACTTAAATCTTTTAATTTACATTTAACAACCTATTTTACTCTTTTTACCAATATACGCAATCTACTTTCTTCTTTTCACCATAACAGAAATGCACTTATATATATATATATATTTACCCAAAGTAGCAATGTACTTGCATCTTCTTATCAATTATAGTAATTTGTAACCTGATGTGAAACTTATAAATTATAATTTGTCTGTTACACATGCTAGCAATGCATTTATATAATAGCAATGCACTTACATATTATTATCCACAATAGCAATGTACTTACATATATATATTTACCCATAATAGCAATGTACTTACATTTTTTTTATCGATAATAGTAATATGTAACCTGAAATTGATAATTTTTAATTTATATTTTACACAAAATAATAACATACTTAAATTTATTTACACATAATATCAATGTATGTACATTTGTTTACCGATAATAGCAATATGTAACAAATAGAGTACACTTCTCATATTGGTAGCAAAATTTAAGTACATTACCATTATTGGTAAAAAATATTTAAAAGTATCTTTTGCTATTATAGGTAAAAGACAGTAGGTTGCTATTATGGCTAAAACAATATGAGTAATCAAGTACGTTGCTATTATGGGTAAAGCAATATAAGTAGTTACATACGTCGGTATACACGAGAGTAAAGTACATTACCATTTTTAAAAAAATTTGGACTTGTAATATACTTTCTTTGATTGTATTTATCTTTTAGCGAGAAGAACTCCGAGCTCGTTTTCCTGGTGTACCCGGTGACCTTGTGAACTACTTTTTATATGTTGCTGAGGAGGTAATTTCTTTCTTTCAAGTCTCAAAAACCTACAAGCTATTATCTATTAATTTATAAAAATAATAAATTATATTGTATTCATGATTTACAGGTTAGAAGCACATTGGCTCAACTGGGGTACGAAAAGCTTGATGACATTATCGGACACACAGAGCTATTAAGACCTCGAGACATCTCATTAGTGAAAACTCAGCATCTTGATCTTAGCTACATGCTTTCTGTATGTATATGCTCTCAAAACTGCATTTTTCTTGTTTGTAAGAAAATAATCATAATATTTCATTTTTTATTTTCAGAATGTTGGGTTCCCGAAATGGACCAGCACCACAATCAGGAAGCAGGAGGTTCATAGTAACGGTCCTGTTCTAGACGACATTATGCTCTCTGACATCGAGGTAAATCTTAAAAAATTTACTGTTTCCTACTTTTTATGGAATTTACCTTTTTTTTTTCCCTACAGATATCAGATGCAATTGAAAATGAAAAGGTGGTAAATAAAACCTTCAAGATATACAATGTGGACCGTGCTGTTTGTGGTCGTGTAGCAGGTGCTGTTGCAAAGAAGTATGGAGATACCGGTTTTGCAGGACAGCTGAACATAACGTAAGATACTGATTCCAATATAAAAGTATTTTTTAATTCTATATAAAATGAAACTTGTTTAATAATTTTCATATTTAAAATTTTCTAGATTCGAAGGGAGTGCGGGACAGTCGTTTGCGTGTTTCCTTACACCGGGAATGAACATACGATTAATAGGAGAAGCCAACGACTACGTGGGAAAGGTATCATGTGTGGATGCATTGACTTTAGTCAAAGTTTGAAATTTTTTTCACATTGGAAGTTGTGTGTTGTTAGGGTATGGCGGGCGGTGAATTGGTTGTGAAGCCCGTTGAAAATACCGGATTTGTCCCCGAGGAAGCTGCGATTGTAGGAAACACGTGTTTATACGGAGCCACGGGCGGTCAACTTTTTGTCAGAGGCAAAACCGGGGAGCGGTTTGCGGTTAGAAACTCGCTTGCTCAGGCGGTGGTGGAAGGCACCGGAGATCATTGTTGTGAGTACATGACTGGCGGTTGTGTTGTTGTCATTGGAAAGTAAGACACAATTTCTTTTTTTTTTTTTTTTTTNNNNNNNNNNNNNNNNNNNNNNNNNNNNNNNNNNNNNNNNNNNNNGGATAAATGATGTTTACAAAGTATTTGAGAAAATGATGAAAAATAAATTGTCAACTGTTTGGTTGTCGTTTTGTAGGGTTGGTAGAAACGTGGCAGCTGGTATGACAGGAGGTTTGGCATATATTCTTGATGATGATGATACCCTCATTCCCAAGGTTTGTATCTAACTTTTTTTTTTTCCCTTTATAAATACAATCAAATCATGATTAGTTTCTAACCAATCAAATAATTTGTTAGATAAATAAAGAGATTGTGAAGATTCAGAGAGTGGTTGCACCCGTGGGGCAGATGCAGCTCAAGAGCCTCATCGAAGCGCATGTTGTAAGTACAAAAAAGCATCTCTTGACACCATTTAAATATCAAATAAAATTTAATTTGAGAAAGTTTCATTTTCCTGTCTTTATTTGATGTAGGAAAAAACCGGGAGCACCAAAGGCGCTACTATCTTGAAGGAATGGGACAAATATTTGCCTCTGTTTTGGCAGTTGGTTCCACCAAGTGAAGAAGATACACCCGAGGCTTGTGCTGAGTACGAGCAAACAGCCACTGGGCAGGTGACAAGTGTCCAGTCTGCATAAGAGGTCAACAATACACAAAAAGGATTCTTTCAAATCGATTTCACTTGAGTAGATAGCACGGGGTTCGGGTTCACTTGTGGAAAAAAGATTAAAATGGAAGAAATGGAGGTCTGGTTTGATTTGTTTGGTTGANNNNNNNNNNNNNNNNNNNNNNNNNNNNNNNNNNNNNNNNNNNNNNNNNNNNNNNNNNNNNNNNNNNNNNNNNNNNNNNNNTTTGTTTGGTTGAAAAAAAAAAAAACATGCCCTTTGAAGGTGAAAATGTATAGACAGTAGAGATGGTGCAAAAAGTGTAAATTCCTAGGGTACAATATTTAATAAATGTACTTTTGTGGAAGGATGGAACACCATAAGTATTTACTTGTATTTGCATATGTGGTGTTGCTACATATTCCATTTTATACTTTCGCATCAACCTTTCAAATTCGATGTTGAAATTCTTGTTCAGCGATACCATAAGTGTGGTTAGCATTAATTATAATACATACAATGAAGTGTTGTTGTCTGATATAGTCGTACAATCAACAGATATGGATTGAGAAGTTAAAAGAATGACTTGGTTCAAATCAATTTTGATCATGTTGGAAATTAACAGATTTCATTAGAGCGCTAAGGGTTTAAGCCCAATCTATAACATCTCGAAATTGGAGATATAATTAGGATTCATATTTAGTATTGTGGATGTTTTGGGAAGCCAAATAAAAGATGCAACACCAACACCAGACCTCGTAATCGCACTAGAATAGAGTAAGAGAGAAAAATGTCCGTAGTCATGCCTCCATTGGATGATCGTAGAGCAACTCCTTCACATGGATCACCTCTTTCATGTGAGATTGTGAGGCACAAATCTATAGTCGAAACAAATAAGTAAACTGGGTGTAAGAAGAAGAACAACACGGGACAAAGTCCTATCACATACCCTTAGTTTTGAGATGATCAATGCCACTCCTAACATATTTTTGATAGTCGCATTTGGAATCCTAGCCAACAATGTCCTACCGCATACCCACTACAATATCCTTATTTCTGCTACTTCCATCTTCCTCTAATAATCCTTCTTAATCGGCCAACACTCCGAATCCATACATCATAGCCGGTTTAACGGCCACTCTAAAAACTTACCCTTCGATTCTAACAGAACCTTCTTGTGTCACAAGACACAGGAGCCGTCCTCCACTTAAACCAATCCGTCTTTATGCGATAAGTAATATCTACCTTCATGCCTCCTATGGATCATAGAACCCAAATATTTATAAGTCTTATTTTATTCATTAGAAGTGTTTGGCAAAGTTGCAAAAGAATTGAGGATCATTTGACCATAGTACAAGGACTTATAGTGCAAAAAGCAAGTTTTCGGGCCTGATGAGACTTGGATCGGCCGGGCTTGAGGTCAACGGAGATGGAGCCCAACCGGACCATGACTCTATTGCCCCAAAACTTTTCCAAATCGAGTTCGACTTATGTCATGGCCCTTCAGTTGTTCTATCTCCTTTATATAAGTGTATAAAAAGGGTTTTACCACCCAAACCATAGTTTGACTTTTAAGAGTATTTTGGTCATTTTTAACCCTTTGACGACAATCATTAGGTGGTGAAAATGAGGTCTAGATAATGTCATTTGAACCTTAAAGAGTTGTGAATCATGGCATTTAGCTAAGGTGAATTTATTGGGAGTGCATGTCACTATACTTTGTTGTTTTTATTAGTTGATTTAATATAATATGTTTTTATGTGCAATGTGCTACCATACCTCTTACTTGTGCAATGCATGAAAGCTTATTTTCAAAAAAGTTATTTCATTTAAAAAGCTTGTTGATTTAATCGATTTTGGCATCGTACATGTGCATTCATAAGGATATGAGACTTTGTAGTCCTTCATTATATTTATGCGGGATTCAAGTGATTAGATTTCTATATAATCATCAATATGCACAAGTGTGGGTAGCTATCCCTTGTGTGGTGCATGTGTGATGATTCACATGTCACCTAGTTGTAGTTGGGCACGTAAAAGGGTATTATAGTTGGAGCCATTCATATATACCCTGGGGTGATTTTAGCATGGTGGTGTGATGCTTTCATGCTTGAGCCTATTTAACCATCTTTCTGAGTTTGGACCTTTGAGATGAACTTGGGTATTGGGATGGATGCATATTACTTTGTCATCGTCATTTATTGTTGGAATAATTACACTTGTTTTGTTAAAATTTATTGGAATTTGTTGATTCTATTTTGCATTTGGTTTTTGGTTTTAAAGTGATCTTTTTGGAGGTTTGGCTTGAGTTAGAATGATGGCCATCATTCTTACTCATAATAAGTCAACATCGAGTATTCTCACTAGGCATTCAATTGGCTAACTTATTTTGACATGTACCCATATTGAAGACTTAGAGACTTTGAATAGAATATAATTAAGATATTAAAAAGGTTACAGGTTGTGTGGTTTCATTTGTACTATGCGATGTTAATAATTGACGCTACCTTGCGCTTTTGAACAACTTATTGCATTTTTAACTATGGAACCGTATGACTTTCTATAACTTAAAACTCTGAATCAATATTGATTTCAAAATCAAATGATTTTAAAAGGTATTTCAAACTAGCCTAGTGAGGGGTGTTTCACAAGCTAGCTTCCCACTCTTTTACGTGCCTTTACCTAGGTCATTCATTTGATCACCGTGGTTAGATGTTTGGACCTTGTTACTCAATGAGTTTGAAAGGTTTTCTGCAACTATGAACTAAAAATCCTAAGTTCACATGCAACCTATTTAGATC

***CiGLT***

> scaffold4494

ATATATAATATCTTATCATCACATATATCAATTAATTAAAAATAATATATTTATATAAATAGGATGTATATTTATTAACCGTAAAATGATGGGTTTAAAATCATCCTAAAAATAACCATTTACTTGTTTGATAGTAAACCGTCTCTTTTGACTAATAAGACAACGTTGGGTTATTACTTCGTACAAAGATTAATAATTCTCGTTGCCAAACTTCACAAAAAAAATATATATATACATTACTACTAGTATTAGTATCATGTTAATATTACCAGCATGTGTTAGATATACAAATTATAAAAAGGAGTTTTTCAAGATTATTAATGTTTTGAAATTAATACAGTATCCGATCATTTTAATTATTAAATTTACAGTCAACAAATTAGATTGTTAAGTTAGACTTTATGAAAAAGTATGGTAACACATGAAAAACGCAAACAAAGTAAAGAATTGATTTGAATAAAAATAAATGTTTAGATTCTATCATTTCCAGGTACTGAAAGAAAGTTTACCCTAAAAAGTGGCAATTGGTAAAGCGGGGCCCACACCCAACAATGGCGACCTTGTTGAGAAGATGAGCTGGTACCCCCCTGCCTCCCTCTATAAAATCAACCTACCACCCACCTCGTCGAAGCCGAGAAAAACCCCCACCCACCAAACCCTCTCTCTCTTTCTCTTTCTCTTTCTCTCTCTAGTGCGTGTGTATATCCGCAGCCGTCGGCTTGCTACGTCGGTTGTTTAGTTTTGTTCGTTTTTTCTATCCAAGATTCCTCTTGAATCATATAAACCACCTGCATAGCGAATCCATATTCGAAACCCCAACTAAATCGACCTGTCAATCGGAGGAAAGACTGAAAGTGGGTGTTTGCTTTGAAGTTTGAACTGATTACTTCTTCAAACTAAGCTGTGTCTATCCAGATCTGATATTAAGCCTTTCATAAATCTGTTTATCGGAAAGATCTGGACCGGAGGAAGTTTTTGATATAATTATAGGCATCTTATAATGTCGGTGGCTTTGAATCATGCTGTCCAATCCCTTCCCGACTCGGACGTAACAAAACCTTTCGTTTCCCGTCCGTTAAACGGGTTAGCCCGGATTGGGGTTAGCCGGAGCCGGACATGGGCGAGCCGTGGCTCGGTTGTTAAACAGTCTAGTCTTTTAGGGAAGAAGTTATACGGGACTAGGCTGCTACGGGGGTCGGCTTCCGAGTCGCTCCATCAATGGAAATCGGATGGCCCGGGTCGTGACCCAAAGCTTAAGGTGGTGGTTCGGTCATCGTTGTCGCTAGTACCGGAGAAGCCCTTAGGGCTTTACGATCCATCTTTTGATAAGGATGCATGTGGGGTCGGGTTTGTTGCTGAATTATCCGGCAAAAGTAACCGCAATACGGTATGCATCGCTTTCCGTTACACTTACAACTTTGTTTTTTTGCGTCTGTTTTCAGTCAAAATTGACAACTGAATTACGAATTTACCTATGGGATCTGATTTTAGGTGACTGACGCGATTGAGATGTTGGTGCGTATGTCACATAGAGGTGCTTGTGGATGTGAAACAAACACCGGCGACGGCGCCGGAATTCTCGTCGGTCTTCCACATGACTTCTACAAGGAGGTATATACATACATCTACACGTAAAGTTCTTCCTATACAAACCTTAATCAGTACATAAACTAGCATATCGAAGCTAGTTACTAGTTTCATCTGGCTCGATGAATCGTTTAGGTTTATATCACTATCAACATCAAATCCGATTTGAAATAATAACAATTTGTGCGACCAGGTTGCAAAGGATGAGGGTTTCGAGCTACCACCACCTGGGAAATATGCAGTCGGCATGTTCTTCTTGCCTACTTCTGAAACTAGAAGGGAACAAAGCAAGATTGTCTTCACAAAGGTATCATAACTAATCTTTATTTCATGGTAAATGCGTAGTTAGGTTTTTCTACCGTATTTGATTTAAAATTTCACAAATTTATTTCCAGGTTGCTGAATCACTTGGGCACACTGTTCTTGGCTGGCGTACTGTCCCAACAGATAATTCTGGATTGGGGAAGTCGGCTATACAAACAGAACCAGTCATCGAACAAGTTTTTCTCACACCCACTTCTAGGTCAAAAGCTGATTTTGAACAGCAGGTATATATACGACTATACGAGCTTAAATTTCTTCATCTTTTTACCTTTTATCAACATAATAAATTACCAATTGCTATTTGCAGATGTACATACTAAGAAGGGTATCAATGGTAGCTATCCGAGCTGCATTGAACCTTCAACATGGTGGGATCAGGGACTTCTATATATGTTCTTTATCATCTAGGTTTGTTTTTCTTTTGAATTTCTTCAATTCTTGATGTGGGTTTTTAAAATTGAAATGATTTTTTTTTTGCAGAACTATTGTGTATAAAGGTCAGTTAAAACCAAACCAATTGAAGGACTATTATTATCAAGATCTTGGGAATGAAAGGTTTACGAGCTACATGGCCCTGGTAAATATTTATACCATATACTTATTATTTTTCCACTGACTTCATACTTAACAACAGCATTGGATCAACCGGTTAACGGGAGAGAGATGCTCGGTTAACTTATGAATTTGGATTGGATGCTCGATTCGTTAGTGGTGAGTTGGTAATAACCTCATTAAGAAGTTCATCATGTTTACAAAAACTTGGTGGCACCTTTTTGCTCGGTTCTTGATCATGAAACTTCATGAGGTTGTATAGGGCTTACATGGTGTTCTGATAGAAATCTTGATTACTTTTTTAACTATTATCATGGATGCAAATCATTGAAACTATTCTGTCTTTTTGTCTAGATTCATTCTAGGTTTTCAACAAACACTTTTCCTAGTTGGGATCGTGCTCAGCCTATGCGAGTTCTTGGCCACAATGGTGAAATTAACACACTTAGAGGGAATGTTAACTGGTAAGTCCTTTCTTGCACACCATGTGTTTGATCAAATGCCAATCTTTTATCTTTACTAGTATTCAGTTATTCATCCATTTTTGTCTACTTTTATGTTTCTTTATTCAGGATGAAGGCACGTGAAGGTCTTCTAAAATGCAAAGAGCTCGGTCTTTCCAAGAACGAAATGAAGAAACTTCTTCCCATTGTGGATGCCAGCTCATCAGATTCAGGTGCTTTTGATGGTGTTCTTGAGCTTTTAGTGCGAGCTGGTAGAAGTCTTCCGGAAGCTGTTATGATGATGATTCCAGAAGCATGGCAAAATGACAAGAATATGGATCCTCAAAGGAAGGCATTATATGAATACTGCTCAGCTTTAATGGAGCCATGGGATGGACCTGCTTTAGTTTCATGTATGTGTTTCTGTTTCATTTCACTCTCTAAACCCTAAACCCTAAACCCTAAACTTGTGTTTGGTTCTTGATACATATAATTTGTCAAATGCAGTTACTGATGGTCGTTATCTTGGAGCTACATTGGATAGAAATGGACTGCGTCCAGGTCGATTTTATGTTACTCATAGTGGAAGAGTTATAATGGCAAGTGAAGTTGGAGTTGTTGATATTCCACCTGAGGATGTTGCTAGAAAAGGCAGACTTAACCCTGGAATGATGCTTCTTGTGGACTTTGAGAAACATACCGTTGTAGATGATGAAGAATTGAAAAAACAATATTCGTTACAAAGACCATACGGAAAATGGCTTGAACAACAAAAGATAGAATTGAAGCACATTGTTGAATCTGTTAACAAATCCGCCCGCGAATGTCCCCCTATTGCTGGAGTGGCACAAGTACGAATCTTTTCATTGTTTTCCCAAAATCTTTTCTATAAAAATTGTGTTAACTTGTTAAAGTTTCAATTTTTAGGCATCTAATGAGGATGATAACATGGAGAACATGGGCATTCGTGGTCTTTTAGCCCCATTGAAGGCTTTCGGGTATGTGGTTTTAATACTATAACAGTTCTTGATTCTTGAATCTTTGTTAGGGGTGAAATGGGAATTTGATTAAAGTAACTATGCAGTTATACTATCGAATCTTTGGAGATGCTTTTACTCCCAATGGCTAAAGATGGTGTTGAGGCACTTGGTTCAATGGGAAATGATGCTCCATTAGCTGTGATGTCAGATAGAGAGAAGTTAACATTCGAGTATTTTAAGCAAATGTTTGCACAAGTAACAAATCCACCAATCGATCCTATCAGGGAGAAAATCGTAACTTCCATGGAATGTATGGTGGGACCAGAAGGTGATCTTACAGAGACAACAGAAGAACAATGCCACCGTCTTTCTCTAAAAGGACCACTTTTGTCTATTGAAGAAATGGAGTCCATTAAAAAGATGAACTTTAGAGGATGGAGAAGCAAAGTTCTTGACATAACATACCCTAAAGAACTTGGCAAAAAGGGTTTGGAAGAAACCCTAGATAGAATCTGTACAGAAGCTCACAATGCCATCAAAGAGGGTTATACTACATTGGTCCTTTCTGACAGAGGTATTAAGACACTACCCTTTTACCTATTTACCCTTCTTTTTCCTCCCAGCTTATAAAATTGATGAATATTATGCAAGGAAAAAACGGTCATTTAGTCTCATTAAGATAGTTTATTCAGCCTTTAAAAGAAACAAACTTTATGTAGACATCGCTTCACCCATATAGGACTTAATGGGAAAAAAAGAAACAACCCCTTATGTGTTTTTCTTATAATTTCTTGAAAATTGCAGCCTTTTCATCAAAACGAGTGGCAGTAAGCTCCCTATTGGCTGTTGGTGCGGTCCACCATCATTTAGTCAAGAAACTTGAACGTACAAGAGTTGCTTTAATGGTGGAATCAGCCGAGCCACGTGAAGTACACCATTTTTGTACTTTAGTTGGATTCGGTGTTGATGCCATCTGTCCCTATTTAGCAGTCGAAGCCATTTGGAGGATGCAAGTTGATGGTAAAATCCCACCAAAGTCAAACGGTGACTTCCATACAAAACAGGAATTAGTCAAGAAATACTACAAAGCATCTCACTATGGAATGATGAAGGTTCTTGCCAAAATGGGAATATCAACTTTGGCTTCATACAAAGGTGCACAAATCTTTGAAGCAGTTGGATTATCAACAGAAGTAATGGAAAGATGCTTTAAAGGAACTCCAAGTAGAGTTGAAGGAGCAACATTTGAAGCACTTGCTGGTGATGCTCTTGAGTTACATAATTTAGGGTTTCCAACACGCGAGTACCCTCCAAACAGTGCTGAAGCTGTTGCTTTACCTAACCCTGGTGATTATCATTGGAGAAAAGGTGGTGAAATCCATTTGAATGACCCTCTTGCTATTGCAAAATTACAAGAGGCTGCTAGAGGTAACAGTGTTGCTGCGTATAAGGAGTATTCGAAACGTATACATGAGTTGAATAAAAGCTGTAATCTTCGTGGACTTTTGAAGTTTAAAGAAGGGAAAGAAAAGGTTCCTTTGGAAGAAGTTGAACCTGCTAGTGAGATTGTGAAAAGATTCTGTACTGGAGCCATGAGTTATGGTTCTATTTCACTTGAAGCACATAGTACTCTTGCGATTGCCATGAACAAAATTGGAGGCAAATCTAACACTGGTATGTACACTTAAAGAAATACCCTTTTTGTTATTTCGTTGAAAAAGCTTATGAATCTGTCGTTGAAATTATTTTTGTAGAAAAATAATCGAAACCCTTTTGTTATATCTTTAAAAAGCTTATGAATTTGGAAGATCTTCAACACCCTTTTCTACGATGAAAGATCTTTAACACCCTTTTGTATTTTTATTAAAAAACTTATGAATTTGTTTTCGAGATCATTTGTGTTAAAGATCTACAACACCTTTTTGTACAATGAAAGATCTTCAACTCCCTTTTTTGCACAATGGAAGATCTTTAACACCCTTTTGTACAAATGAAGATCTTTAACACCAATTTGTTCTTTGAAGATCTACAACACCCATTTGTACAATGAAAGATCTTTAACACCCATTTGTTATATCTTTAAAAAGCTTATGGATTTGTTGTTGTAATCATTTGTGTTGAAAGATCTTCAACACCCTTTTGTTATATCTTTAAAAAGCTTATGAGTTTGATGTTGAATTTCAGCACCCATTTGTTAAATCTTCAAAAAGATTACTCATTTGTTGCCAAAATCATTTCTTTAGGTGAGGGAGGTGAGAATCCATCTCGTTTGGTACCGCTTGCAGATGGATCAATGAACCCAAAAAGAAGTGCTATCAAGCAAGTAGCTAGTGGACGATTCGGGGTTTCTAGTTATTACCTCACAAACGCAGATGAACTACAGATCAAAATGGCACAGGTACAGTAACTTTCATTCATCATTCTCTAAAACCTAAAGTCCATCTCACTCTCCCTGTTAGTTGGAACAATACGTTCCTCTTGTTCATGAATTTATGTTAATGGACAGATGATGCCGAATCTGCAAAGCACCTCATGGTGACTTTGCATCTTTAAGTGATTACCCCATTTGGCATTTGGTGAGGTTAAATAAACCACAATCGTTTATCGTATACCAAACAACTCTACCATTTACTTTCAAATCTCATCTTCATCACAGGGAGCAAAACCAGGTGAAGGCGGTGAACTTCCCGGCCACAAAGTCATCGGAGACATCGCAGTCACTAGAAACTCGACAGCTGGCGTCGGTCTAATCAGCCCGCCACCTCATCACGACATCTACTCAATCGAAGATCTCGCCCAACTAATCCACGATTTAAAAAACGCAAATCCCTCAGCTCGAGTCAGTGTAAAGTTAGTATCCGAAGCTGGTGTAGGAGTAATCGCAAGTGGAGTAGTCAAAGGCCATGCGGACCATGTCTTAATTTCCGGTCACGATGGCGGGACCGGTGCGTCCAGGTGGACCGGTATCAAAAGTGCTGGACTCCCATGGGAACTCGGTCTAGCCGAAACCCACCAAACCCTAGTTGCAAACGACCTTCGCGGTCGAACCGTTCTTCAAACAGACGGTCAGTTAAAAACCGGACGCGATGTCGCCATTGCTGCTCTTTTAGGAGCCGAAGAGTTCGGATTCAGTACAGCACCACTCATCACACTCGGCTGCATCATGATGCGAAAATGCCACAAAAACACATGTCCCGTAGGCATCGCCACACAGGATCCCGTCCTCCGTGAAAAATTCGCAGGAGAACCCGAACACGTCATCAATTTCTTCTTCATGTTAGCAGAAGAAATGCGCGAACTCATGTCCGAAATGGGCTTCCGTACTGTCAACGAAATGGTGGGCCGTGCAGACATGCTTGAAGTCGATAAAGATCTAACGAAAAACAACGAAAAACTAAAAAACATCGATTTATCGCTTCTACTTCGTCCCGCTGCTGACATCAGATCAGACGCTGCACAAACTTGCGTCCAAAAACAAGATCACGGTCTTGACATGGCGCTCGATCAACGGCTGATATCGCTCGCGAAACCCGCTTTAGAAAAAGGTCTCCCCGTTTACATCGAGTCGCCAATTTGTAACGTGAACCGTGCGGTCGGGACAATGCTTAGTCATGAAGTCACTAAACGGTACCATTTACCGGGACTACCAACCGATACGATCCATATTAAGCTTCATGGAAGTGCGGGGCAGAGTATCGGGGCCTTTCTTTGTCCCGGTATAATGCTCGAGCTTGAAGGTGACAGTAATGACTATGTTGGAAAAGGTTTGTCTGGCGGGAAAATCGTTGTCTACCCTCCAAAGGGAAGCGGGTTTGATCCGAAAGAGAATATTGTGATTGGAAATGTCGCGTTATACGGGGCAACAAGTGGCGAGGCGTATTTTAATGGAATGGCGGCTGAGAGATTCTGTGTGAGAAACTCAGGGGCAAAAACGGTAGTTGAAGGTGTTGGTGATCATGGATGTGAGTACATGACAGGCGGGACGGTTGTAATTTTGGGAAAAACCGGGAGGAATTTTGCTGCGGGTATGAGTGGTGGGATTGCGTATGTTCTTGATGTTGACTCGAAGTTTCGGTCAAGGTGTAATGCTGAGCTGGTTGATCTTGATAAGGTTGAAGATGAAGAGGATATTATGACTTTGAAAATGATGATTCAGCAACATCAAAGGCACACAAACAGCCAGCTGGCGAAGGAAGTTCTTGCGGACTTTGATCATCTTTTGCCTAAATTTGTGAAGGTGTTTCCACGGGATTATAAGCGGATTTTGGCCACCATGAAAGAAACGGAAAACGCTAAAAAAGCTGCTGAGCTGGCGGCTGAAGAAGCTGAGATTCGTGAGGAAGATGTGTTGAAGGAGAAAGACGCGTTTGAAGAACTTAAGAAGTTGGCAGCAAAGTCTTTGACTGAGACAGTTAATCAGTTGATCGAGACTGTTAATCAGGTGAATCTTTTTACTACCCTTTCGAATGGTATAGTATTTAATTTTTGGAAGGGTAGAAGAGTAATTTTTTGATTTGGGATTTCAGGTGAAAGAGGATGAAAAAGCCGAACAGGCGACCCGACCGTCTCGAGTTGCTGATGCGGTCAAACACCGGGGTTTTGTTGCGTACGAGCGTGAGGGTGTATCTTACCGGGACCCGACTGTTCGTATGAATGATTGGAATGAAGTTATGGAAGAGTCAAAACCGGGCCCGCTTTTGAAAACTCAATCTGCACGGTGCATGGATTGTGGTACGCCTTTTTGCCATCAGGTAACCAACAACTTTCTTTAACTAAGCCCGTTGTATAATTACTTCAATCTTTGGGTAGTCGTTTATAATTGTAAAATATAATTACAGGAGAACACGGGATGCCCTCTTGGGAACAAAATTCCCGAATTCAACGAGCTAGTTTACCAAAATAGATGGCGTGAGGCATTAGACCGGCTTCTCGAGACAAATAACTTCCCGGAGTTTACGGGTCGGGTATGCCCCGCTCCATGTGAAGGTTCATGTGTTCTCGGTATAATCGAAAACCCGGTCTCGATCAAAAGCATCGAGTGTTCTATCATAGACAAAGCCTTTGAAGAAGGGTGGATGGTTCCTAGACCCCCCCTCAAGAGAACAGGGTATGTAAAAAATTAAAAATATATTTCCATCATCTCTAAAATTGTCAAATCTAGTTGGTTTTCTCACTAGACTATTTATTTGCAGGAAAAAAGTCGCTATTGTTGGAAGCGGGCCCGCGGGTTTGGCTGCAGCTGATCAACTAAACAGAATCGGTCATACCGTGACCGTGTTCGAGCGGGCTGACCGGATCGGTGGACTGATGATGTACGGTGTTCCCAACATGAAAGCTGACAAGATCGATGTTGTTCAAAGACGGGTCGACCTCATGGCAAAAGAAGGTGTGAATTTCGTGGTCAACGCCAATGTGGGGACTGATGCATCGTACTCAATCGAGCGCCTTCGTGAAGACAACGACGCGGTTATTCTAGCAGTGGGGTCCACAAAGCCACGGGACCTTCCTGTACCGGGACGAGAGCTATCGGGAGTACATTTTGCCATGGAGTTTCTTCACGCGAATACTAAAAGCCTATTGGACAGCAATCTTGAAGACGGAAACTACATCTCTGCAAAGGGAAAGAAAGTGATTGTGATCGGTGGAGGTGACACCGGTACCGATTGTATCGGGACATCTATTAGGCACGGGTGCACCAGCATTGTGAACCTCGAGCTACTTCCCGAGCCACCTCGTACACGGGCCCCAGGCAACCCTTGGCCACAATGGCCTCGTGTATTCCGTGTAGATTACGGACACCAGGAAGCTGCTACAAAATTCGGAAAAGACCCGCGGTCTTACGAGGTGTTGACTAAACGGTTTATTAGTGACGAAAACGGAAAGGTCAATGGGTTGGAGTTGGTTCGTGTTCAATGGGGGAAAGATGAAAGCGGAAGGTTCCAGTTCAAGGAAGTCGAAGGTTCCGAAGAGATCCTTGAAGCTGATTTGGTCTTGCTAGCCATGGGCTTCCTTGGTCCTGAACCGGTTAGTAATTTTTACTAATTTTCAAACATATTGGTGTTTGCTAAATGAGTTTATATATTTATTTATTTAATTTTGTTTTGATGGTGATAGGCAATAGCGGACAAATTGGGGTTGGAGAAAGACGCGAGGTCGAACGTGAAGGCGGAATACGGGAGATTTGCGACGAATGTGGAAGGAGTGTTTGCGGCCGGGGATTGCAGGCGAGGACAGTCGTTGGTGGTGTGGGCGATATCGGAGGGGCGGCAGGCGGCGGCTCAGGTGGATAAGTTTGTGATGAAGGATGAGGAGGTTGAAGGCGGGAGGTTGCAGGAAGAGGAGGGGAAGAAAAGGGCGGAGCAACAGGCGGTTAGGACATAGACACACATACACATAGAGAGGGGAGTTGGGGTGGTAATTGATATTTAGTAGTGAGTGGTGAGGTGTGGAAGTTTAGTTTTGTATCCTCCTCCATCTCCTCTGTCTTGGTAGTTGCTGTTTATAGCCTTATTTTTTTCTCTCTGGTTTTTGGTGGTTTGCTTGTTTTGAACTTTGAAGAAATAAATAAATGTACAGATGAGTCTGTTTCGTTTTTTGCTGTGATAATTGGTTTCTTTATGCTCTCATTGGAATGAAATCTCCCAGTGATGTATTTATACTCCAACGGCAGTTTTATTCCCACGTTAGCCAAATTGATAAAGGCTTAAGATGATATTTCATAAAAAGTTCCCTAACATTTAAAAAACATATTCAAATTCAAAAACCGATACAATAAAAAGTTAAAAGCGTTTGTGCAAACACATCCAAATTCAAGTACTTATACGAATAAAAAGTCACAAACGTTAATGCAAAAGAGACATAACCACAATTCTCATTAACTCCCCAAAACAATTTTGGTTTTCGTTTCTTCAACATCCTTTCTGTCAAAATTTTCTTCTTCATAACGAGCTTTAAGTAGAAATAATATGCAAAGTTACATTTATGACTTCTTAAAAGATACATAAATAAAAATTCTACAATTGGTATAAAAAACTCTTCAAACGTTGTCTTAAAGTTTATAGTGGTGTAGCTTCTGTTTTTGTTTTCGTCAAAGTTCATATTTGTTATGTTTTGAGTTAGGTTTTCAACATTAACATTCGAATCGACTGTATACGAAATAGAAGAAAATGAATAATCAATACAAACTATTAGTTAATATATGTAAACAAGAAATTATACACCTTTTTCTCCAAACAACAAATCCTTATAACAAATTAAAGCTTCAACAATTTCAGACCATCATGGAACTTTGGTATTCATTAGAATTTTTCAACCGATACTAAAAGCTTATTCACATGCTACTGTTGAAATGAAGAAATGGAGAGACAAAGTATATCCATATATAATTTAGCTAATTCTGGATACCTATATTGTTGGGATTTCCAAAACTCAAGCATGTTAATGAAAGTCTTTTAGCTATTGACTCATGAAGGTATACTTGTAGTGAACTTTTTGTAGTCGAACTTGAATCATCACTATTGTTAGACTTGGCAAACGTGTCGTGTTGTGTTGGGTTCGTGTCAAGTCAAAACATTAAACGGGTTGAGAGAGATTGACCATAATATGACCCGTTTATAATTGTGTCGTGTCATGTCAACTGTTTATTTTCGTGTTGAGTTCGGTTAAAGGGTTTAGGGGTTAGCCTTGTAAATGAGTCATGCTGTATAATGTTGAAATAAAAAACAAGTTGAGAAAGTAGGAGTTGGTGAGGCAGAAATAAAGAAGTAAAAGATTAGAAAGAGTAGAAGTGAGAAAATAAAATGGAAAAATCCATTGCAAGTTTAGAGGGAAAGTGGAAAGAGTAAAAGCTGGAGAAGCAGATGAACAAGGCCAACATGATGAGAGAGTGGCTGAGTGGGATATGGAGCATATAAAATAATGGAAAAACTAAAAACCAAGCATTGGATTAGGTGGCAAAGTTGTTTTACATTACAAAGTGACTCAACACAAAGTTTTTCTTTTATATTTGATTTTAGTCTTGGGTCCATCGGATTTATCCAAGGAGTTCAAAAATTGCATATAAATAAACAAGTCATTTTCGAGTTGATAGTCTCAACTCTAACCTTACATGTTTAATAATCATGTCGTGTCGTGTCAACCTTAAAAACTATAAAGCAATTGTAATGAGAATATGATTTACCTTAAGAATGCTAAGAATGTTGTGACTAGGGCCTCATCCAACATTACCATGTCGAGAATTAATGTCGTTGCTATCTTGAAATGATGTCGATTCAGATGCTACTATTGACTTTTTTGAAGCTTGCATGTATTCATCTCAAGTGCAGAAAGGGAATCCTCAACCATCTTTAATTGTGATGAACCTTCATCATAGAGCTTATTATAGCTAAACTCAACAAAAGTGAACTTATATTGGGGATCGAACACCACGACTATAGCCCGATATTAGCTAAACTTATTCCATATTTGAGAAGCTATTTTTTCGTATACAGATCAGGACTTCCCATATATACATACACACTGACACATACATAATACACCATTGAAAAATCACACAAGGGAACGAGAGAAAATGAGAAATACTAAAATAAAGAGGAGGAAATGAGAGACCAAGTGCTTACCTTTTTCGTCGCGTCACGGGTGTTGCTACATAAAATATCACAATCATATAAGGGTTGACCTTCACCTATTGGCTATCGCTGGTGTAATGGTGTACTAGTGTTAGCGTGTAGCTGCATCGATTTAGATTTTAGGGCATAGGCACAAAGCTAAGCCGTAAACTCGTAACGAGCATTAGCAAAAAAACGTTTTCTATTTTTTTAAACCTAATATAATAATCGGGTTGTCTAGTTAACTTATGAACCTATCGACCAAACTGAACATGACACAAAACCATTTATTTGTGTGTCAAGTTGTTATCGTGTTTTGTGCCATGTTGATAATTTCTACCCCCCTAATTGTCATGTTCAAAATACTTTATCATTCACAAATTGTGTTGTCGCTGACTGCAAAAGCCCAAGTTCCAACTGAGCAGCATCCTGTTCTGCTGAGCAGGATATGTGCAGCGGAACAAGATACAGATCACGCATTCTTCCTCAACTGAGCAGTATCCACACGTTCTGGAGGCAATATGAATTCCTATGTTTCGAAGCGCTCATACGGTTGGGATGCGGCCTAGATTCGCTCGCCATATGAAGCACACAACCTTGAGTAGAACATCATGGGTCCATTCCATCATTCTTTCCATGGAGACAGTGAAGGGGCTATCAATTTTGAGCCTCAACACATCCACATAATATAACCCATCTCCAGAATATCACATGTACAATAATCAGGCCTCGAAGTCGGCTGGAAGGTGCCGAGTAAAGAGGAGAGTGTGTTTAAGTTGCTGACCTAAACAAAAGATGTGGGGGCCGATCTCTAATCCCAAGTCAAATGTCCGATTTGAACCCTGTCTTCTATCTTGCACCTTTTTCTCCAATCTAGTTGAAGAAGATCCGTGAAAACATCTTTGAGAGGGTCGTTTCCAATCCAATTATTTGTTGGATTAGGTGTCTAAGCCATGAATTTATATTTATATTTATCTTGTTAATTGAAGCAAACTCCATTTGGGTTGCCCTCATAGCTGGA

***CiAspAT1***

>TRINITY_DN14801_c1_g1_i5

TTTTTTTGGAAAATCTCAAGTATTATAAATTATGATAGACTTGATCAACACTGATCTGGATCCATCTATTAAAGTAATTCTACAATAACAATACAACAATCAATATCCACTACAACCAATCAGAACCCATAACTAGGTGAAGTCCTTTGGCCTTCTTTATTTTCTTTTAAAAAGCAAGAGATGATTATCATTTATTCTCAGCATATAATAATAAAATACCTGACTCAATTTAAAACCAAAAAACCTCATAATAAATCACCATAAAGGTCCTTTTAAGTTTTTGTAACTTCATGGATAGCTTTGGCCAAATACTCAACATTGCCAGTTGTTATCCCTGCCATACTAATACGCCCATTACGGGTCATATAAATATGAAATTCATTTGTCAAACGATCAACTTGTTCAGGAGTCATTCCACTATAGCAGAACATGCCGATTTGATTAGTTATGTGCTCCCATGAGAGAGGGGAACCCAATTTTTCAATGTTTTCCCTGAGAGCACTTCTCATTCCAATGATCCGGTCTGCCATACCCTTGACTTCCTTGAGCCACAATTGCTTCAAATCTGGATTACCAAGAACAGTTGAAACAATAAGTGCACCATGAACAGGTGGATTACTGTACATGGGTCTTGCAATTTGCTGCAGTTGACTTTTCACAGCCACAGCTTGCTTTTCATCTTCACATACCAAACTGAGGCATCCTACTCTTTGGCCATAAAGACCCATATTCTTAGCATATGATTGAGAACATCCAATCAAATGACCATCTTCAAGAAAAATCCTAATGGATTGAACATCTCTCTCTGGATCACCACTTGCAAATCCTTGATATGCCATGTCAAAGAAAGCAAAATGACCTTTTACCTTAAACTGGTAGGACATTTCTTTCCATTGTTCTACTGTAGGGTCAACTCCTGTAGGGTTATGAGCACAAGCATGCAGCAAAAAGAATGAACCTTTTGGTGCATTCTTAACATCATCCATGAGAGAGGCAAAGTCAAGACCTTTTGTTTCTGGATGATAATAATGGAATGTTCTTTGTGGTACATTAGCATCTCTCCATATGTTATGGTGGTTTGACCATGTGGGGACTGGAATATAGATATGAGAATCAGGGGAAAAGCGTTTCTGGAAGTCTGCAAAAAGTCTACATGCACCAGTACCAGAAAGAGCTTGTACTGCTGCAATTCTTTTATCTTTGATTAAATCAGAATTTTCCCCATAGGCTAACTTTAGAGTTTCATCAACCGTCTTGTTGCTTCCTCCCATTGGAAGATATTCCATGTTTATGCTTTCAGCAATCCGTCTCTCTGCTTCTCTGACACACTCGAGAACCACCGGCTTCCCGTTGTCATGACGATAAGCTCCATGGTAATTCATTCATTAGAAACCAAATACTATCTAAAGTAATCTTAACCAATGAAAATTTGGGGTTTGATCGGCAAAAATCTCAATCCTCGTGAAGTGAAAATGAAAAGAAAACAAAGACTTACTACGCCAACGTTAACTTTGTCAAGATTAGGATCTGCAACAAAAGCTTCGTGCATAGAAACCGAACCCGAGACGACCCCCTTGTTCCGTCATCGTCTTCGCCAAGAATCTTCTCCTCCACAGTTGATTTTAATTTCTGTCCAGAACTCCTCTCAGCAAGAACCCTAATTTCATATACAACAACATACAAAGCTTCTAATGCCCATGCT

***CiAspAT2;1***

>TRINITY_DN14194_c4_g1_i8

TCTATCTGGATCCTCGTGCCGACGTAACGTCGTCACGCCCTCATCCTTATCGTGGCACTATAGACCCCACCATCTCCATCCACGTCACCGCTTCCCTCGCTCGCCACGTCACCTTCATTCTATAAAGTCAGACCGATCTGAATATTTTTCAAGTCAATCGACTAAAAAGAGCGGTGCAGACGCAATAGACAAACGGAAAACGTAAACAAGTTAAAAATGAACTCCTCGTCGGCCGATCGGAGGTTGTATATGCTTTCACGTCACCTGGCCGGAACAATTAGTAATGAAGGGAGTGGTATCTCTTCTTCCCCAACTTCTGCCGGAAATTCTGTCTTCGCTAACGTAGTTCGTGCTCCCGAAGATCCGATCCTCGGGGTCACTGTTGCATATAACAAAGACCCAAGCCCTGTCAAGCTAAATTTGGGTGTTGGTGCCTATCGTACAGAGGAAGGAAAACCTCTTGTATTAAATGTTGTGAGAAAAGCAGAACACCTACTCGTTAATGACAGGTCTCGTATCAAGGAATATCTCCCCATTGTTGGACTGGCAGACTTTAACAAATTAAGTGCAAAACTTATACTCGGATCTGACAGTCCTGCTATTCAGGGAAACCGTGTGACAACAGTACAATGTTTGTCTGGAACTGGTTCATTGAGAGTTGGAGCTGAGTTTCTGGCTAAACACTACTACCAACGTACAATATATATACCCAATCCAACATGGGGAAACCATACAAAAATATTCACTCTTGGTGGTTTGACTGTGAAAACATATCGCTACTATGATCCAGTAACCCGTGGACTCAACATCCAAGGTTTACTAGAAGACCTAAATTCTGCTCCATCTGGCTCAATTGTCCTTCTACATGCATGTGCCCATAATCCCACCGGTGTTGACCCAACATTTGACCAATGGGGGCAAATAAGACAATTAATCAGATCCAAAAATCTCTTACCATTTTTCGATAGCGCATATCAGGGTTTTGCAAGTGGAAATCTTGATAAAGACGCTGAATCGGTTCGCATGTTTGTTAGTGATGGTGGCGAATGTTTCATAGCTCAAAGTTATGCAAAAAACATGGGCCTTTATGGTGAAAGAGTTGGTGCTTTAAGCATTGTTTGTAGGAATGCTGACGTGGCAAGTCGAGTAGAGAGCCAGCTAAAACTTGTGATAAGGCCAATGTACTCGAGTCCACCTATCCACGGTGCTTCAATTGTCGCCACCATTCTCAAAGACAGGTAAAATTTAAAAAGGGTATTTACGTCTTTTTGTCATGTTGACATGTGGTGGATTTTTGCAGAAACCTGTACAATGAATGGACACTTGAATTGAAGGCAATGGCTGACCGTATTATTACCATGCGCAAACAGCTCTTTGATGCTTTATCTGCTAGAGGGACACCTGGAGACTGGACCCACATTATTAAGCAAATCGGGATGTTTACTTTCACGGGGCTTAACACAAAACAAGTCGCGTTTATGACGAAAGAATTCCACATTTACATGACATCTGACGGGCGTATTAGCATGGCGGGTTTAAGCTCAAGGACAGTGCCTCACCTTGCAGACGCGATACATGCGGCGGTAACCACTGTTGGTTAAGTTAGTTGGTAATGATTATTATTATATTCTTTTTTATTTTTCTCCAGTTGTGGATTTTTATATGGGAAAATTGCTTTTGTTGTAATAAGGGACAATGCATGATAATCTGAATTGTGTGTTGTTTTAGAGAGAATAAAGGTTTGTTTTGTTAATCAATATTAGCTTATGGTCCATACAACTTTCACAAAATAAATGCAACACTTTTCTTAAACTTTGA

***CiAspAT2;2***

>TRINITY_DN18542_c0_g1_i2

CAAGTTTGACAGTAAACATGTCGGTTTAAAGGATTCGCTAGAAGGTTCTTGAAATCGATAATAAATTTGTTTAGACTTACGACAAAAAACAGGAACAAGTAGTTGGTTTATGCAATGGTGGTAACAGCAGCGTGCATAGCATCAGCAAGATGAGGGATTGTCCTTGTACTCAGACCAGCCATGCTGATTCTCCCATCGGAAGTCATGTAGATATGATATTCTTTCCTCATGAAAGCAACTTGTTCAGCATTCAGCCCTGTGAAAGTGAACATTCCAATTTGCTTTATAATGTGAGTCCAATCGCCAGGTGTTCCTTTTGCTTTTAGGGCCTCCAATAACTGTGTGCGCATGCTGATGATACGATCAGCCATTCCTTTGAGCTCGACCGTCCACTCATTGTACAAATTACTGTCATTGAGTATAGTGGAAACAATAGATGCACCGTGAAGAGGTGGACTGGAATACATTGGGCGAATAACCAGTTTAAGTTGACTCTCTACCCTACTTGCAACATCAGCTGTTTGGCATACAATGCTAAGGGCACCAACACGTTCACCGTAAAGTCCCATATTTTTGGCAAAACTTTGAGCGGTAAAGCATTCACCGCCATCTGCAACAAACATGCGAACAGGTTTTGCATCTGCATCCAGGCTGCCACTTGCAAAACCCTGATATGCACTGTCAAAAAAAGGTAATAAGGGTTTTGATCTGATTAACTTTCTGATTTCTTCCCACTGTTGAATGGTTGGGTCGACACCTGTCGGGTTGTGAGCACATGCATGAAGAAGTACTACTGCTCCTGATGGAGCATTGCCCAGATCTTCTAGAAGCCCTTGGAAATCAAGTCCACGTGTTGTAGGATCATAATAGCGGTATGTTTTGACTGACAATCCTGCAAATGTGAAGATTTTTGGATGGTTTCCCCATGTTGGGGTTGGAATGTATATGGTTTTTTCATGATAATGTCTTGCTAAAAACTCTCCACCAACCCTCAGTGAGCCAGTGCCAGATAAACATTGGACAGTGGCGACTCGCTTCTCTTGAATTGCAGGGCTATCAGCACCAAAAATGAGCTTGGCACTACCTTTATTGAAATCTGCAAGTCCAACAATGGGAAGATACTCCTTAATCCGAGACTGGTCATTAACAAGTTTTTGCTCTGCTTGCTTTACAACATTGAGAACCAGAGGTTTTCCTTCCTCAGTTCGATAAGCACCTACACCCAAGTTCAACTTTGAAGGGTCCGAATCTTTGTTGTAAGCAACAGTAACCCCTAAGATAGGATCTTCAGGAGCTTGGGCGATTTGGGAGAAGACGGAATGATCAGACATTTCTCTTTCAAAAATGGTTTTTGGTGTTTGCTGTAATTGGTTAAAAAATCTAGTGTGATTTGAGAGAGAAGTAGGGTTTTGTACGAAAGTGGATCTGGCGTTTGGAATATTGTGATTGTATTAATGACTGTGGGCGCAACTGGTTTTCAGTCGCTGTGTTTTGTGGACGAACCGTTTGCCGCACGCACTTTAGCGTCGGGTAAAAATTTGGCCGACAGTGATTCTGATTACATTCTTGCAACACCACTTACTAATATATTTTAATCGTATACTTTAATTATGATAAGAAAAG

***CiAspAT3***

>comp3246_c0_seq1

AGCTGAGAGATGGTTTCTACATTCCTTTCACTCCCCTCAGTTTCTCCATCTGCTGCCCTATCTGTGCAAGACAAGTGCAAGGACACAATGAAGTCTGTAACCTCTTTTCAAAGTTCTTTCTTTGGGAAAGACAAACAACCTCTCTCCATGAAAACAAAGTCTCATGGTCGTATAACAATGGCTGTTGCTGTAAAAGCTTCACGTTTTGATAACATAACAATGGCTCCCCCTGACCCCATTCTTGGAGTTTCTGAAGCATTCAAAGCTGACACAAGTGAATTGAAACTTAACCTTGGTGTTGGTGCATACCGTACAGAAGAACTTCAGCCTTATGTCCTCAAAGTTGTCAGAAAGGCTGAAAACCTTATGCTGGAAAGGGGAGAAAACAAGGAGTATCTTCCTATTGAGGGTTTGGCTGCATTCAACAAAGCCACAGCTGAGTTGCTGTTTGGAGCAGATAATCCAGTTATTCATGAACAAAGGGTTGCTACAATTCAAGGTCTCTCAGGCACTGGTTCCCTAAGAATTGCTGCAGCATTAATTGAACGATACTTTCCCGGATCAAAAATTCTAATATCATCACCAACTTGGGGTAACCACAAGAATATTTTCAATGATGCGAGAGTTCCTTGGTCCGAGTACAGATACTATGATCCAAAAACAGTCGGCCTAGATTTTGATGGAATGATCGAGGACATAAAGGCAGCTCCGGAAGGTTCCTTTGTGCTTCTTCATGGATGTGCTCACAATCCAACAGGCATTGATCCAACTCCTCAACAATGGGAGAAAATTGCTGATGTCATTCAAGAAAAAAACCATTTTCCATTTTTCGATGTTGCTTATCAGGGATTTGCTAGTGGAAGTTTAGATGAAGACGCATCTTCTGTGAGATTATTTGCAGCACGTGGAATGGAGCTTTTAGTTGCACAATCATACAGTAAAAATCTTGGACTTTATGCTGAAAGAGTCGGGGCTATAAACGTCCTTTGTTCATCACCTGATGCAGCTATAAGGGTGAAGAGCCAAATGAAAAGAATTGCAAGACCAATGTACTCAAATCCACCTGTCCACGGGGCCCGCATTGTTGCCAATGTTGTTGGAACCCCTGATTTTTTCAACGAATGGAAAGATGAAATGGAAATGATGGCTGGAAGGATTAAAAGTGTTAGACAAAAATTATACAATAATCTCTCTTCAAAGGATAAAACCGGAAAAGATTGGTCTTTTGTTCTTAAGCAAATTGGGATGTTTTCTTTTACCGGTCTCAATAAAGCCCAGAGTGATAATATGACGGATAAATGGCATATTTATATGACGAAAGATGGGAGAATATCATTGGCTGGTTTGTCTGCTGCGAAATGTGAGTATCTTGCGGATGCCATTATTGATTCGTATCATAATGTGAGTTAGAAAAAGGGGGAGTAAGTTTGAATAATTGGAGAAGGTGAGTGTTGTTTTTGTAAGTTTTGAACAAATGTGGGAATAATGTTTGGACTTTATATAGGGGGAAAACCCTTTTAACTTTTGTAGTTGAAGTTGTATACAAGTTATGTTAATTACTTCCCTTGGATTGGGGAAGGCATGTGTTGGAACTTATGGACTGTTAA
