## Supplementary material for "Asparagine accumulation in chicory storage roots is controlled by translocation and feedback regulation of asparagine biosynthesis in leaves": Sup S1-S12

**Fig. S1** Diversity of Asn accumulation in the chicory germplasm. Free Asn content (mg/100 g dry matter) in storage roots of selected chicory genotypes 180 days after sowing. Data are mean ± s.d. of 5 biological replicates containing 4 or 8 storage roots (n = 20 or n = 40). Statistically significant differences are represented by different letters (one-way ANOVA, *P* < 0.05 followed by Scott-Knott test, *P* < 0.05). Genotypes with letters a, b, c were classified as Low Asn content (LAsn), Medium Asn content (MAsn) and High Asn content (HAsn), respectively.

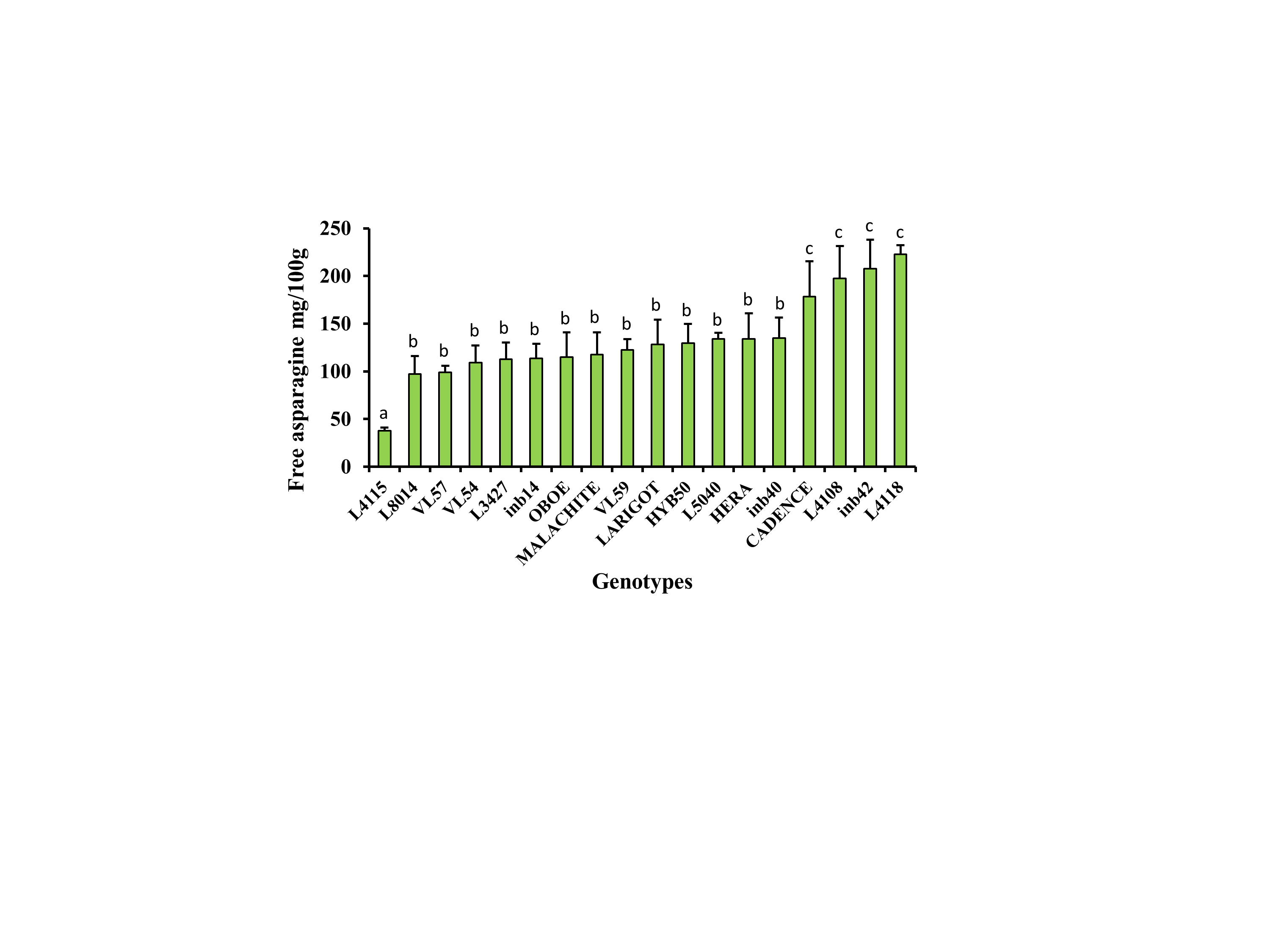

**Fig. S2** Correlation between weight (g) and Asn content (mg/100g) (a); diameter (cm) and Asn content (mg/100g) (b) of storage roots from the 18 genotypes analyzed for Asn content. Each dot represents 1 biological replicate containing up to 8 storage roots. Pearson correlation coefficient (r) was used to measure the linear correlation between the variables X and Y.

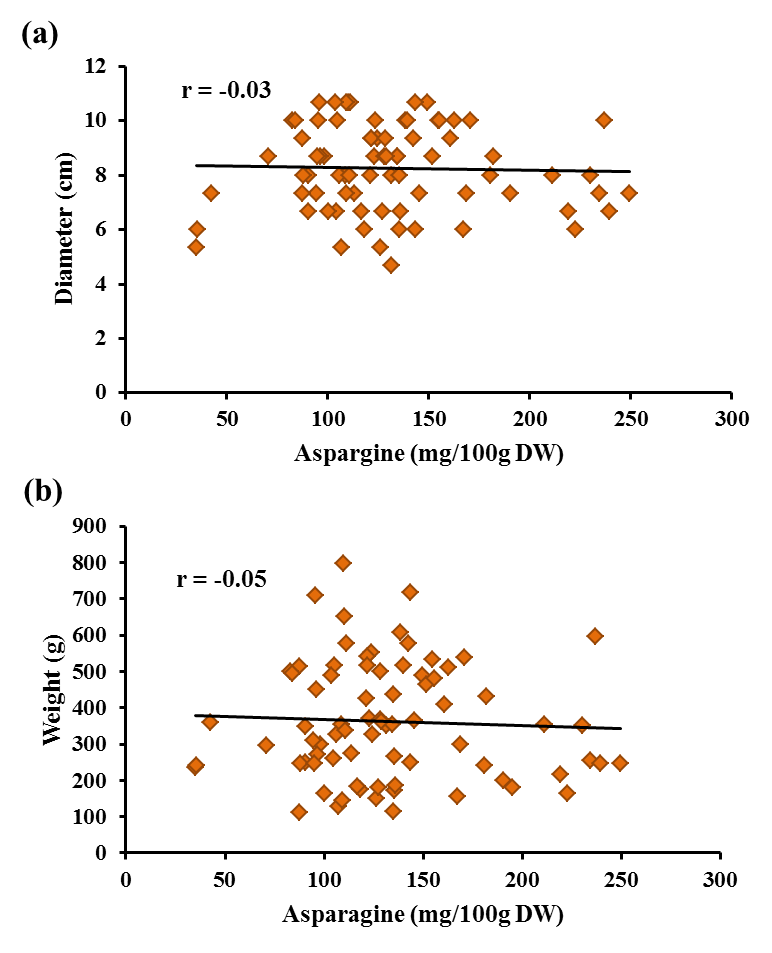

**Fig. S3** Phenotypic variation of diameter (a) and weight (b) in chicory storage roots 180 days after sowing. Each bar represents the average ± s.d. of 5 biological replicates containing 4 or 8 storage roots (n = 20 or n = 40). Statistically significant differences are represented by different letters (one-way ANOVA, *P* < 0.05 followed by Scott-Knott test, *P* < 0.05).

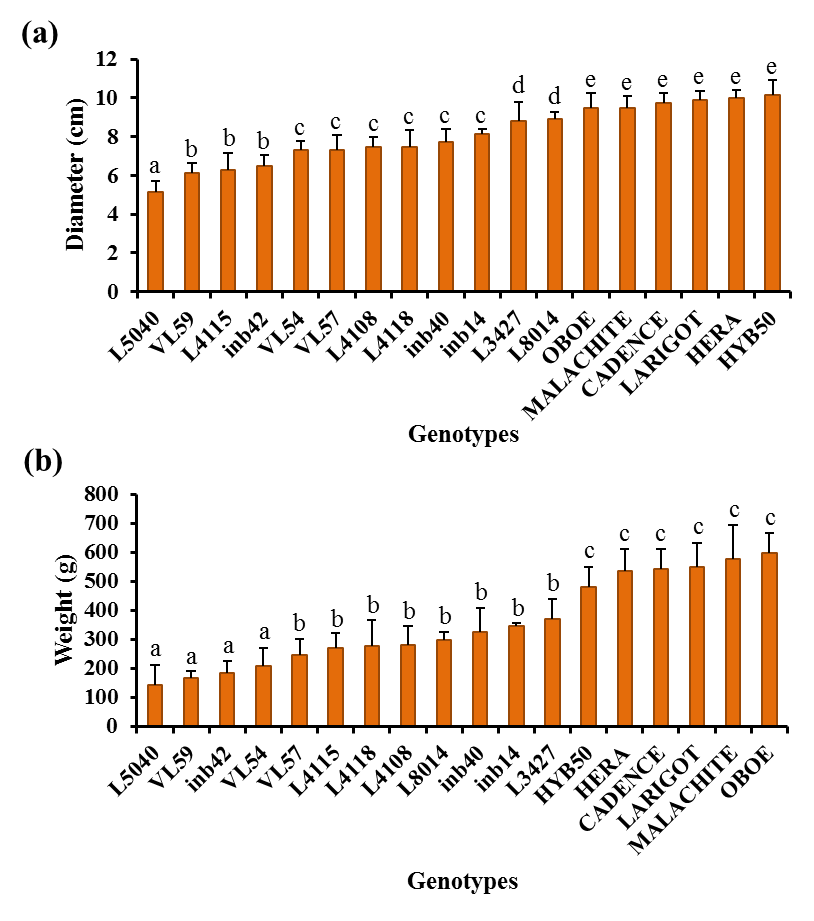

**Fig. S4** Free Asn content in storage roots from 18 genotypes of chicory measured at 30, 60, 90, 120, 150 and 180 days after sowing. At the period of 180 days those genotypes were classified as Low Asn content, Medium Asn content and High Asn content based on the Asn level in the storage roots. Each time point represents mean ± s.d. of 5 biological replicates containing 4 or 8 storage roots (n = 20 or n = 40). The coefficient of determination (R^2^) is shown for the best adjusted model to each genotype.

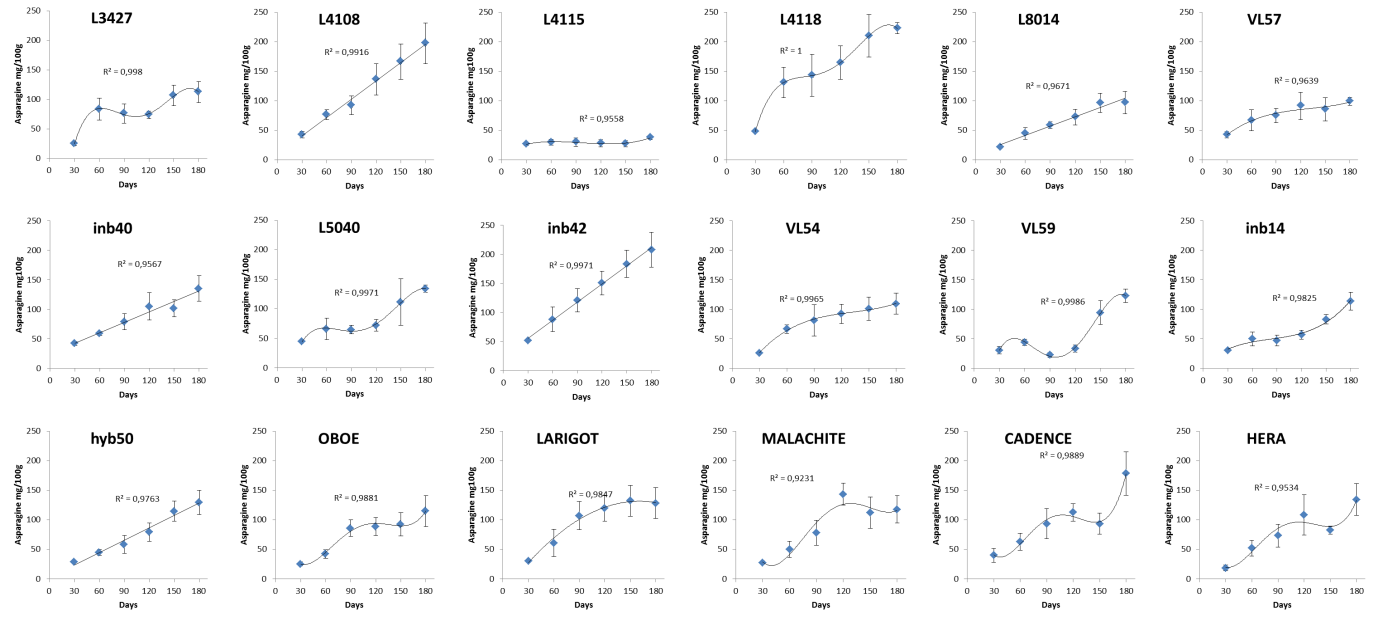

**Fig. S5** Stability evaluation, over 2 years, of the Asn content in 7 genotypes of chicory representative of the three contrasting groups to Asn level in storage roots. Each time point represents mean ± s.d. of 5 biological replicates containing 6 storage roots (n = 30).

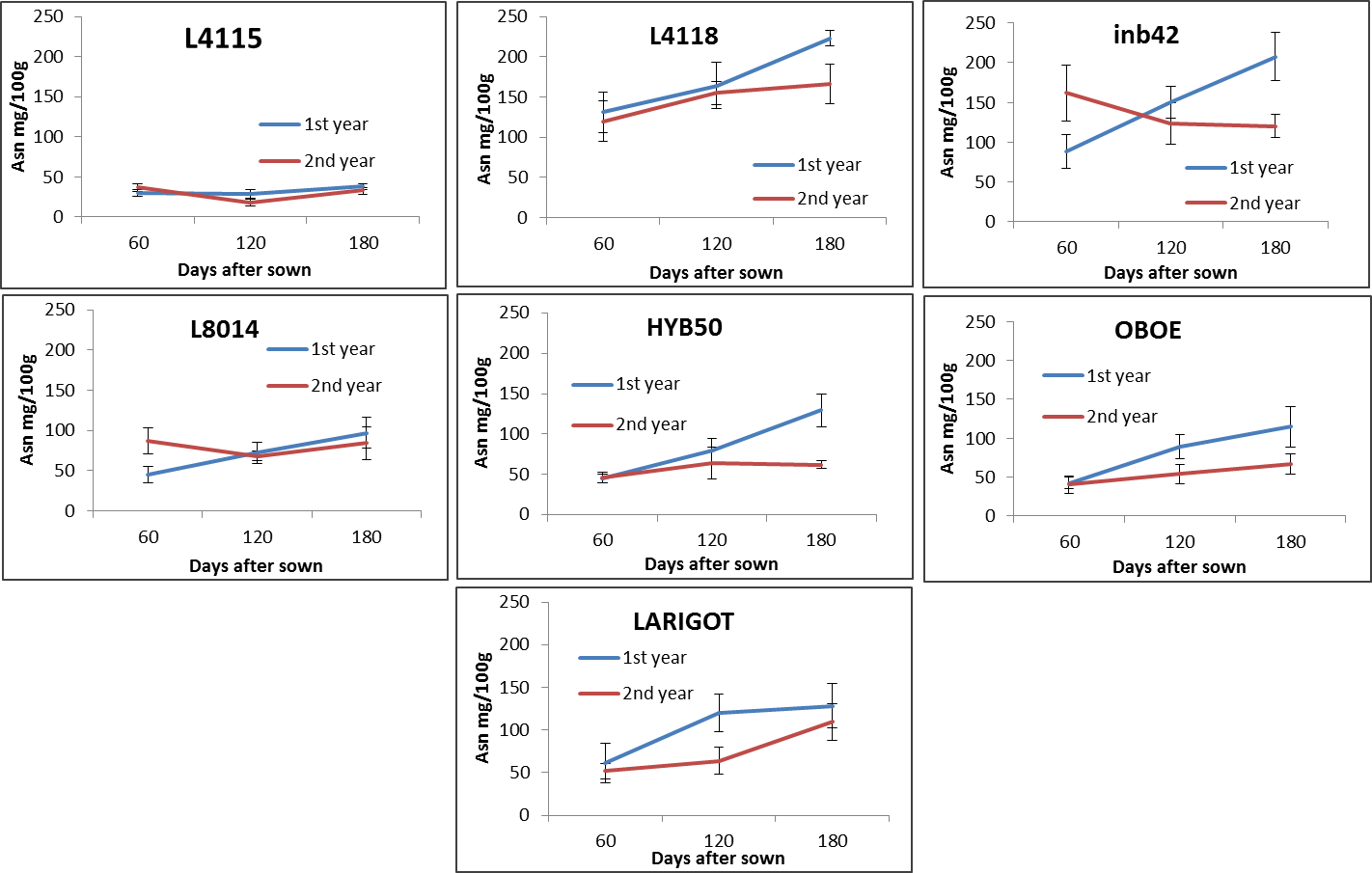

**Fig. S6** Correlation between Asn content (mg/100g) in leaves and storage roots from the genotypes L4115-LAsn, inb40-MAsn, inb42-MHAsn, L4118-HAsn. Each dot represents 1 biological replicate containing samples of 4 or 8 plants. Pearson correlation coefficient (r) was used to measure the linear correlation between the variables X and Y.

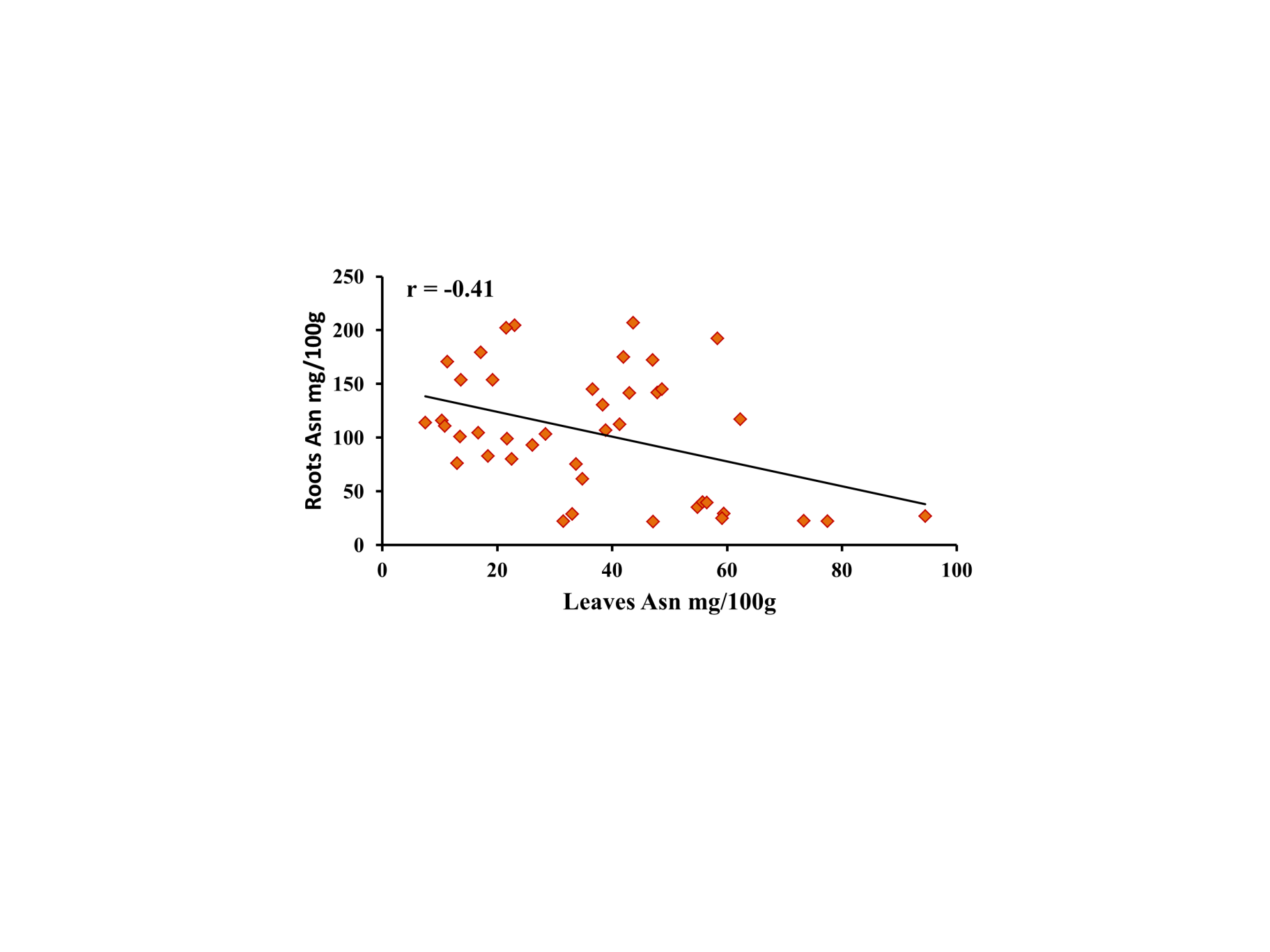

**Fig. S7** Number of genes from the Asn biosynthetic pathway in *A. thaliana* and crop plant species with high acrylamide potential based on the PLAZA 4.0 database ^11^

| Species | Ploidy level | GS | GOGAT | AS | AspAT |
| --- | --- | --- | --- | --- | --- |
| ***Arabidopsis thaliana*** | 2n | 6 | 3 | 3 | 6 |
| ***Cichorium intybus*** | 2n | 4 | 2 | 2 | 4 |
| ***Coffea canephora*** | 2n | 4 | 2 | 4 | 6 |
| ***Solanum tuberosum*** | 4n | 7 | 1 | 2 | 8 |
| ***Triticum aestivum*** | 6n | 12 | 6 | 14 | 15 |

**Fig. S8** Grafting method for chicory (a-f). Because chicory has a very short stem, seeds were germinated in the dark for 5 days to produce elongated stems and to facilitate grafting (a). After germination in the dark, seedlings were transferred to a 16/8 h photoperiod for 25 days to enable stem hardening (b). Leaves from the scions were removed prior to cleft grafting (c). The junction region was covered with a porous tape (d). Plants were harvested 3 months after grafting (e) and storage roots (f) were used for further analysis.

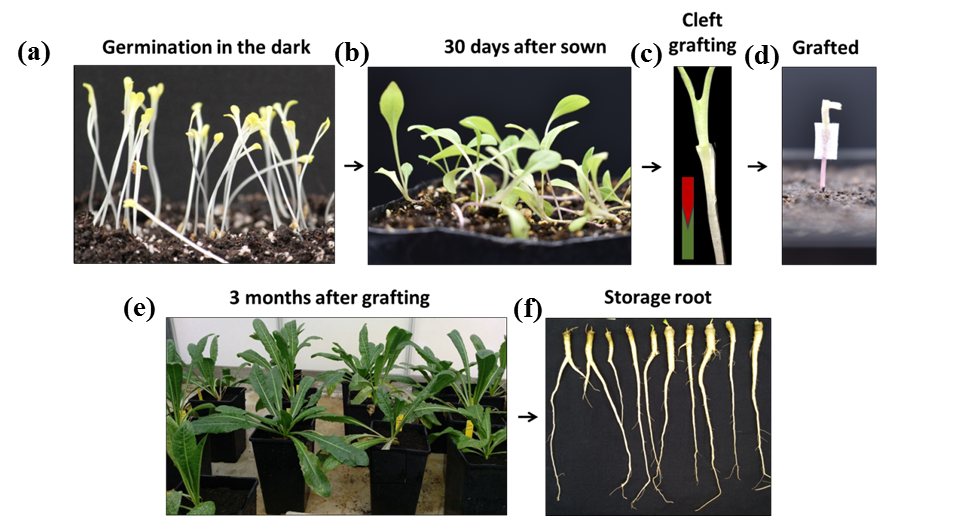

**Fig. S9** Overview of the grafting experiment. Heterografts were done with the most contrasting genotypes for Asn content in storage roots. Homografts of the same genotypes were used as controls. Plants were harvested three months after grafting.

**High asparagine**

**L4118-HAsn**

**Low asparagine**

**L4115-LAsn**

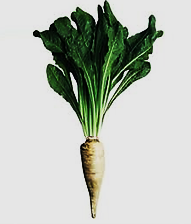

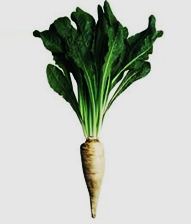

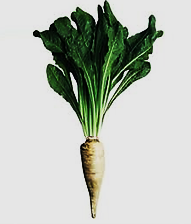

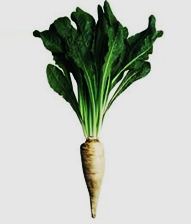

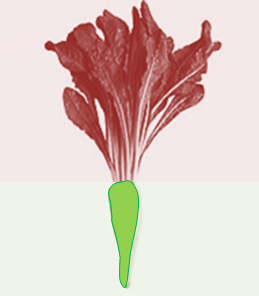

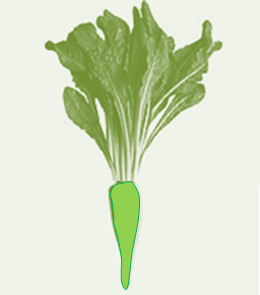

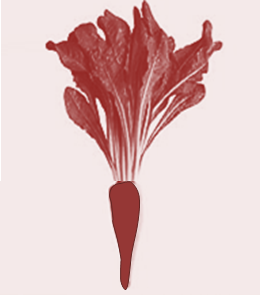

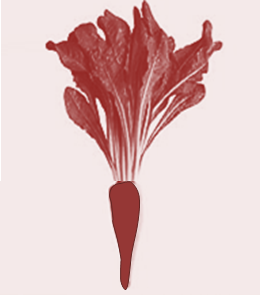

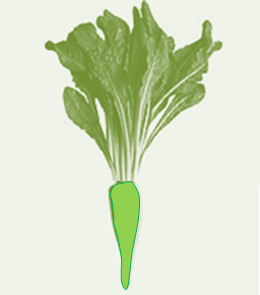

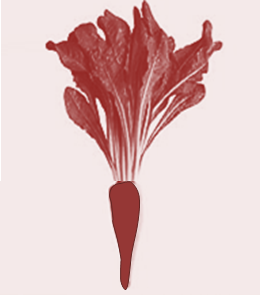

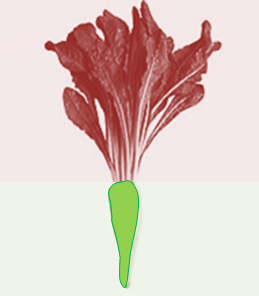

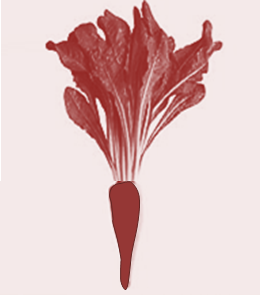

**Phenotype**

**Grafting arrangements**

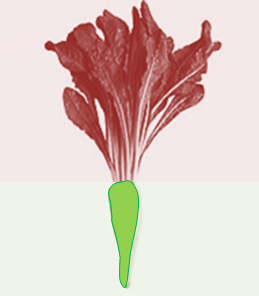

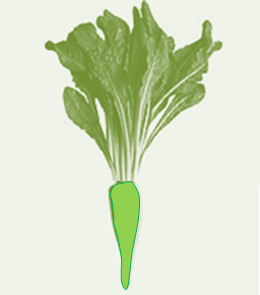

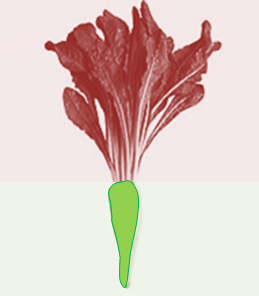

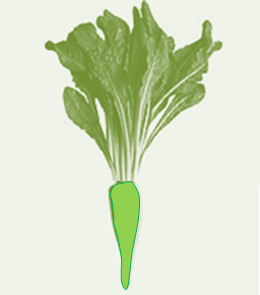

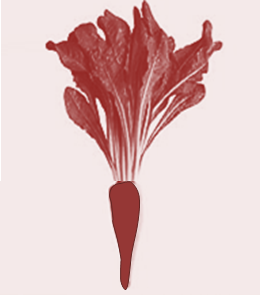

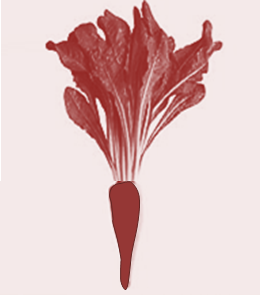

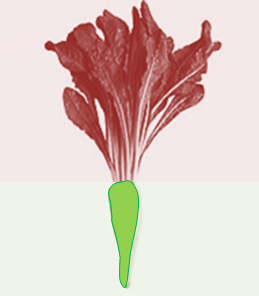

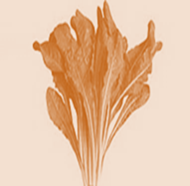

**Results**

**Grafting 1**

**Asn content**

**Low**

**High**

**L4118-HAsn**

**3 months**

**after grafting**

**L4115-LAsn**

**L4115-LAsn**

**L4118-HAsn**

**L4118-HAsn**

**L4115-LAsn**

**L4118-HAsn**

**L4115-LAsn**

**L4118-HAsn**

**L4115-LAsn**

**L4115-LAsn**

**L4118-HAsn**

**L4118-HAsn**

**L4115-LAsn**

**L4118-HAsn**

**L4115-LAsn**

**Self-grafting**

**Self-grafting**

**LAsn/HAsn**

**HAsn/LAsn**

**Grafting 2**

**Grafting 3**

**Grafting 4**

**Fig. S10** Asn feeding experiment. Protoplasts from chicory leaves were placed in 24-well plates with or without Asn and then harvested after 36 h for gene expression analysis. + Asn, protoplast suspensions were supplemented with Asn 190 mM (final concentration); - Asn, no Asn was added; NTC, negative control for the PCR in which cDNA was replaced with water; red arrowheads represent the expected band for *CiASN1*; blue arrowheads represent the expected band for *CiEF* . Three independent experiments were performed.

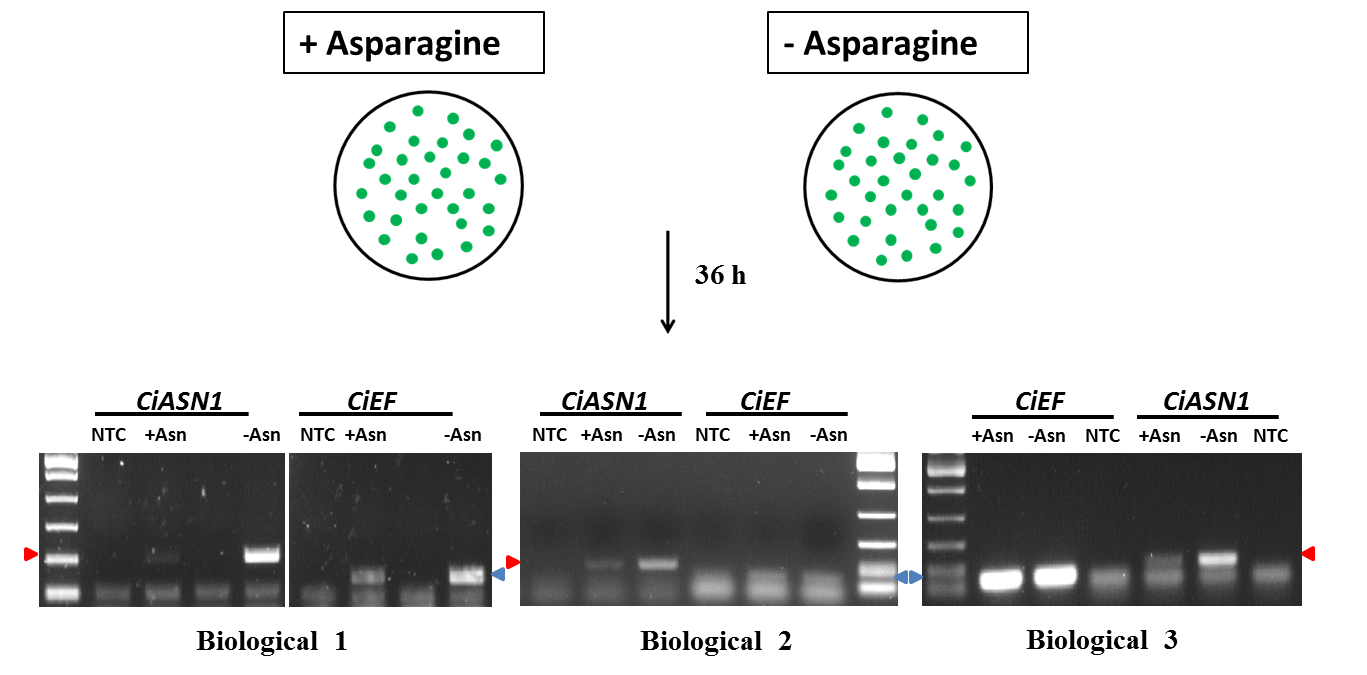

**Fig. S11** Determination of the optimal number of reference genes, according to the pairwise variation V from geNorm, for an accurate normalization in chicory storage roots (a) and leaves (b). The green line represents the cut-off value bellow which we find the optimal number of reference genes to be used for normalization ^17^. For both, storage roots and leaves, the optimal number of reference genes determined was 2.

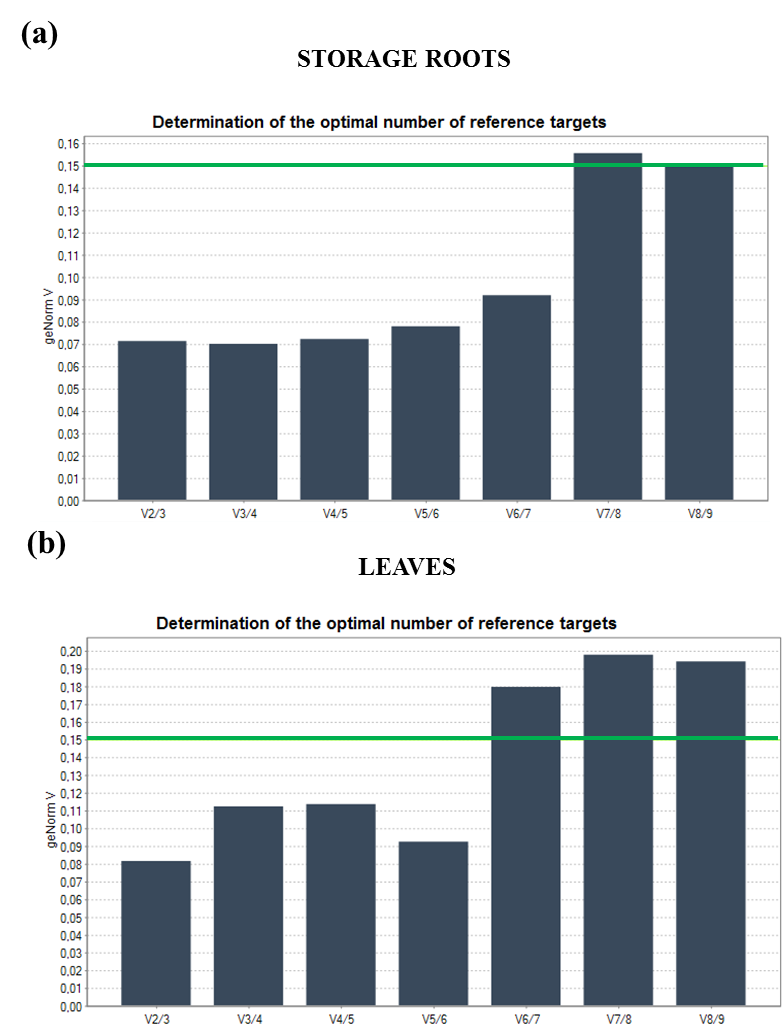

**Fig. S12** Selection of the best reference genes for normalization in storage roots (a) and leaves (b). The reference genes were chosen after ranking in BestKeeper, NormFinder and geNorm. A score based on the sum after ranking with each tool was attributed to each gene in which the lowest score represents the best gene. The genes in green represent the ones chosen for normalization in the respective tissues.

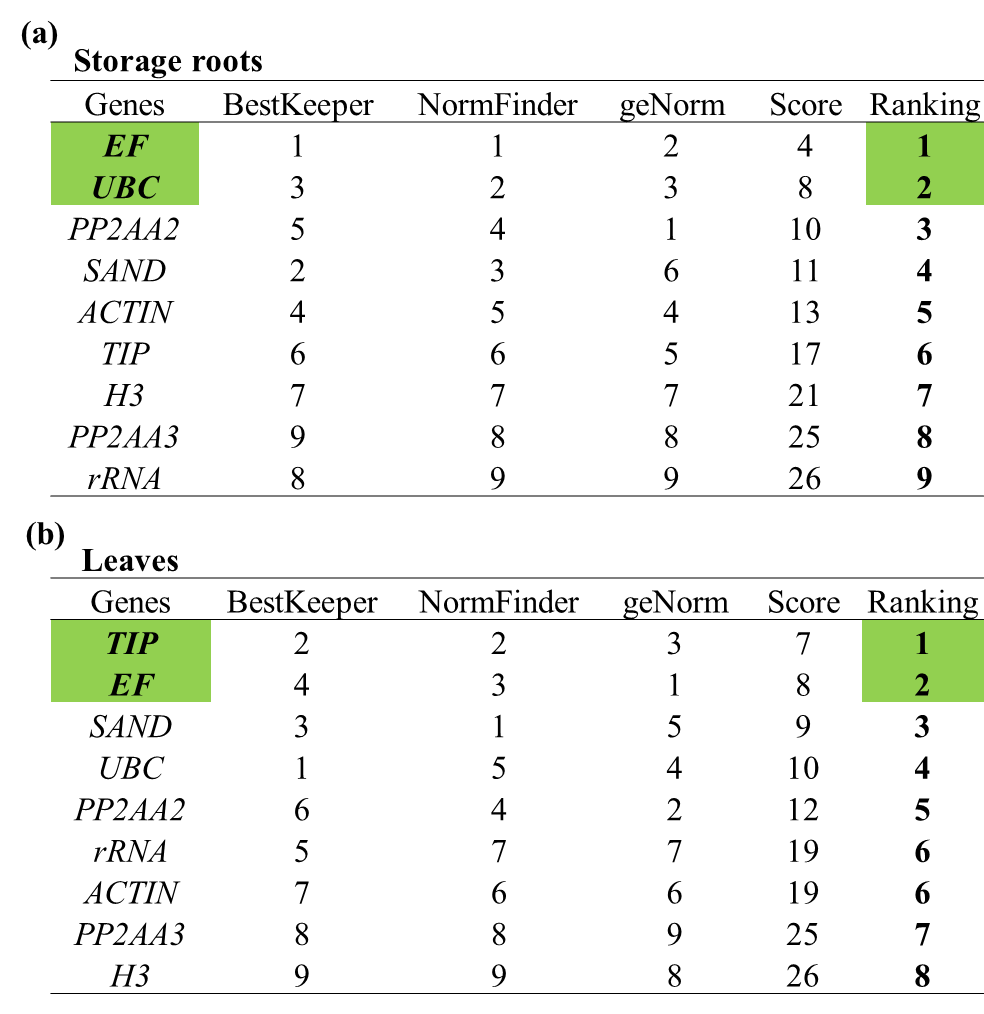
